# Realization of phosphorylation hypothesis of sleep by mammalian CaMKIIβ

**DOI:** 10.1101/2021.10.11.463945

**Authors:** Daisuke Tone, Koji L. Ode, Qianhui Zhang, Hiroshi Fujishima, Rikuhiro G. Yamada, Yoshiki Nagashima, Katsuhiko Matsumoto, Zhiqing Wen, Shota Y. Yoshida, Tomoki T. Mitani, Rei-ichiro Ohno, Maki Ukai-Tadenuma, Junko Yoshida Garçon, Mari Kaneko, Shoi Shi, Hideki Ukai, Kazunari Miyamichi, Takashi Okada, Kenta Sumiyama, Hiroshi Kiyonari, Hiroki R. Ueda

## Abstract

The reduced sleep duration observed in *Camk2a* and *Camk2b* knockout mice revealed the role of Ca^2+^/calmodulin-dependent protein kinase II (CaMKII)α/CAMKIIβ as sleep-promoting kinases and lead to the phosphorylation hypothesis of sleep. However, the underlying mechanism of sleep regulation by kinases and protein phosphorylation is largely unknown. Here, we demonstrate that the phosphorylation states of CaMKIIβ regulates sleep duration and sleep needs. Importantly, the activation or inhibition of CaMKIIβ can increase or decrease sleep duration by almost two-fold, supporting the role of CaMKIIβ as a core sleep regulator in mammals. This sleep regulation depends on the kinase activity of CaMKIIβ in excitatory neurons. Furthermore, CaMKIIβ mutants mimicking different phosphorylation states can regulate various sleep steps including sleep induction, sleep maintenance, and sleep cancelation. Key CaMKIIβ residues responsible for the mode switch undergo ordered (auto-)phosphorylation. We thus propose that ordered multi-site phosphorylation of CaMKIIβ underlies multi-step sleep regulation in mammals.

## INTRODUCTION

A wide range of biological phenomena, including organism-level behaviors, rely on the regulation of protein activity by phosphorylation. The circadian clock is an excellent example of the marked role of protein phosphorylation in the regulation of an organism-level behavior ^1–3^. Genetic screening of animal behavior revealed that the *period* (*per*) gene is a core factor for the circadian clocks ^4^. Casein kinase I (CKI) phosphorylates the PER protein, and a human lineage showing abnormalities in circadian behavioral rhythms had a single amino acid substitution at the phosphorylation residue ^5^. The phosphorylation of PER by CKI is considered a major regulator of circadian period length for the following reasons: first, the targeted mutation of a single phosphorylation residue in PER can bidirectionally change the period length of the circadian clock ^6, 7^. Second, the effect is significant, with changes in CKI-kinase activity resulting in a more than two-fold change in period length, at least in culture cells ^8^.

The sleep-wake cycle, like the circadian clocks, is a physiological function that governs the organism-level behavioral rhythms and is believed to regulate synaptic function ^9^. However, the molecular mechanisms regulating the daily amount of sleep and the transitions between sleep and wake phases are not fully understood. Genetic screening studies have revealed that protein kinases play an important role in sleep duration regulation. In particular, knocking out the first sleep-promoting kinases discovered, *Camk2a* and *Camk2b,* markedly reduced sleep duration in mice ^10^. Subsequent phosphoproteomics studies have shown that the phosphorylation states of neuronal proteins vary with the sleep-wake cycle and in response to sleep deprivation ^11–13^. The phosphoproteomics profile revealed an alteration of the phosphorylation states of Ca^2+^/calmodulin-dependent protein kinase II (CaMKII)α/CAMKIIβ and its potential substrates (e.g., Synapsin 1). These results suggest that CaMKIIα/CaMKIIβ plays an important role in mammalian sleep regulation and support the phosphorylation hypothesis of sleep (the idea that sleep is regulated by protein phosphorylation).

The phosphorylation hypothesis of sleep ^10, 14^ assumes that the neural activity associated with wakefulness acts as an *input* to activate sleep-promoting kinases such as CaMKIIα/CaMKIIβ ^10^, SIK1/SIK2/SIK3 ^15, 16^, and ERK1/ERK2 ^17^. Another prediction is that sleep-promoting kinases may need to *store* some form of information associated with wakefulness. This is because awakening does not immediately lead to sleep, but rather *stores* a history of awakening as a sleep need. As an *output* of sleep regulation, sleep-promoting kinases might induce sleep by phosphorylating their substrates. CaMKIIα/CaMKIIβ has unique features that might make this kinase suitable for achieving the *input*, *storage*, and *output* mechanism of sleep regulation. A well-known mechanism of CaMKIIα/CaMKIIβ activation is the intracellular Ca^2+^ influx that occurs upon excitatory synaptic input and subsequent neuronal firing ^18, 19^. Intracellular Ca^2+^ binds to calmodulin (CaM), which binds to CaMKIIα/CaMKIIβ and switches its kinase domain to the exposed open and kinase-active form. The kinase-active CaMKIIα/CaMKIIβ undergoes autophosphorylation along with phosphorylation of other substrate proteins. T286 (CaMKIIα) and T287 (CaMKIIβ) are the first residues undergoing autophosphorylation upon activation of CaMKIIα/CaMKIIβ. T286 and T287 phosphorylation switches CaMKIIα/CaMKIIβ to its kinase-active form even in the absence of Ca^2+^/CaM ^20–22^. The maintained kinase activity due to T286 and T287 phosphorylation is called autonomous activity. Finally, the activated CaMKIIα/CaMKIIβ phosphorylates several neuronal proteins. The sequential autoregulation of CaMKII activity serves as a neuronal timer in a minutes time scale in fruits fly ^23^. However, the effect and mechanism of CaMKIIα/CaMKIIβ on sleep regulation and duration in mammals have not been rigorously investigated.

Furthermore, the dynamics of sleep-wake are not only characterized by the duration of sleep, but also by the distribution of sleep and wake episodes. Indeed, *Camk2a* and *Camk2b* knockout mice are less likely to transition from wake to sleep and from sleep to wake ^10^. This suggests that CaMKIIα/CaMKIIβ elicits the transition between wake and sleep. It should be noted that sleep duration and sleep-wake transition can be independently regulated: for example, knocking out *orexin* barely affects sleep duration, but significantly increases the sleep-wake transition ^24, 25^. Given the physiological process of the sleep-wake cycle, it is reasonable to assume that organisms employ multiple and stepwise mechanisms to regulate sleep. It would begin with sleep induction and switch to sleep maintenance. CaMKIIα/CaMKIIβ itself undergoes multiple and stepwise changes (multi-site autophosphorylation, dodecameric oligomerization, and conformational changes) ^18, 26^. Following the phosphorylation of T286 and T287, the activated kinase catalyzes the autophosphorylation of residues such as T305 and T306 (CaMKIIα), and T306 and T307 (CaMKIIβ). Phosphorylation of these residues inhibits the binding of Ca^2+^/CaM to CaMKIIα/CaMKIIβ ^27–29^. The autoregulatory mechanism of CaMKIIα/CaMKIIβ may be more complex than a two-step regulation. It was reported that autophosphorylation can occur multiple residues other than well-understood T286/T305/T306 (CaMKIIα) and T287/T306/T307 (CaMKIIβ) with different efficiency depending on residues ^30^ and the dodecameric CaMKIIα/CaMKIIβ structure may have many intermediate states ^31^. Although the sleep-wake cycle affects the level of such multi-site autophosphorylation of CaMKIIα/CaMKIIβ ^11–13, 32, 33^, little is known about the actual function of the multi-site autophosphorylation in the regulation of the sleep-wake cycle. Of the four *Camk2* homologs (i.e., *Camk2a*, *Camk2b*, *Camk2d* and *Camk2g*), knockout mice of *Camk2b* showed the most pronounced decrease in sleep duration per day ^10^. Thus, this study will focus on CaMKIIβ and aims to comprehensively analyze the sleep phenotype caused by a series of CaMKIIβ mutants mimicking the different phosphorylation states.

## RESULTS

### Phosphorylation of CaMKIIβ regulates sleep induction

To investigate whether CaMKIIβ regulates sleep depending on the phosphorylation state of CaMKIIβ, we conducted an *in vivo* comprehensive phosphomimetic screening of CaMKIIβ. Mouse CaMKIIβ protein has 69 serine (S) and threonine (T) residues that can be the target of autophosphorylation (**Figure 1a**). We assessed the contribution of these residues to sleep regulation by expressing a series of phosphomimetic mutants of CaMKIIβ, in which aspartic acid (D) replaced one of the phosphorylable residues. Each of the 69 CaMKIIβ mutants was expressed under the control of human *synapsin-1* (*hSyn1*) promoter and delivered in wild-type mice brain by an adeno-associated virus (AAV) system AAV-PHP.eB ^34^, which allows broad gene expression throughout the brain (**Figure 1b**). The whole-brain expression of H2B-mCherry reporter under the *hSyn1* promoter delivered by the AAV system was confirmed by whole-brain imaging using the CUBIC method (**Figure 1c** and **Figure 1-figure supplement 1a**). Unless otherwise indicated, we refer to mice with AAV-mediated expression of CaMKIIβ mutants simply by the mutant name (e.g., T287D mice). We measured the sleep parameters of the mice expressing mutant CaMKIIβ using a respiration-based sleep phenotyping system, snappy sleep stager (SSS) ^24^ (**Figure 1b**). Sleep measurements were started at 8 weeks old following the AAV administration at 6 weeks old. Mice expressing AAV-induced wild-type (WT) CaMKIIβ and untreated mice had similar daily sleep durations (733.9 ± 6.1 and 724.7 ± 4.3 min ^24^, respectively; all mice phenotypes are reported as mean ± SEM). In this screening, mice expressing T287D, S114D or S109D CaMKIIβ mutants had top three extended daily sleep duration (846.7 ± 23.7, 839.7 ± 14.1 or 803.4 ± 16.2 min, respectively), though the phenotype of S109D showed no statistical significance (**Figure 1d**). Although no statistical significance was obtained for sleep transition parameters *P_WS_* (probability of transition from wakefulness to sleep) and *P_SW_* (probability of transition from sleep to wakefulness) in this first screening (**Figure 1-figure supplement 2a, b**), the *P_WS_* of T287D mice was higher than that of WT-expressing mice, which is opposite to the phenotype of *Camk2b* knockout mice ^10^. There was no correlation between the ensemble of sleep duration and AAV transduction efficiency among the analyzed mutants (**Figure 1-figure supplement 2c)**, indicating that the observed sleep phenotypes can be attributed to the nature of the introduced mutations rather than to a possible difference in AAV transduction efficacy.

**Figure 1.**
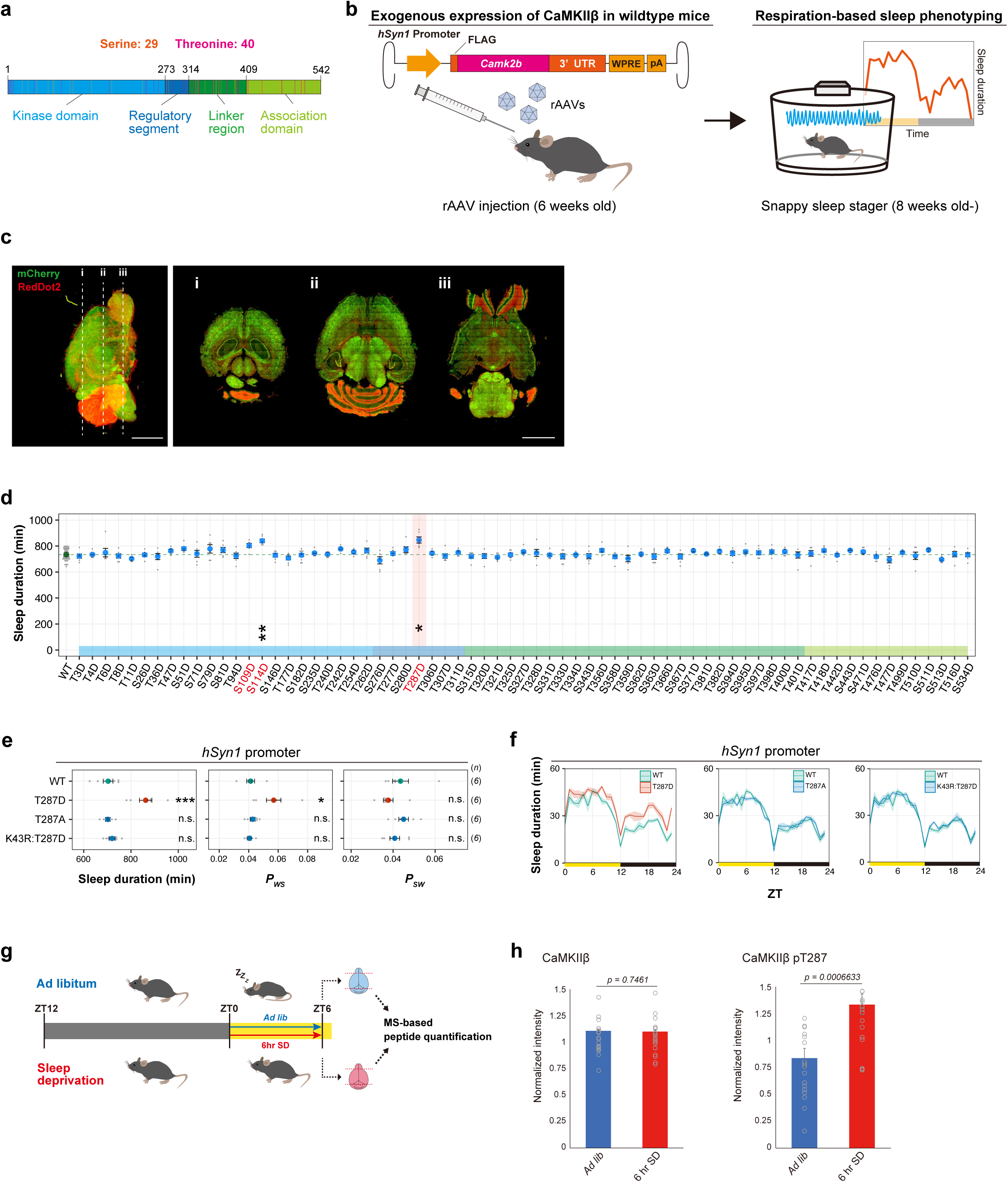
Phosphorylation of CaMKIIβ regulates sleep induction. **(a)** The 29 serine and 40 threonine residues throughout CaMKIIβ. Orange and magenta lines represent serine and threonine residues, respectively. Color-coded regions indicate the functional domains of CaMKIIβ. **(b)** Schematic diagram of AAV-based CaMKIIβ expression and respiration-based sleep phenotyping. UTR: untranslated region. pA: polyA. **(c)** Representative cross-sectional images of the brain of mice expressing H2B-mCherry under the *hSyn1* promoter by the AAV. Data were acquired by whole brain imaging with RedDot2 counterstaining, and detailed images of each brain region are shown in **Figure 1-figure supplement 1a**. Scale bars, 5 mm. **(d)** Daily sleep duration of mice expressing CaMKIIβ phosphomimetic mutants (n = 6–10) in the presence of endogenous wildtype CaMKIIβ. The represented value is the average of SSS measurements over six days. The dashed green line represents the average sleep duration of wild-type CaMKIIβ-expressing mice (WT, n = 48). Multiple comparison test was performed against WT. **(e-f)** Sleep/wake parameters **(e)** and sleep profiles **(f)** of mice expressing T287-related CaMKIIβ mutants, averaged over six days. Measurements are independent of those in **(d)**. Sleep duration is the total sleep duration in a day, *P_WS_* and *P_SW_* are the transition probabilities between wakefulness and sleep. The shaded area represents the SEM. Multiple comparison test was performed against WT. ZT: zeitgeber time. **(g)** Sleep deprivation and peptide quantification procedures. The brains of the sleep-deprived and control mice were collected for MS-based peptide quantification. **(h)** Total CaMKIIβ and T287-phosphorylated peptides from brains of sleep-deprived and control mice, analyzed by SRM quantitative mass spectrometry. Error bars: SEM. *p < 0.05, **p < 0.01, ***p < 0.001, n.s.: no significance. **Figure 1-source data 1** Source data for Figure 1d, e, f. **Figure 1-source data 2 and 3** Source data for Figure 1h.

To confirm the reproducibility of the extended sleep duration for T287D, S114D and S109D mice, we conducted an independent set of experiments. These confirmed the prolonged sleep duration of T287D mice (861.9 ± 26.1 min) and the increase in *P_WS_* (**Figure 1e**). The extended sleep duration of T287D mice does not depend on the circadian timing because the mice showed increased sleep duration at most zeitgeber time of the day (**Figure 1f**). Besides, this second round of evaluation did not show a significant increase in the sleep duration of S114D mice and S109D, although a trend of extended sleep duration was observed for S109D mutant (**Figure 1-figure supplement 2d and 2e**). We concluded that T287D CaMKIIβ is the mutant that robustly increased sleep duration *in vivo*.

Replacing T287 with the non-phosphomimetic alanine (A) did not extend sleep duration (701.5 ± 9.8 min) (**Figure 1e, f**). This supports that the phosphorylation-mimicking property of D caused the sleep duration extension. Furthermore, the extended sleep duration depends on the kinase activity of CaMKIIβ, because the kinase-dead (K43R) version of the T287D mutant (i.e., K43R:T287D) did not extend sleep duration (719.8 ± 12.4 min). Given that the phosphorylation of T287 inhibits the interaction between the kinase domain and the regulatory segment of CaMKIIβ (which leads to the open and kinase-active conformation of the kinase), the normal sleep duration of K43R:T287D mice suggests that CaMKIIβ with open conformation alone is insufficient to lengthen sleep duration. We thus propose that CaMKIIβ induces sleep via T287 phosphorylation and that this process requires the kinase activity of CaMKIIβ.

The robust sleep induction by the T287D mutant suggests that T287 phosphorylation marks the level of sleep need. This has been supported through previous studies; for example, the level of CaMKIIα T286 phosphorylation or CaMKIIβ T287 phosphorylation follows the expected level of sleep need upon six hours sleep deprivation and subsequent recovery sleep analyzed by western blotting ^12^. Moreover, the level of CaMKIIα T286 phosphorylation follows the expected sleep need along with normal sleep wake cycle: a previous study showed the circadian rhythmicity of CaMKIIα T286 phosphorylation peaking at the end of the dark (wake) phase and decreasing throughout the light (sleep) phase ^13^. Consistent with this rhythmicity, another study indicated CaMKIIα T286 phosphorylation is higher at the dark (wake) phase ^11^. Because several studies focus on CaMKIIα and rely on western blotting technique, we also examined whether the phosphorylation levels of T287 in the brain increased upon six hours sleep deprivation by using a quantitative and targeted selected-reaction-monitoring (SRM) analysis. The SRM analysis confirmed that sleep deprivation increased T287 phosphorylation of endogenous CaMKIIβ without changing the amount of total CaMKIIβ (**Figure 1g** and **1h**). In addition, the phosphorylation level of CaMKIIα T286 and CaMKIIβ T287 correlated well, suggesting that these phosphorylation levels similarly respond to sleep deprivation (**Figure 1-figure supplement 2f, g**).

### Biochemical evaluation of sleep-inducing CaMKIIβ mutants

To compare the kinase activity and mice sleep phenotypes, we measured the kinase activity of each mutant *in vitro* using cell lysate system. We prepared cell extracts of 293T cells overexpressing the CaMKIIβ mutants. Relative expression level was quantified for each mutant by dot blot (**Figure 1-figure supplement 3a**). The relative amounts of CaMKIIβ as well as cellular components derived from the extracts were adjusted by mixing CaMKIIβ-expressing 293T lysate and mock-transfected 293T lysate. This adjustment process was not applied for the mutants having <25% expression level compared with wild-type CaMKIIβ. Then, the enzymatic activity of the expressed CaMKIIβ in the presence and absence of CaM (**Figure 1-figure supplement 3b**).

Most mutants as well as WT exhibit kinase activity only in the presence of CaM (**Figure 1-figure supplement 3b**). S109D, T242D, and T287D mutants showed marked enzyme activity even in the absence of CaM. The CaM-independent kinase activity of T287D is consistent with the constitutive kinase-active property of T287D. However, the kinase activity of T287D in the presence of CaM is lower than that of WT. By contrast, S109D and T242D showed no reduction in the kinase activity in the presence of CaM and the CaM-independent kinase activity is higher than that of T287D. The reason of this lower T287D activity is currently unknow but might be, at least in part, due to the inhibitory autophosphorylation that was underway in the 293T cell during the period between the expression of the T287D protein and the preparation of the cell lysate, and structural thermal-instability elicited by the detachment of regulatory segment from the kinase domain ^35^. Since these inhibitory mechanisms are caused by the constitutive-kinase activation (and/or structural alteration from close to open conformation) of the enzyme, the final kinase activity will appear as the sum of positive and negative factors: therefore, it is important to be careful in discussing the relationship between whether a mutation activates or inhibits kinase activity based on the one-point relative strength of the phosphorylation activity alone.

Although there are limitations in the biochemical evaluation of kinase activity in this cell lysate system as described above, it appears reasonable to assume that mutations, in which CaM-independent activity is detected, have at least the property of showing CaM-independent phosphorylation activity, unlike the wild-type enzyme. Similar to the kinase-dead mutation K43R, the mutation that reduces the phosphorylation activity to a level similar to that of the background from cell extracts may also be regarded as a reliable phenotype, basically acting in a repressive manner on the kinase activity. Given that the level of AAV-mediated CaMKIIβ expression is much lower than the level of endogenous CaMKIIβ (**Figure 1-figure supplement 1b**), it would be reasonable to assume that the CaMKIIβ mutants showing CaM-independent activity affected sleep by exhibiting a dominant phenotype (e.g., T287D and S109D), even in the presence of abundant endogenous CaMKIIα/CaMKIIβ protein. It is also quite possible that this sleep phenotype is mediated by the activation of the endogenous CaMKIIα/CaMKIIβ by the constitutive-active mutant. Also, the mutation with reduced kinase activity may not have had a dominant negative effect on sleep in the presence of higher level of the endogenous CaMKIIβ due to its low expression level mediated by AAV vector, and thus did not show a pronounced phenotype in the current screening.

It should be noted that while T287D had high kinase activity in the absence of CaM, T242D and especially S109D showed even higher kinase activity in the absence of CaM. However at least T242D appears not to extend sleep duration and the effect of S109D on the sleep duration is milder than that of T287D *in vivo*. Hence, the results of the present kinase assay using a conventional peptide substrate do not fully account for the quantitative level of sleep induction observed *in vivo*, suggesting the existence of an additional layer of regulation.

Among the kinase-inactive mutants and others, several mutants had significantly reduced expression levels (e.g., S182) (**Figure 1-figure supplement 3a**). Reduced protein expression levels and/or protein stability inherent in such mutants could also be a reason why these mutants do not exhibit a dominant active sleep-promoting activity in the screening *in vivo*. The unstable sleep phenotype of S114D might be related to the unstable/low-expression nature of this mutant at least in culture cell—as with the kinase activity evaluation, protein expression levels in the mouse brain do not always correlate with expression levels in 293T cells, and should be considered carefully though.

### Phosphorylation of CaMKIIβ regulates NREM sleep induction and sleep needs

To further investigate the role of T287 phosphorylation in sleep regulation, we expressed the CaMKIIβ T287-related mutants under the *Camk2a* promoter ^36^, which is a well-characterized promoter inducing gene expression preferentially to the excitatory neurons. As **Figures 1e and f** show, the daily sleep duration of T287D mice was higher than that of WT-expressing mice, which is consistent with the results obtained with the *hSyn1* promoter. WT, T287A, K43R:T287D, and PBS-administrated mice had comparable sleep phenotypes (**Figures 2a, b**). As observed in T287D mice with the *hSyn1* promoter, T287D mice with the *Camk2a* promoter had a significantly higher *P_WS_* (**Figure 2a**), suggesting that the T287-phosphorylated CaMKIIβ promotes the transition from wakefulness to sleep. We also reproduced the increased sleep duration by expressing the T287D mutant under the *Camk2b* promoter cloned in this study (**Figure 2-figure supplement 1a, b**).

**Figure 2.**
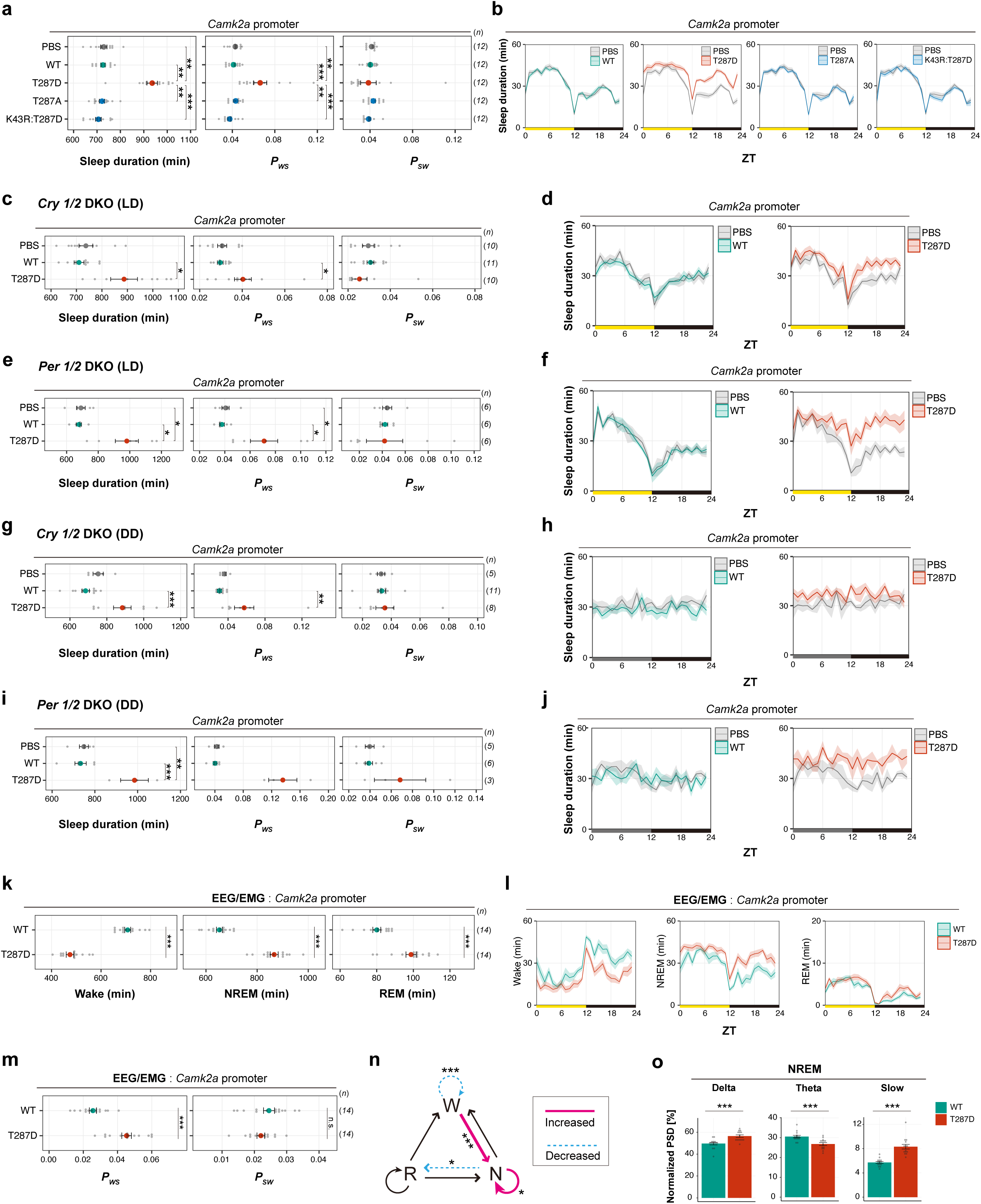
Phosphorylation of CaMKIIβ regulates NREM sleep induction and sleep needs. **(a-b)** Sleep/wake parameters **(a)** and sleep profiles **(b)** of mice expressing CaMKIIβ T287-related mutants under the *Camk2a* promoter, averaged over six days. Shaded areas represent SEM. Multiple comparison test was performed against PBS-injected control mice (PBS). **(c-f)** Sleep/wake parameters and sleep profiles, averaged over four days, of *Cry1/2* DKO mice **(c and d)** and *Per1/2* DKO mice **(e and f)** expressing wild-type CaMKIIβ (WT) or the T287D mutant under the light/dark condition. Multiple comparison tests were performed between all individual groups. **(g-j)** Sleep/wake parameters, averaged over four days, of *Cry1/2* DKO mice **(g and h)** and *Per1/2* DKO mice **(i and j)** expressing wild-type CaMKIIβ (WT) or the T287D mutant under constant dark. Multiple comparison tests were performed between all individual groups. **(k-m)** Sleep phenotypes **(k and m)** and sleep profiles **(l)** measured by EEG/EMG recordings for mice expressing CaMKIIβ (WT) or the T287D mutant. **(n)** Differences in transition probabilities (between wakefulness (W), NREM sleep (N), and REM sleep (R)) between mice expressing WT CaMKIIβ or the T287D mutant. Magenta lines and dashed blue lines indicate when the values for the T287D-expressing mice are significantly (p < 0.05) higher and lower, respectively. **(o)** NREM power density in typical frequency domains of mice expressing WT CaMKIIβ or the T287D mutant. Error bars: SEM, *p < 0.05, **p < 0.01, ***p < 0.001, n.s.: no significance. **Figure 2-source data 1** Source data for Figure 2a-o

*Camk2* plays a role in the regulation of the circadian rhythm ^37^. To examine whether the sleep-inducing effect of the CaMKIIβ T287D mutant depends on the behavioral circadian rhythmicity, we expressed it in *Cry1*^-/-^:*Cry2*^-/-^ and *Per1*^-/-^:*Per2*^-/-^ double knockout mice (*Cry1/2* DKO and *Per1/2* DKO) using the *Camk2a* promoter. Both DKO mice lines are deficient in behavioral circadian rhythmicity in constant dark (DD) ^38–41^. Under light/dark (LD) conditions, the daily sleep duration of T287D-expressing *Cry1/2* DKO and *Per1/2* DKO mice was significantly higher than that of WT CaMKIIβ-expressing mice (**Figure 2c, d, e, f**). Under constant dark, where both DKO mice lack a clear circadian behavioral rhythmicity, the sleep duration of T287D-expressing mice increased irrespective of circadian time across the 24 h (**Figure 2g, h, i, j**). This increased sleep duration under constant dark is associated with increased *P*_WS_. These results demonstrate that the sleep-inducing effect of the T287D mutant is independent of behavioral circadian rhythmicity and canonical core clock genes such as *Cry1/Cry2* or *Per1/Per2*.

The sleep-inducing effects of the T287D mutant could be attributed to an impairment in the proper maintenance of wakefulness. To examine whether the arousal system in T287D mice is normal, we assessed their responses to external stimuli. The novel cage environment promotes awakening by stimulating the mice’s exploratory behavior ^42^. Cage exchange significantly decreased the sleep duration of T287D, WT, and PBS-administrated mice compared with the baseline duration (**Figure 2-figure supplement 1c**), suggesting that the sleep-extending effect of the T287D mutant is not due to abnormalities in the arousal system.

Since the sleep-inducing effect of the T287D mutant depends neither on the circadian rhythms nor on an abnormal arousal system, it might directly alter sleep needs, which can be estimated through the delta-wave of an electroencephalogram (EEG). We recorded EEGs and electromyograms (EMG) of the mice expressing the CaMKIIβ T287D mutant under the *Camk2a* promoter. The EEG/EMG recordings revealed that T287D mice had significantly higher daily non-rapid eye movement (NREM) and REM sleep duration (**Figure 2k, l**) and *P*_WS_ (**Figure 2m**) than WT-expressing mice. This data is consistent with the SSS measurements (**Figure 2a**). The analysis of transition probabilities between wake, NREM, and REM episodes revealed a large decrease (p < 0.001) in wake maintenance (W to W) and increase (p < 0.001) in the transitions from wake to NREM (W to N) compared with WT-expressing mice (**Figure 2n**). These results suggest that the T287 phosphorylation of CaMKIIβ induces sleep by increasing wake to NREM transitions. Besides, we confirmed that T287D mice had significantly higher delta power and slow power during sleep episodes (**Figure 2o** and **Figure 2-figure supplement 1d**), suggesting elevated sleep needs. We obtained similar EEG/EMG recordings with mice expressing T287D mutant under the *hSynI* promoter (**Figure 2-figure supplement 1e-j**). These results demonstrate that T287-phosphorylated CaMKIIβ provokes physiological sleep needs and acts on the transition from wake to NREM sleep.

### Phosphorylation of CaMKIIβ in excitatory neurons regulates sleep induction

A potential limitation of the use of *Camk2a* promoter is that the expression is highly enriched in excitatory neurons but not exclusively localized ^36^. We then investigated the neuronal cell types responsible for the CaMKIIβ-mediated sleep induction by using other strategy using AAVs carrying double-floxed inverted open reading frame (DIO) constructs and mouse lines expressing *Cre* recombinases in specific neurons (*Cre*-mice) (**Figure 3a**). CaMKIIβ T287D expression in *Vglut2-*specific neurons significantly increased sleep duration compared to the WT CaMKIIβ-expressing mice (**Figure 3b, c**), while expression of the T287D mutant in *Gad2*-specific neurons did not affect sleep phenotype (**Figure 3d, e**). These results confirm that glutamatergic excitatory neurons are involved in the sleep promotion by the CaMKIIβ T287D mutant.

**Figure 3.**
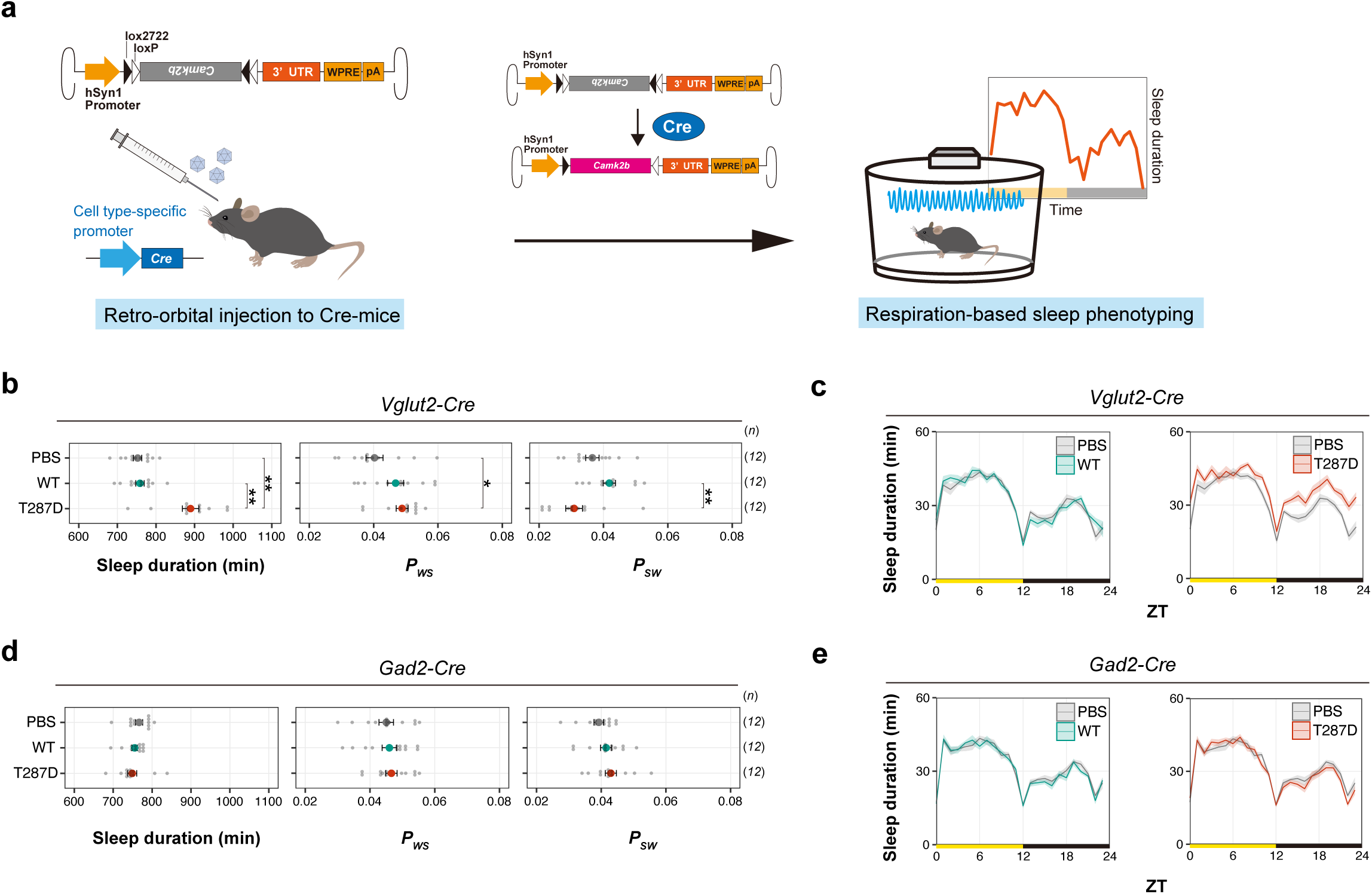
Phosphorylation of CaMKIIβ in excitatory neurons regulates sleep induction. **(a)** Schematic diagram of cell type-specific expression of CaMKIIβ using AAV and Cre-mice. Cre-mediated recombination of AAV genomes results in *Camk2b* gene expression in the target cells. **(b-c)** Sleep/wake parameters **(b)** and sleep profiles **(c)** of *Vglut2-Cre*-mice administrated with AAV-DIO-*Camk2b*, averaged over six days. Shaded areas represent SEM. Multiple comparison tests were performed between all individual groups. **(d-e)** Sleep/wake parameters **(d)** and sleep profiles **(e)** of *Gad2-Cre*-mice administrated with AAV-DIO-*Camk2b*, averaged over six days. Multiple comparison tests were performed between all individual groups. Error bars: SEM, *p < 0.05, **p < 0.01, ***p < 0.001, n.s.: no significance. **Figure 3-source data 1** Source data for Figure 3b-e

### Kinase activity of CaMKIIβ bidirectionally regulates sleep

Having the different efficacy of sleep-inducing activity among the biochemical constative-active CaMKIIβ mutants (e.g., T287D and S109D), we next sought to confirm the relationship between the CaM-independent enzymatic activity of CaMKIIβ and sleep promotion by using another type of constitutive-active CaMKIIβ. To this end, we used CaMKIIβ deletion mutant that lacks the C-terminal half involving the regulatory segment, linker region, and oligomerization domain ^43^). The CaMKIIβ deletion mutant is constitutively active due to the exposed kinase domain but does not retain T287 and subsequent residues (**Figure 4a**). Similar to T287D mice, mice expressing the deletion mutant (del) showed an extended sleep duration and increased *P_WS_*. The extended sleep duration depends on the kinase activity because mice expressing the deletion mutant with the K43R point mutation (K43R:del) and the WT-expressing mice had similar sleep phenotypes (**Figure 4b, c**). These results support that the constitutive kinase activity of CaMKIIβ induces sleep. Furthermore, sleep induction by CaMKIIβ does not require the dodecameric structure of CaMKIIβ or the regulatory segment and the linker region.

**Figure 4.**
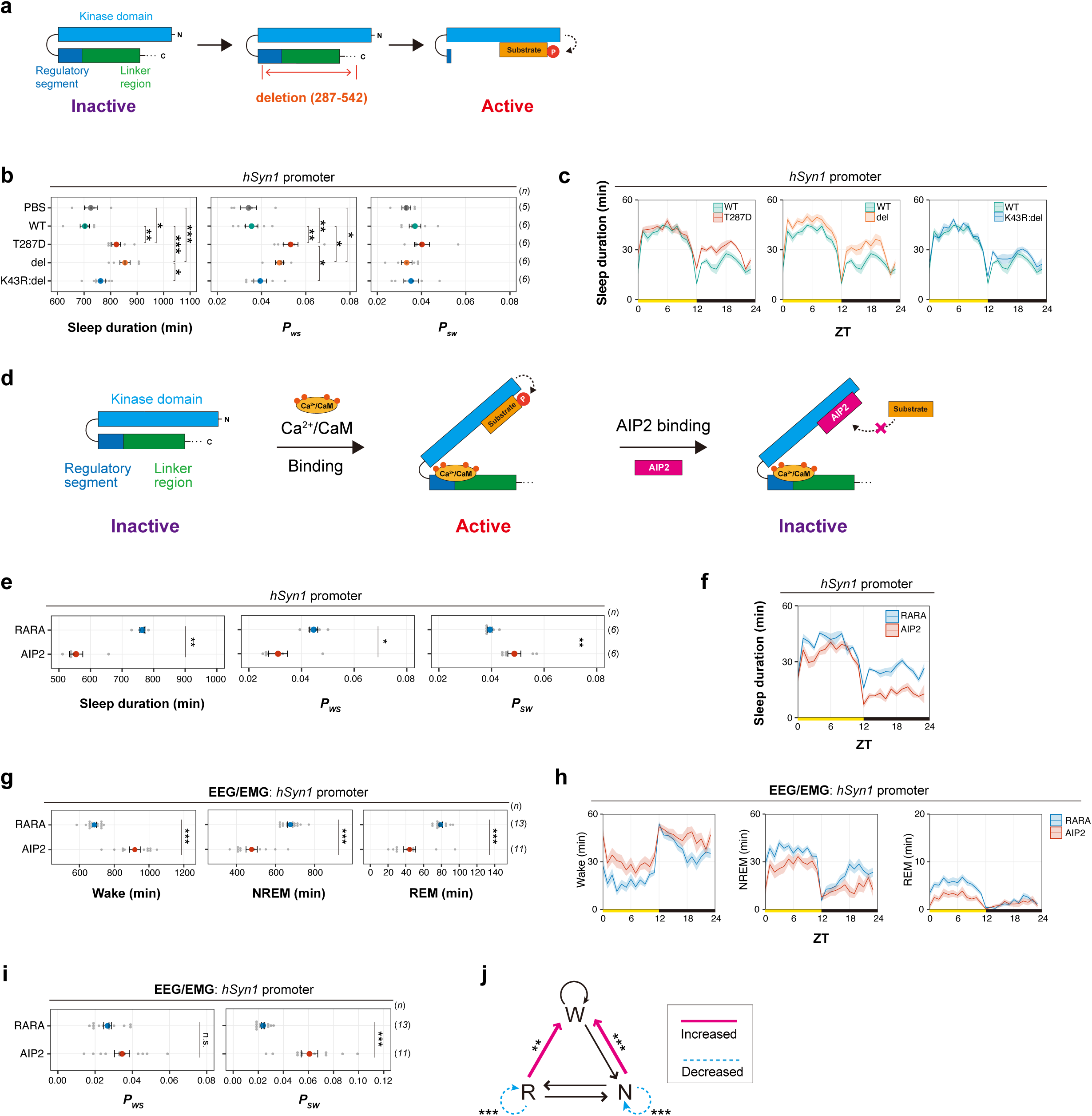
Perturbation on the kinase activity of CaMKIIβ bidirectionally affects sleep duration. **(a)** Schematic diagram of CaMKIIβ activation via deletion of its C-terminus. Deletion of the regulatory segment and linker region exposes the kinase domain of the CaMKIIβ and makes the enzyme constitutively activated. **(b-c)** Sleep/wake parameters **(b)** and sleep profiles **(c)** of mice expressing the CaMKIIβ deletion mutant (del), averaged over six days. The shaded areas represent SEM. Multiple comparison tests were performed between all individual groups. **(d)** Schematic diagram of CaMKII inhibition by AIP2 expression. AIP2 competitively binds to the kinase domain and inhibits substrate phosphorylation. **(e-f)** Sleep/wake parameters **(e)** and sleep profiles **(f)** of mice expressing AIP2 or the RARA mutant measured by the SSS, averaged over six days. **(g-i)** Sleep phenotypes **(g and i)** and sleep profiles **(h)** of mice expressing AIP2 or the RARA mutant measured by EEG/EMG recordings. **(j)** Differences in transition probabilities (between wakefulness (W), NREM sleep (N), and REM sleep (R)) between mice expressing AIP2 or the RARA mutant. Magenta lines and dashed blue lines indicate when the values for the AIP2-expressing mice are significantly (p < 0.05) higher and lower, respectively. Error bars: SEM, *p < 0.05, **p < 0.01, ***p < 0.001, n.s.: no significance. **Figure 4-source data 1** Source data for Figure 4b, c, e, f, g, h, i, j

We carried out a complementary approach by inhibiting the kinase activity of endogenous CaMKII. We used autocamtide inhibitory peptide 2 (AIP2), which inhibits the enzyme activity of CaMKIIα and CaMKIIβ by binding to the kinase domain and inhibiting the substrate-enzyme interaction (**Figure 4d**) ^44, 45^. Mice expressing the mCherry-fused AIP2 exhibited a decreased sleep duration and *P_WS_* along with an increased *P_SW_* compared with mice expressing the inactive mutant of AIP2 (RARA) (**Figure 4e, f**), demonstrating that the CaMKIIα/CaMKIIβ kinase activity is critical for normal sleep induction and maintenance. These results were consistent with the phenotype of *Camk2a* or *Camk2b* knockout mice ^10^, except for the *P_SW_* change: the genetic knockout of *Camk2a* or *Camk2b* slightly decreased *P_SW_*. This difference might account for the postnatal and kinase activity targeted inhibition of CaMKIIα/CaMKIIβ by AIP2 expression.

We further investigated the architectural and qualitative sleep changes under suppressed CaMKIIα/CaMKIIβ activity. The EEG/EMG recording of mice expressing AIP2 showed a significant decrease in NREM and REM sleep duration (**Figures 4g-i**). The increased transition probability from NREM/REM to wake and decreased transition to keep NREM and REM episodes in AIP2-expressing mice suggested that CaMKIIα/CaMKIIβ inhibition impaired the maintenance mechanism of NREM/REM sleep (**Figure 4j**). There was no significant change in normalized delta power during NREM sleep (**Figure 4-figure supplement 1a, b**). Note that there were differences in the waveforms of the EEG represented by the increased power of slow-wave oscillations (0.5 Hz–1 Hz) in all three states of vigilance (**Figure 4-figure supplement 1c**), though no difference was observed in the local field potential recordings of awaking mice cortex with the adult deletion of both *Camk2a* and *Camk2b* ^46^. Consistent with the phenotype of AIP2-expressed mice, EEG/EMG analysis showed that *Camk2b* knockout mice had decreased NREM and REM duration (**Figure 4-figure supplement 1d-g**) as well as decreased *P_SW_*. The knockout mice were established in previous study ^10^ but not analyzed for the sleep phenotype by EEG/EMG recordings. *Camk2b* knockout mice might have a decreased delta power, although we could not conclude on this because the changes in delta power depend on the normalization procedure of the EEG power spectrum (**Figure 4-figure supplement 1h-k**). The reduced sleep duration in SSS by AIP2- expression or *Camk2b* knockout can be attributed to the reduced NREM sleep because NREM sleep constitutes the most portion of total sleep time, though CaMKIIα/CaMKIIβ may also have a role in the control of REM sleep as observed in reduced REM sleep duration in these EEG/EMG recordings.

### Multi-site phosphorylation of CaMKIIβ can cancel sleep induction

Supposing that the autophosphorylation of T287 in CaMKIIβ encodes information on sleep need, the encoded information should not be decoded when it is not required. We thus investigated whether the phosphorylation of additional residues could cancel the sleep-inducing function of T287-phosphorylated CaMKIIβ. To this end, we created a series of double-phosphomimetic mutants of CaMKIIβ, in which besides T287, we mutated one of the remaining 68 S or T residues to D. The screening of these double-phosphomimetic mutants *in vivo* identified several mutants that exhibit a sleep phenotype similar to WT-expressing mice (**Figure 5a**, **Figure 5-figure supplement 1a-b**). In other words, the additional D mutation cancels the sleep-inducing effect of T287D. We focused on the five mutants (+S26D, +S182D, +T177D, +T311D, and +S516D; hereafter, we refer to the double-mutants by the additional mutated residue preceded by a plus sign) with the top five closest sleep parameters to WT, even if they had transduction efficiencies comparable to that of T287D (**Figure 5-figure supplement 1c-e**). To confirm that the observed phenotype of these five mutants came from the phosphomimetic property of D, we evaluated the phenotypes of non-phosphomimetic A mutants. The +S26A, +S182A, and +T311A mutants lost the effect of the D substitution, supporting the idea that phosphorylation of S26, S182, and T311 cancels the sleep-inducing effect of the co-existing T287 phosphorylation. On the other hand, the sleep phenotypes of +T516A and +T177A mice were similar to those of +T516D, +T177D, and WT mice (**Figure 5b, c**). This indicates that both A and D substitutions for these residues disturb sleep inducing effect of co-existing T287D mutation and thus the effect of D mutant may not rely on its phosphomimetic property.

**Figure 5.**
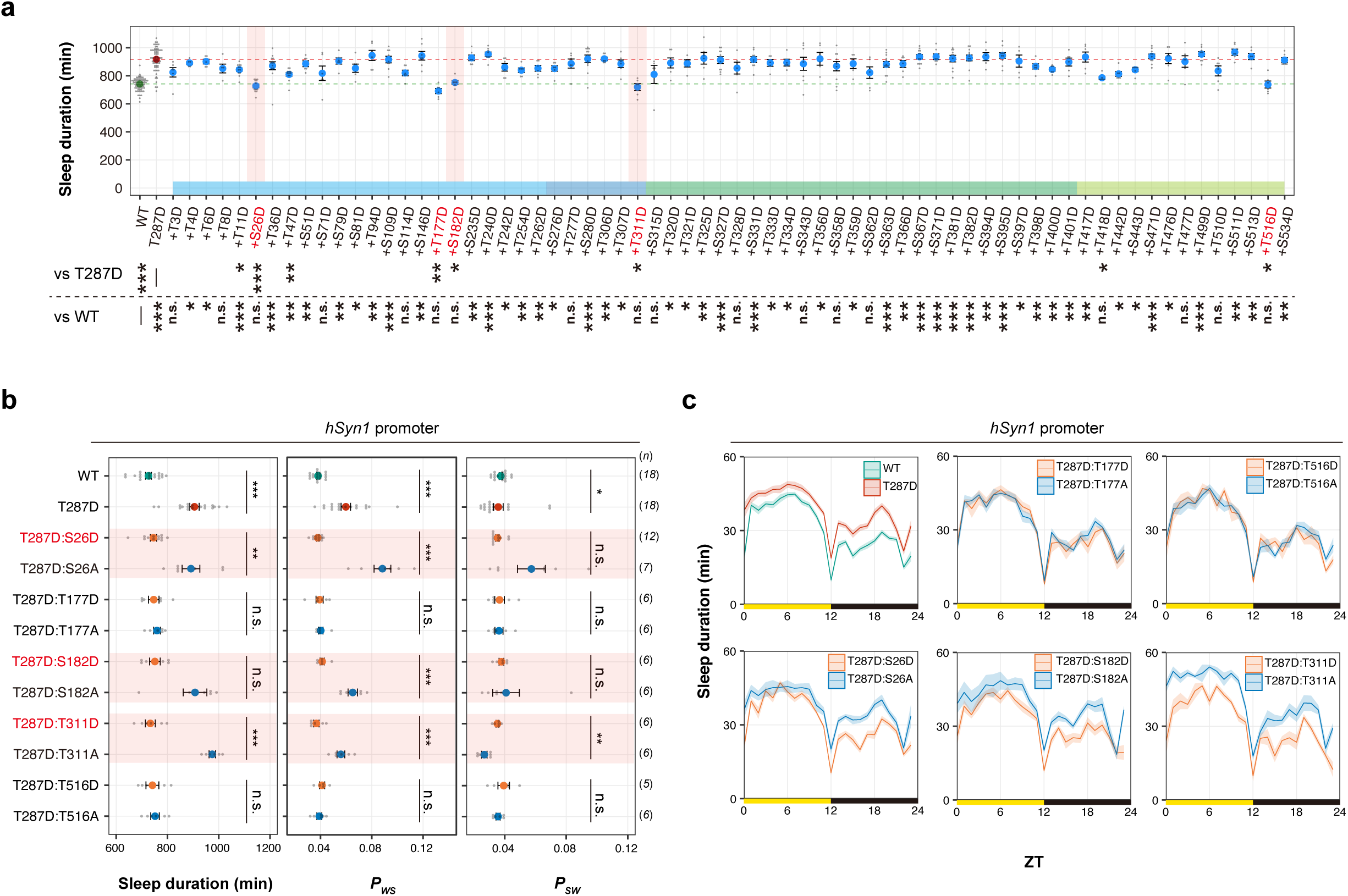
Multi-site phosphorylation of CaMKIIβ can cancel sleep induction. **(a)** Daily sleep duration, averaged over six days, of mice expressing CaMKIIβ double-phosphomimetic mutants (n = 5–12). The dashed green and red lines represent the averaged sleep duration of mice expressing wild-type CaMKIIβ (WT, n = 71) and T287D mutants (T287D, n = 68), respectively. The plus sign in a mutant’s name indicates a combination with T287D. Multiple comparison test was performed against WT (vs WT) or T287D (vs T287D). In the comparison with T287D mutant, “n.s.” labels are omitted for visibility. **(b-c)** Sleep/wake parameters **(b)** and sleep profiles **(c)**, averaged over six days, of mice expressing CaMKIIβ mutants with D or A substitutions of residues that cancel the sleep-inducing effect of T287D in **(a)**. Measurements are independent from those in **(a)**. The shaded areas represent SEM. Error bars: SEM, *p < 0.05, **p < 0.01, ***p < 0.001, n.s.: no significance. **Figure 5-source data 1** Source data for Figure 5a-c

### Biochemical evaluation of double-phosphomimetic CaMKIIβ mutants

We next evaluated the kinase activity of double-phosphomimetic CaMKIIβ mutants. Consistent with the result of single D mutants kinase assay (**Figure 1-figure supplement 2g**), T287D single mutant showed CaM-independent kinase activity and the level of CaM-dependent kinase activity is lower than that of wild-type (**Figure 5-figure supplement 1f**). Most of the double D mutants locates around the T287D suggesting that most of the second phosphomimetic mutations do not affect the kinase activity of T287D mutant significantly. It can also be seen that there is a correlation between CaM-dependent and CaM-independent kinase activity for T287D and double D mutants. We do not exclude the possibility that this variation/correlation is due to incomplete correction of relative CaMKIIβ levels in the cell extracts using dot blot (**Figure 5-figure supplement 1g**).

However, several mutants showed phosphorylation activity that was markedly different from T287D, to an extent that is difficult to be explained by the technical limitations of adjusting expression levels. Mutants locates at the left-bottom corner of **Figure 5-figure supplement 1f** had negligible kinase activity similar to kinase dead K43R mutant. +T311D mutant impaired the kinase activity in the absence of CaM compared to T287D, but the kinase activity in the presence of CaM is similar to T287D, suggesting that +T311D mutant abolished the constitutive-active property of T287D single mutant but the kinase activity is not abolished significantly. +S71D showed markedly higher kinase activity in the presence and absence of CaM. The dot blot quantification (**Figure 5-figure supplement 1g**) indicated that +S71D showed elevated expression level in 293T, but the kinase assay using the cell lysates with adjusted CaMKIIβ expression level suggests that the apparent catalytic rate constant for the kinase reaction of +S71D mutant is also elevated compared with T287D single mutant.

The comparison between kinase assay and double D mutant screening in vivo further supports that the constitutive and CaM-independent kinase activity is one of the factors responsible for the sleep-inducing effect and its cancellation. +S26D, +T47D, +T177D, +S182D, +T311D, and +T516D are the top 5 potential T287D-canceling mutants suggested by the AAV-based screening (**Figure 5a**). At least four of these five mutants had impaired kinase activity in the absence of CaM (i.e., +S26D, +T47D, +T177D, +S182D, and +T311D), and +T516D also showed reduced kinase activity compared with T287D mutant. The fact that +T311 shows kinase activity in the presence of CaM might indicate that the CaM-independent kinase activity is rather more important for the sleep phenotype in our AAV-based in vivo screening. Among these five mutants, four mutants (+S26D, +T47D, +S182D, +T311D, and +T516D) except for +T177D showed reduced expression level in 293T cells, and thus we could not evaluate the apparent catalytic rate constant for these low-expressed mutants. It is highly possible that the sleep cancelation effect of these mutants is mediated by the reduced expression level rather than the reduced catalytic constant, although there should be a considerable difference between protein expression levels in human cultured cell line and those in mice brain.

It should be noted that there are several mutants showing kinase activity that cannot be fully reconciled with the results of AAV-based screening *in vivo*. For example, although sleep canceling mutants (e.g., +S26D, +T47D etc) had the reduced kinase activity especially in the absence of CaM, there are also several mutants showing the very low kinase activity (e.g., +T8D, +S81D etc) but exhibit sleep-promotion effect comparable to the level of T287D single mutant. The reasons of these differences between in vivo phenotype and in vitro kinase activity are currently unknown.

### Multi-site phosphorylation of CaMKIIβ regulates sleep stabilization

Sleep duration and probabilities between sleep and awake phase switching (i.e., *P_WS_* and *P_SW_*) can be altered independently. For example, both *P_WS_* and *P_SW_* can have increased value without markedly changing sleep duration as observed in *Hcrt* knockout mice ^24^. The sleep-wake dynamics underlying the extended sleep duration can be subdivided into two types by using *P_WS_* and *P_SW_*: one is increased sleep “induction” activity characterized by an increase in *P_WS_* (higher probability of switching from awake phase to sleep phase). The other is increased sleep “maintenance” activity characterized by decrease *P_SW_* (lower probability of switching from sleep phase to awake phase). The T287D single mutant increases *P_WS_*, which can be categorized as an elevated sleep induction activity. Interestingly, we noticed that several double-mutants showed extended sleep duration due to an elevated sleep maintenance activity rather than sleep induction activity. **Figure 6a** shows the double-mutants plotted according to their *P_SW_* and *P_WS_*. The “T287D-canceling” mutants such as +S26D, +S182D, and +T311D locate close to WT. Notably, several mutants such as +T306D and +T307D locate at the bottom-left corner of the *P_SW_*- *P_WS_* plot, indicating that these mutants had lower *P_WS_* and *P_SW_* compared with single T287D mutants. In other words, the extended sleep duration of these double-mutants can lie in the increased sleep maintenance activity (i.e., decreased *P_SW_*) rather than sleep induction activity. The double mutants locate at the bottom-left corner can be categorized through clustering analysis indicated as “cluster III” (**Figure 6-figure supplement 1a**). Among the seven double mutants categorized as cluster III, T287D:T306D, T287D:T307D, and T287D:S534D robustly exhibited prolonged sleep duration, unchanged *P_WS_*, and reduced *P_SW_* compared with WT- expressing mice in the independent experiment (**Figure 6b** and **Figure 6-figure supplement 1b, c**). As the reduced *P_SW_* suggests, these three mutants prolonged sleep episode duration, indicating that they stabilize sleep (**Figure 6c**). We focused on the sleep maintenance function of T306 and T307 because these residues are a well-known autonomous negative-feedback control for CaMKIIβ kinase activation. We substituted these residues with the non-phosphomimetic residue alanine. The T287D:T306D:T307D and T287D:T306A:T307A mice both exhibited extended sleep duration compared to the WT (**Figure 6d**). As with T287D single mutant, the prolonged sleep duration for the T287D:T306A:T307A can be explained by an increase in *P_WS_* (i.e., sleep induction). However, the T287D:T306D:T307D mice showed decreased *P_WS_* and *P_SW_*, indicating that the extended sleep duration can be explained by sleep maintenance rather than the sleep induction (**Figure 6d and Figure 6-figure supplement 1d, e**). In support of this, T287D:T306D:T307D showed prolonged sleep episode duration (**Figure 6e**). The difference between T287D, T287D:T306D:T307D and T287D:T306A:T307A can be clearly visualized in the *P_WS_* and *P_SW_* plot (**Figure 6d**). A similar relationship can be observed between T287D:T306D and T287D:T306A mutants.

**Figure 6.**
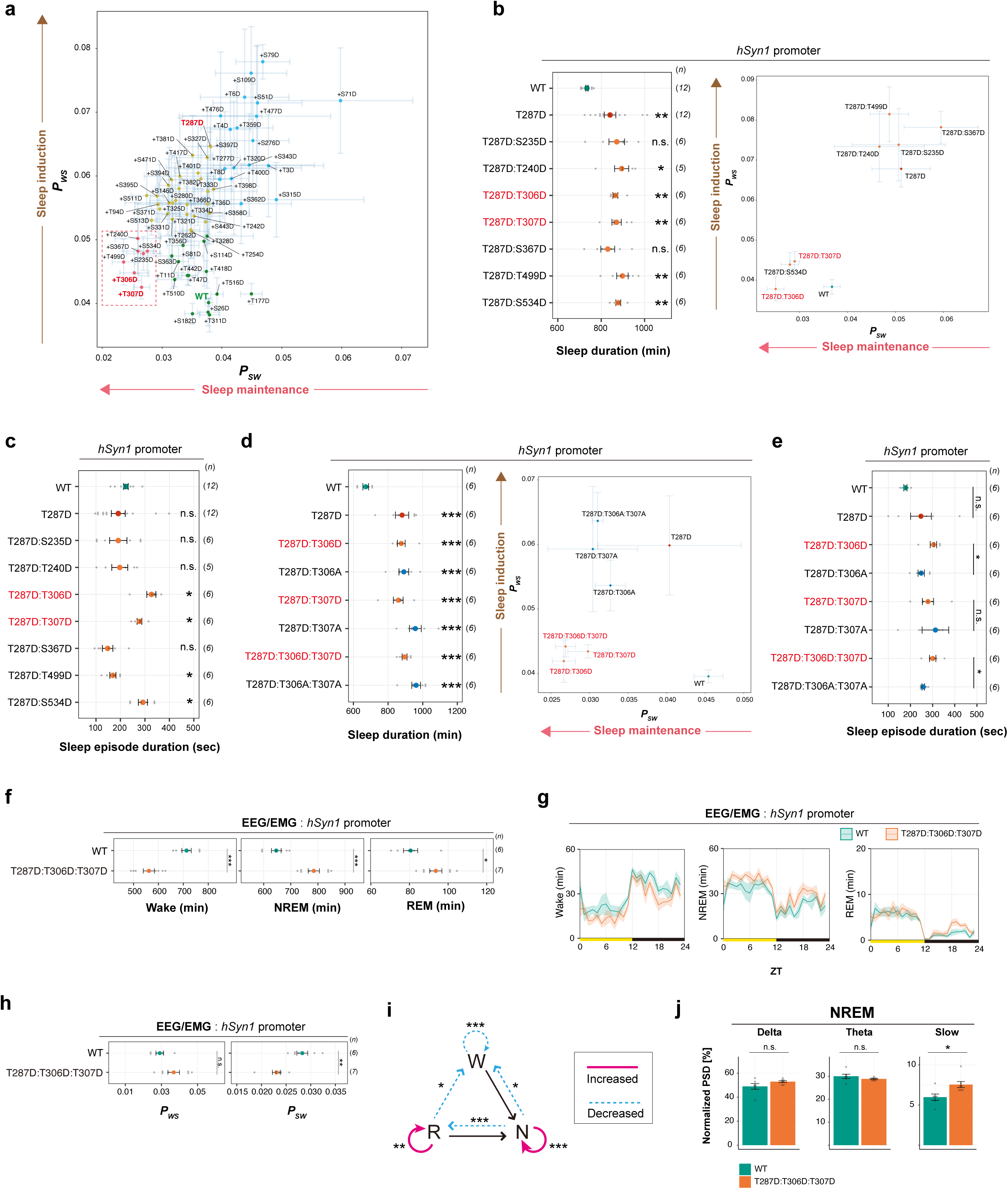
Multi-site phosphorylation of CaMKIIβ regulates sleep stabilization. **(a)** Correlation diagram of daily *P_WS_* and *P_SW_* of mice expressing the CaMKIIβ double-phopshomimetic mutants shown in Figure 5a, averaged over six days. The color of the dots correspond to the result of the clustering shown in **Figure 6-figure supplement 1a**. The mutants in the dotted magenta box had extended sleep duration with lower *P_WS_* and *P_SW_* (i.e., higher sleep maintenance activity). **(b)** Sleep duration and correlation diagram of daily *P_WS_* and *P_SW_* of mice expressing double-phopshomimetic mutants with sleep maintenance activity. Measurements are independent from those in Figure 5a. For the comparisons of sleep duration, multiple testing was performed against wild-type CaMKIIβ-expressing mice (WT). **(c)** Sleep episode duration, averaged over six days, of mice expressing the double-phopshomimetic mutants shown in Figure 6b **(d)** Sleep duration and correlation diagram of daily *P_WS_* and *P_SW_* of mice expressing CaMKIIβ mutants with D or A substitutions of sleep-stabilizing residues. For the comparisons of sleep duration, multiple testing was performed against wild-type CaMKIIβ-expressing mice (WT). **(e)** Sleep episode duration, averaged over six days, of mice expressing CaMKIIβ mutants with D or A substitutions of sleep-stabilizing residues shown in Figure 6d **(f-h)** Sleep phenotypes of mice expressing WT CaMKIIβ or the T287D:T306D:T307D mutant measured by EEG/EMG recordings. **(i)** Differences in transition probabilities (between wakefulness (W), NREM sleep (N), and REM sleep (R)) between mice expressing WT CaMKIIβ or the T287D:T306D:T307D mutant. Magenta lines and dashed blue lines indicate when the values for the T287D:T306D:T307D-expressing mice are significantly (p < 0.05) higher and lower, respectively. **(j)** NREM power density in typical frequency domains of mice expressing WT CaMKIIβ and the T287D:T306D:T307D mutant. Error bars: SEM, *p < 0.05, **p < 0.01, ***p < 0.001, n.s.: no significance. **Figure 6-source data 1** Source data for Figure 6b-j

We analyzed the architectural changes of sleep caused by the sleep-stabilizing mutant T287D:T306D:T307D using EEG/EMG recordings. The results showed an increase in NREM and REM sleep duration (**Figure 6f** and **6g**), a significant decrease in *P_SW_*, and no significant change in *P_WS_* (**Figure 6h**), which is consistent with the SSS analysis. These mice had higher NREM to NREM and REM to REM transition probabilities than WT-expressing mice. However, unlike T287D, this mutant did not increase the wake to NREM transition probability (**Figure 6i** and **Figure 2-figure supplement 1h**), suggesting that the additional phosphorylation(s) of T306 and/or T307 stabilize NREM and REM sleep. Mice expressing the T287D:T306D:T307D mutant and those expressing WT had similar delta power, but the mutant increased slow power (**Figure 6j** and **Figure 6-figure supplement 1f**). Thus, phosphorylation of T306/T307 also seems to elevate sleep need levels.

Phosphorylation of T306 and T307 in CaMKIIβ suppresses the kinase activity by inhibiting CaM binding ^27, 47^. To test whether the sleep maintenance function of the T287D:T306D:T307D mutant depends on its enzyme activity, we examined the sleep phenotype of mice expressing its kinase-dead version (K43R:T287D:T306D:T307D) and found that these mice did not exhibit a sleep-stabilizing phenotype. They had similar sleep parameters to the WT (**Figure 6-figure supplement 1g**). Furthermore, the T287A:T306D:T307D mutant, in which T287 was replaced by a non-phosphomimetic A, also resulted in similar sleep parameters to WT. These results suggest that the sleep maintenance function of CaMKIIβ with phosphorylated T306 and T307 depends on its enzyme activity and that this function requires T287 phosphorylation. We thus propose that multi-site phosphorylation of CaMKIIβ (residues T287, T306, and T307) converts the sleep-inducing effect of T287-phosphorylated CaMKIIβ into a sleep maintenance activity.

### Biochemical evaluation of sleep-stabilizing CaMKIIβ mutants

We then examined *in vitro* kinase activity of these sleep-stabilizing multiple D mutants and corresponding A mutants (**Figure 6-figure supplement 2a and b**). Consistent with the role of phosphorylation at T306 and T307 for the inhibition of the interaction with Ca^2+^/CaM to CaMKIIβ, mutants having the D substitution at either of T306 or T307 (i.e., T306D, T307D, T306D:T307D, T287D:T306D, T287D:T307D, T287D:T306D:T307D) showed reduced kinase activity in the presence of CaM. On the other hand, any mutants having the T287D mutation including sleep-stabilizing mutants annotated in AAV-based analysis (i.e., T287D:T306D and T287D:T306D:T307D) showed CaM-independent kinase activity compared with wild-type. This is also consistent with the role of T306/T307 phosphorylation because these phosphorylation does not actively inhibit the kinase activity of CaMKIIβ, and thus the CaM-independent activity of T287D mutant should be maintained if T287D is combined with T306D and/or T307D. The CaM-independent kinase activity was more evident with another substrate called autocamtide-2 (**Figure 6-figure supplement 2b**).

By contrast, kinase activity of mutants having the A substitution at T306 or T307 will need to be carefully interpreted. Introducing A substitution to either or both of T306 and T307 results in the CaM-independent kinase activity without having the T287D mutation (i.e., T306A, T307A, or T306A:T307A). We speculate that such CaM-independent activity might be caused by autophosphorylation of CaMKIIβ in the 293T cell. 293T cell expresses endogenous CaM protein. Although the cell-endogenous CaM is not sufficient to fully activate the over-expressed CaMKIIβ, it is reasonable to assume that there is a background level of CaM-dependent activation of CaMKIIβ in the 293T cells. Because T306A or T307A mutation impairs the auto-inhibitory mechanism, the T306A, T307A, or T306A:T307A mutants would be more susceptible to CaMKIIβ activation, which occurs at a lower efficiency in the 293T cell. Therefore, by the time 293T cell lysates are prepared, some portion of T306A, T307A, or T306A:T307A mutants may already be in an autonomously activated state with autophosphorylation at T287 residue.

### Ordered multi-site phosphorylation of CaMKIIβ underlies multi-step sleep regulation

The above *in vivo* analysis proposes that different CaMKIIβ phosphorylation states can induce sleep (T287), maintain sleep (T287:T306:T307), and cancel sleep promotion (S26:T287, S182:T287, and T287:T311). We assumed that phosphorylation at T287 precedes the other phosphorylations. We then aimed to biochemically confirm the ordered multi-site phosphorylation. We analyzed the time course changes in the phosphorylation levels of each sleep-controlling residues in CaMKIIβ (S26, S182, T287, T306, T307, and T311). The purified CaMKIIβ was incubated with CaM under four conditions with different concentrations of Ca^2+^ in the reaction buffer. Condition #1: 0 mM Ca^2+^ and 10 mM EGTA, supposing the presence of a negligible amount of free Ca^2+^. Condition #2: 0 mM Ca^2+^, supposing the presence of low Ca^2+^ concentration, possibly coming from the purified CaMKIIβ and/or CaM. Condition #3: 0.5 mM Ca^2+^, assuming a sufficient amount of free Ca^2+^ to activate CaMKIIβ. Condition #4: 0.5 mM Ca^2+^ and 10 mM EGTA at 5 min, where EGTA was added 5 min after incubation started. This type of condition induces the phosphorylation of T305 and T306 upon CaMKIIα activation ^48, 49^. Although we could detect the peak corresponds to S182 phosphorylation appeared during the CaMKIIβ incubation, it is hard to clearly separate the chromatogram of the peptide with S182 phosphorylation and that with the adjacent T177 phosphorylation (**Figure 7-figure supplement 1a**), so the quantification value of pS182 presented below include the signal from pT177 peptides.

**Figure 7a** indicates that T287 phosphorylation occurs in the presence of 0.5 mM Ca^2+^ (conditions #3 and #4), but not in the absence of explicitly added Ca^2+^ in the reaction buffer (conditions #1 and #2). The level of phosphorylation reaches a saturation level 5 min after CaM addition. Under condition #3, S26 and S182 phosphorylations follow T287 phosphorylation. However, conditions #1 and #4 do not phosphorylate these residues.

**Figure 7.**
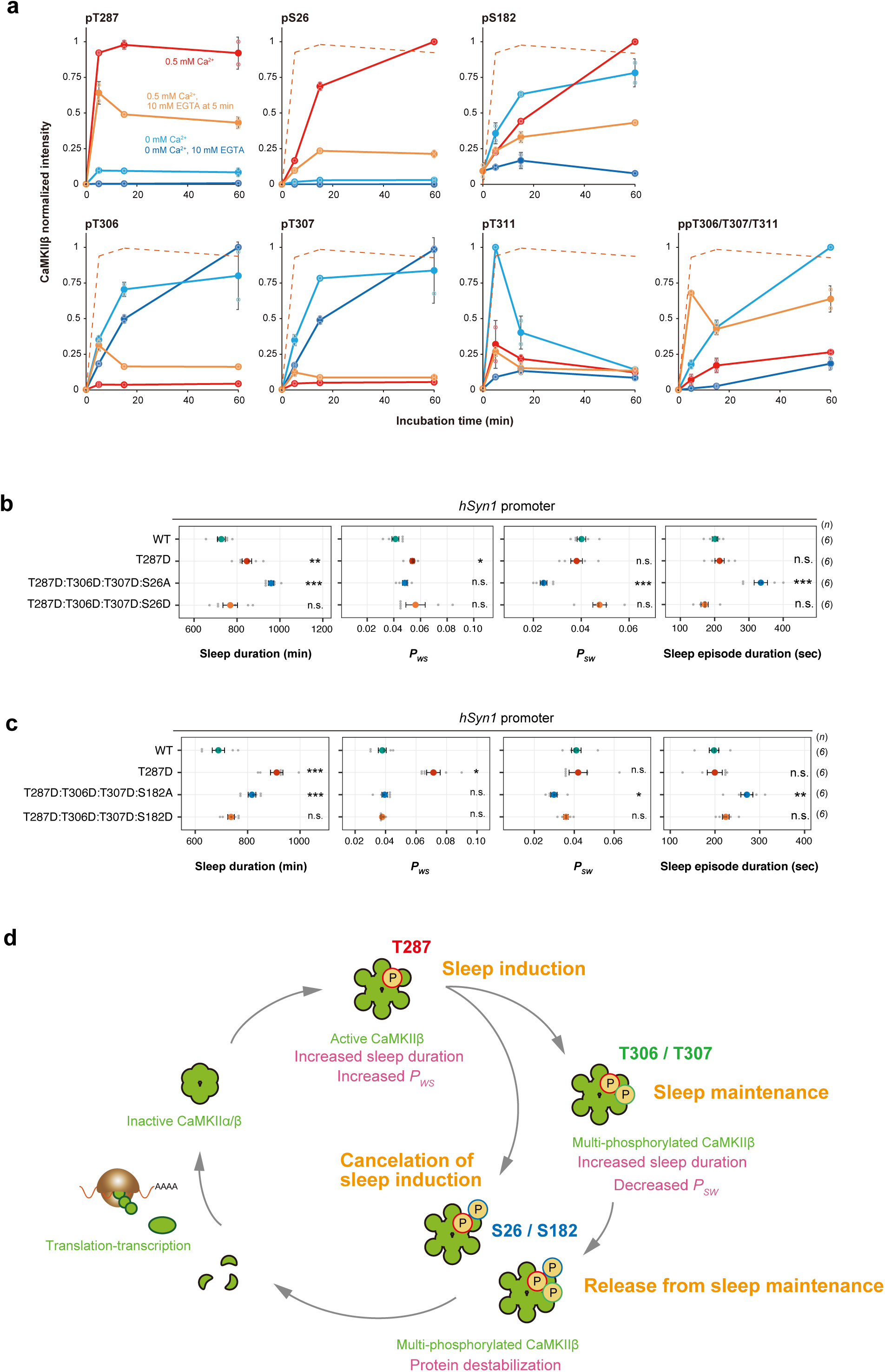
Ordered multi-site phosphorylation of CaMKIIβ underlies multi-step sleep regulation. **(a)** Time series changes of sleep-controlling residues phosphorylation under different Ca^2+^ conditions *in vitro*. The represented values are the mean ± SD (n = 2 independent experiments). The signal intensity of the detected peptides was normalized to the maximum value in the time series. The quantified values at 0 min were obtained from the sample before adding CaM and were shared in every Ca^2+^ conditions. The dashed lines trace the dynamics of T287 phosphorylation in the 0.5 mM Ca^2+^ condition. **(b-c)** Sleep/wake parameters of mice expressing quadruple-phosphomimetic CaMKIIβ mutants related to S26 **(b)** and S182 **(c)**. Multiple comparison test was performed against wild-type CaMKIIβ-expressing mice (WT). Error bars: SEM, *p < 0.05, **p < 0.01, ***p < 0.001, n.s.: no significance. **(d)** Ordered multi-site phosphorylation states of CaMKIIβ in sleep regulation. Note that this model only describes about the possible relationship between CaMKIIβ and the sleep-wake cycle without considering the difference between NREM and REM sleeps. **Figure 7-source data 1 and 2** Source data for Figure 7a. **Figure 7-source data 3** Source data for Figure 7b-c.

On the other hand, T306 and T307 remain unphosphorylated in the presence of a high amount of Ca^2+^ and CaM (condition #3). Shielding the Ca^2+^ after CaMKIIβ activation (condition #4) triggered T306 and T307 phosphorylation. This is consistent with previous studies suggesting that the stable binding of Ca^2+^/CaM renders T306 and T307 inaccessible to the kinase domain of CaMKIIβ, and their phosphorylation requires the temporal removal of Ca^2+^/CaM from the kinase ^48–51^. We also confirmed that the optimal Ca^2+^ concentration for T306 and T307 phosphorylation is lower than that for T287 phosphorylation: gradual phosphorylation of T306 and T307 occurs in the absence of apparent Ca^2+^ in the reaction buffer (condition #1 and #2) ^47, 52^. The low Ca^2+^ concentration condition also promotes T311 phosphorylation, which is spacially close to T306 and T307. The time course of T311 phosphorylation in condition #2 is different from that of T306 and T307 phosphorylation: the phosphorylation of T311 peaked 5 min after CaM addition and then decreased, presumably because of the progressive phosphorylation of T306 and T307. It is unlikely that a misregulated, Ca^2+^/CaM-independent kinase activity phosphorylated T306, T307, and T311 under low Ca^2+^ concentration conditions because chelating Ca^2+^ with EGTA abolishes the appearance of double-phosphorylated T306/T307/T311 peptides (condition #1; ppT306/T307/T311). In summary, the biochemical analysis suggests that T287 phosphorylation initiates the ordered phosphorylation of S26, S182, T306, T307, and T307 (**Figure 7-figure supplement 1b**) at least in our *in vitro* experimental condition.

With the ordered phosphorylation events observed *in vitro*, the CaMKIIβ might reach multi-phosphorylated states such as pS26:pT287:pT306:pT307, pS182:pT287:pT306:pT307, or pT287:pT306:pT307:pT311 *in vivo*. To investigate the effect of such multi-phosphorylated states in sleep regulation, we expressed CaMKIIβ mutants mimicking quadruple-phosphorylation in mice. Inclusion of S26A or S182A to the T287D:T306D:T307D recapitulated the sleep maintenance function observed in T287D:T306D:T307D mutant (**Figures 7b, c** and **Figure 7-figure supplement 1c, d**), with decreased *P_SW_* and a prolonged sleep episode duration. On the other hand, the substitution of in S26 or S182 to D resulted in the loss of the sleep maintenance function. The mutant with the T311D substitution added to the T287D:T306D:T307D retained sleep maintenance activity (**Figure 7-figure supplement 1e**). Therefore, the sleep induction and maintenance effect of CaMKIIβ elicited by T287 phosphorylation followed by T306 and T307 phosphorylation appears to be terminated by S26 and S182 phosphorylation, which also follows T287 phosphorylation. Based on these results, we propose that the ordered multi-phosphorylation states of CaMKIIβ underly the sleep regulation steps, namely the induction (pT287), the maintenance (pT287/pT306/pT307), and the cancelation (pS182 or pS26). These multi-site phosphorylation states might be connected, and finally completed as a cycle by the turnover of phosphorylated CaMKIIβ promoted by the protein destabilization effect of S182 or S26 phosphorylation (**Figure 7d**).

## Discussion

In this study, we demonstrated that the conditional induction or inhibition of CaMKIIβ kinase activity could bidirectionally increase or decrease mammalian sleep duration. The bidirectional effect as well as the near two-fold difference in sleep duration caused by the activation (e.g., 936.7 ± 22.6 min; **Figure 2a**) and inhibition (e.g., 554.1 ± 21.2 min; **Figure 4e**) of CaMKIIβ further supports the role of CaMKIIβ as a core sleep regulator, rather than auxiliary inputs that either induce or inhibit sleep upon environmental responses. Assuming the role of CaMKIIβ as one of the core kinases in the sleep control, the next question would be how CaMKIIβ relates to other phosphorylated enzymes, such as CaMKIIα ^10^, SIK1/SIK2/SIK3 ^15, 16^, and ERK1/ERK2 ^17^, to shape the phosphorylation signaling network for sleep regulation.

The postnatal conditional expression of CaMKIIβ and its inhibitor changes the sleep phenotype, which rules out, at least in part, neuronal developmental abnormality potentially caused by the embryonic knockout of *Camk2b* ^53^. Although the embryonic double knockout of *Camk2a*/*Camk2b* caused developmental effects ^46^, the sleep reduction caused by the conditional expression of CaMKII inhibitor AIP2 supports that the reduction of kinase activity reduced sleep duration in the *Camk2a* KO and *Camk2b* KO mice ^10^, not the neuronal structural abnormality potentially caused by the gene knockout. Given the inducible adult deletion of both *Camk2a* and *Camk2b* resulted in lethal phenotype ^46^, our AIP2 expression condition would only partially inhibit the kinase activity of CaMKIIα and CaMKIIβ.

Third, the effect of AIP2 and kinase-inhibitory CaMKIIβ mutants (e.g., K43R and S26D) indicate that the sleep-promoting effect of activated CaMKIIβ comes from the enzymatic activity of CaMKIIβ (**Figure 4**). The sleep-promoting effect of the truncated CaMKIIβ kinase domain further indicates that CaMKIIβ oligomerization is not necessary for the sleep-promoting effect. This is in stark contrast with the non-enzymatic role of CaMKIIβ through its interaction with F-actin ^54, 55^. The truncated CaMKIIβ used in this study lacks the actin binding domain. Another well-known binding partner of CaMKIIα/β is NR2B ^56^, which has a low affinity for monomeric CaMKIIα ^57^. Therefore, the potent sleep-inducing effect of truncated CaMKIIβ suggests that other downstream targets (such as phosphorylation substrates) are responsible for the sleep-inducing effect of CaMKIIβ. Future research should focus on identifying such downstream targets, but at least the present study excludes the core circadian transcription factors and functional transcription-translation circadian feedback loop as downstream factors of CaMKIIβ sleep promotion (**Figure 2**).

Finally, comparing phosphorylation-mimicking mutants and non-phosphorylation-mimicking mutants allowed us to attribute the effect of phosphorylation to the negative charge mimicked by the D residue or to any other effect caused by the mutation. As observed in the SIK3 phosphorylation site S551 ^58^, D and A mutations sometimes yield similar results (e.g., increased sleep), making it difficult to conclude that the D mutation mimics phosphorylation. For residues analyzed in **Figure 7a**, we showed that A and D mutants had different effects in sleep regulation *in vivo*, suggesting that the phosphorylation states of these sites in CaMKIIβ can regulate sleep. To the best of our knowledge, this is the first conclusive demonstration of phosphorylation-dependent sleep regulation at single residue level. Besides, these residues are autophosphorylation substrates, at least *in vitro*. These results suggest that the multi-step effects of CaMKIIβ on sleep induction, sleep maintenance, and sleep promotion cancelation can be attributed to the properties of the CaMKIIβ with multiple (auto-)phosphorylation patterns.

The sleep-promoting effect observed with *Vglut2-Cre* but not *Gad2-Cre* (**Figure 3**) suggests that CaMKIIβ promotes sleep by acting on excitatory neurons rather than inhibitory neurons and glial cells. However, these data do not exclude the possibility of the contribution of non-excitatory neurons and glial cells for CaMKIIβ-dependent sleep regulation because the Cre-expression specificity may not be perfectly selective to desired cell types. Furthermore, endogenous *Camk2b* is widely expressed in neurons and constitutes ∼1.3% of postsynaptic density ^59^ and glial cells also express *Camk2b* ^60^. Future research will have to precisely elucidate where CaMKIIβ exerts its sleep function in terms of both neuronal cell types and brain regions as well subcellular localization. In the data shown in **Figure 3b**, focused expression of T287D to *Vglut2*-*Cre* positive cells might induce the sleep maintenance activity (i.e., extended sleep duration and low *P_SW_*) in addition to the sleep induction activity (i.e., high *P_WS_*), suggesting that different types of neurons might be involved in the sleep induction or maintenance activities to different degrees. Notably, homeostatic regulation of sleep/wake-associated neuronal firing was recapitulated in cultured neuron/glial cells ^61, 62^. Given the ubiquitous and abundant expression of CaMKIIβ in neurons, investigating the relationship between sleep homeostasis in cultured neurons/glial cells and CaMKIIβ phosphorylation states would reveal valuable information about the ubiquitous and cell-type specific function of CaMKIIβ in the sleep control.

Multi-site phosphorylation encodes complex biochemical systems such as the sequential triggering of multiple events and the integration of multiple signals (such as AND logic gates) ^63, 64^. One of the most intriguing properties of CaMKII is the multi-site autophosphorylation combinations that regulate kinase activity and protein-protein interactions. In this study, we conducted comprehensive mutagenesis of single or multiple potentially (auto-)phosphorylable residues. We revealed that the phosphorylation of kinase-suppressive residues can cancel the sleep-promoting effect of the active T287-phosphorylated CaMKIIβ (**Figure 5**). Sleep-suppressing mechanisms may include CaMKIIβ destabilization (e.g., through S182 phosphorylation) and other biochemical mechanisms inhibiting either the kinase activity or the CaMKIIβ-substrates interaction. The combination of sleep-promoting and sleep-suppressing phosphorylations of CaMKIIβ may underlie the mechanism regulating sleep need to an appropriate level, depending on the animal’s internal conditions and external environments. Considering this, it would be interesting to quantify the phosphorylation level of each residue (other than T287) in response to signals causing acute and chronic changes in the sleep-wake cycle (such as inflammation and stress).

Next, we found that combining phosphomimetic mutations of T306D and T307D to T287D (i.e., T287D:T306D:T307D) does not affect sleep duration (compared with the T287D single mutation) but causes unexpected differences in sleep maintenance and sleep induction (**Figure 6**). The transition probabilities (*P_WS_* and *P_SW_*) allowed us to quantify these interesting differences. For example, the T287D mutant has a higher *P_WS_*, suggesting that T287 phosphorylation plays a role in sleep induction. On the other hand, the T287D:T306D:T307D mutant has a lower *P_SW_*, suggesting that T287/T306/T307 phosphorylation plays a role in sleep maintenance. The autophosphorylation of T306 and T307 has a well-known inhibitory effect on the CaMKIIβ-CaM interaction ^27, 28^, creating an auto-inhibitory feedback regulation of CaMKIIβ. CaMKIIβ deletion mutant shares some properties with T287D:T306D:T307D; both mutants lost the CaMKIIβ-CaM interaction and have the CaM-independent kinase activity. Nevertheless, T287D:T306D:T307D has the sleep maintenance activity while the deletion mutant shows sleep induction activity. Thus, the mechanism of sleep maintenance by T287D:T306D:T307D may not be attributed to the loss of CaMKIIβ-CaM interaction itself. The outcome of CaMKIIβ kinase activity with different phosphorylation patterns and molecular mechanisms underlying the sleep induction/maintenance activities are currently unknown. Recent studies suggested that the phosphorylation of T305/T306 of CaMKIIα promotes the dissociation of CaMKIIα dodecamer ^65^. Another study demonstrated that the same phosphorylation promotes the translocation of CaMKIIα from the spine to dendrite ^66^. It is plausible that different patterns of multi-site phosphorylation or combination of D mutants of CaMKIIβ affect sleep induction/maintenance through the different interactions of endogenous CaMKIIα/CaMKIIβ and neuronal proteins.

We also showed that other sleep-controlling residues (such as S26, S182, and T311) also undergo autophosphorylation (**Figure 7a**). S26 autophosphorylation occurs in CaMKIIγ ^67^ and suppresses the kinase activity ^68^, which is consistent with our results in CaMKIIβ. The other phosphoproteomics study identified S25 autophosphorylation in CaMKIIα ^30^. Although the level of phosphorylation at S25 was indicated for ∼5% of total CaMKIIα at 4 min incubation time ^30^, it is possible that the level of this phosphorylation continuously increases given the slow dynamics of autophosphorylation at S26 of CaMKIIβ found in this study. We also note that peptide phosphorylated with S26 can be found *in vivo* brain sample ^12^ (**Figure 7-figure supplement 1f**), although it is unable to distinguish CaMKII isoforms because of the identical sequence around S26 phosphorylation site. Reports suggest that T311 autophosphorylation occurs in CaMKIIβ ^30^ and that the phosphorylation level of the corresponding residue in CaMKIIα was reduced during the dark phase (mostly awake phase in mice) ^11^. The T311 phosphorylation was also detected in the other set of phosphoproteomic analyses of in vivo mice brains ^12^, although sequence identity around the T311 residue makes it difficult to distinguish CaMKIIα and CaMKIIβ. These phosphoproteomics analyses support the possible role of phosphorylation at S26 or T311 in the regulation of CaMKIIα/CaMKIIβ in mice brains *in vivo*. To the best of our knowledge, our study is the first to report the autophosphorylation of S182. Furthermore, S26 and S182 autophosphorylation are slower than that of T287 (**Figure 7a**), consistent with the fact that these residues are not exposed on the surface and thus a kinase cannot easily access to these residues. The mammalian circadian clock regulation appears to use the non-canonical and inefficient phosphorylation residue to encode slower dynamics of circadian clock peacemaking ^69, 70^; it should be rigorously tested whether non-canonical autophosphorylation residues such as S26 and S182 plays a role in the regulation of normal sleep regulation *in vivo* through the knockout/knockdown rescue experiment by re-expressing the unphosphorylatable A mutations at corresponding residues. Through such rescue experiments, it would be possible to approach the question not covered by the current study: whether the sleep cancellation effect is related to the transition from sleep to awake phase in the natural sleep-wake cycle, or to the cancellation of additional sleep needs upon unusual input such as sleep deprivation.

Considering this sequential autophosphorylation of sleep-controlling residues, we aligned the different sleep-promoting effects elicited by each phosphorylation state with the autophosphorylation events (**Figure 7d**). The expected sleep regulation sequence is physiologically plausible: the increased transition rate from awake to sleep phase, the induced sleep is stabilized, and then the sleep-promoting effect is canceled. The cancelation may include complete erasure of multi-site phosphorylation through the destabilization of CaMKIIβ. Because both CaMKIIα and CaMKIIβ are involved in sleep control and have overlapping roles in the control of neural plasticity, the mechanism we found in this study may be shared by CaMKIIα as well as CaMKIIβ. On the other hand, it is also known that there are differences in the dynamics of phosphorylation of T306 and T307 between CaMKIIα and CaMKIIβ ^49^, and it will be interesting to investigate how these differences at the molecular level affect sleep-wake regulation.

This sequence is hypothetical at this stage, and it is still unknown whether the same CaMKIIβ molecule regulates the sequential events or different CaMKIIβ molecules with distinct phosphorylation states operate individually. It is also possible that phosphorylation on several non-canonical autophosphorylation residues (e.g., S26 and S182) is mediated by different kinases, and several residues may be rather effectively phosphorylated during the awake phase as observed in the T310 (CaMKIIα) or T311 (CaMKIIβ) residues. The obvious next question might be: how are the sleep-driven and wake-driven multi-site phosphorylation of each CaMKIIβ molecule integrated and organized by autophosphorylation and phosphorylation by other kinases, such that robust and flexible cycle of sleep induction, maintenance and subsequent transition from sleep to awake phase. Also, the multi-site phosphorylation status of CaMKIIβ might be the key to understand the connection between the sleep-wake cycle and its physiological significance. Indeed, phosphorylation mimicking or non-phosphorylation mimicking mutants of CaMKIIα/ CaMKIIβ have been shown to elicit defects in neuronal plasticity and some type of learning. Because it is well understood that the sleep-wake cycle affects the learning process, CaMKIIβ-expressing mice with changes in sleep phenotype may also have changes in learning phenotype. In this case, it would be interesting to ask whether the changes in learning phenotype are simply due to sleep abnormalities or whether CaMKIIβ plays a more direct role in these relationships as a molecule that controls both sleep and learning processes.

In summary, we showed that CaMKIIβ kinase activity promotes mammalian sleep by acting on the excitatory neurons. We propose that the ordered multi-site phosphorylation and kinase activity of CaMKIIβ compose the *input* (exposure of the kinase domain), *storage*/*processing* (T287 and following phosphorylations), and *output* (substrate phosphorylation) mechanism of sleep need in mammals. Hence, this could be the molecular mechanism of the phosphorylation hypothesis of sleep in mammals.

## FIGURE LEGENDS

**Figure 1-figure supplement 1.**
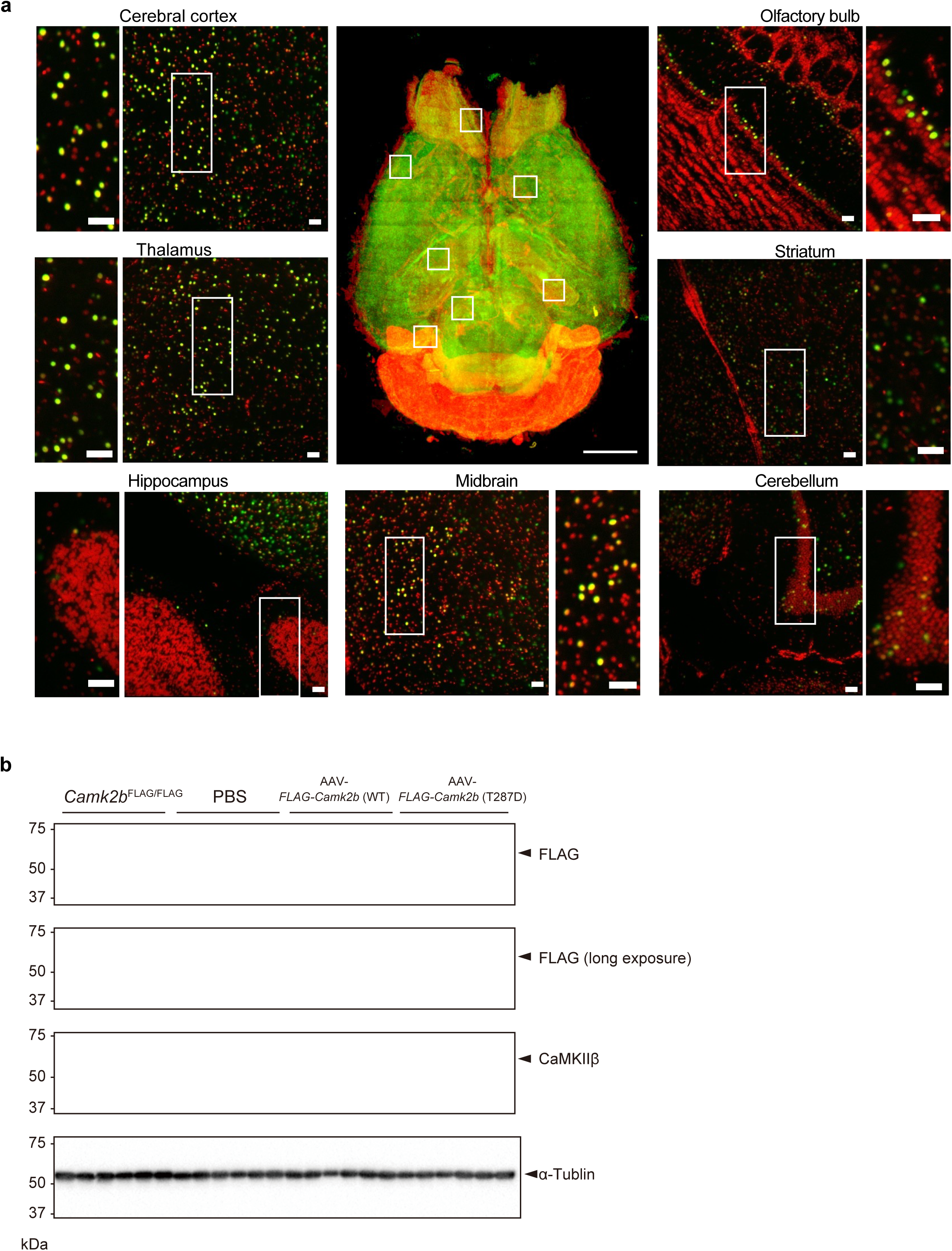
Expression of the CaMKIIβ throughout the brain by AAV-PHP.eB. **(a)** Volume-rendered and single-plane images of the brain expressing H2B-mCherry under *hSyn1* promoter by the AAV (mCherry, green) counterstained with RD2 (red). A volume-rendered image is shown in the center. Single-plane and magnified images are shown for cerebral cortex, thalamus, hippocampus, midbrain, cerebellum, striatum, and olfactory bulb. Scale bar in the center image, 3 mm; other scale bars, 100 µm. **(b)** Expression levels of endogenous CaMKIIβ and AAV-mediated transduced CaMKIIβ in the brain. *Camk2b*^FLAG/FLAG^ represents homo knock-in mice in which the FLAG tag was inserted into the endogenous *Camk2b* locus. PBS: PBS-administrated mice. Immunoblotting against FLAG-tagged protein indicates that AAV-mediated expression of CaMKIIβ is lower than the expression level of endogenous CaMKIIβ. **Figure 1-figure supplement 1-source data 1** Uncropped blot images for Figure 1-figure supplement 1b. **Figure 1-figure supplement 1-source data 2** Raw blot images for Figure 1-figure supplement 1b.

**Figure 1-figure supplement 2.**
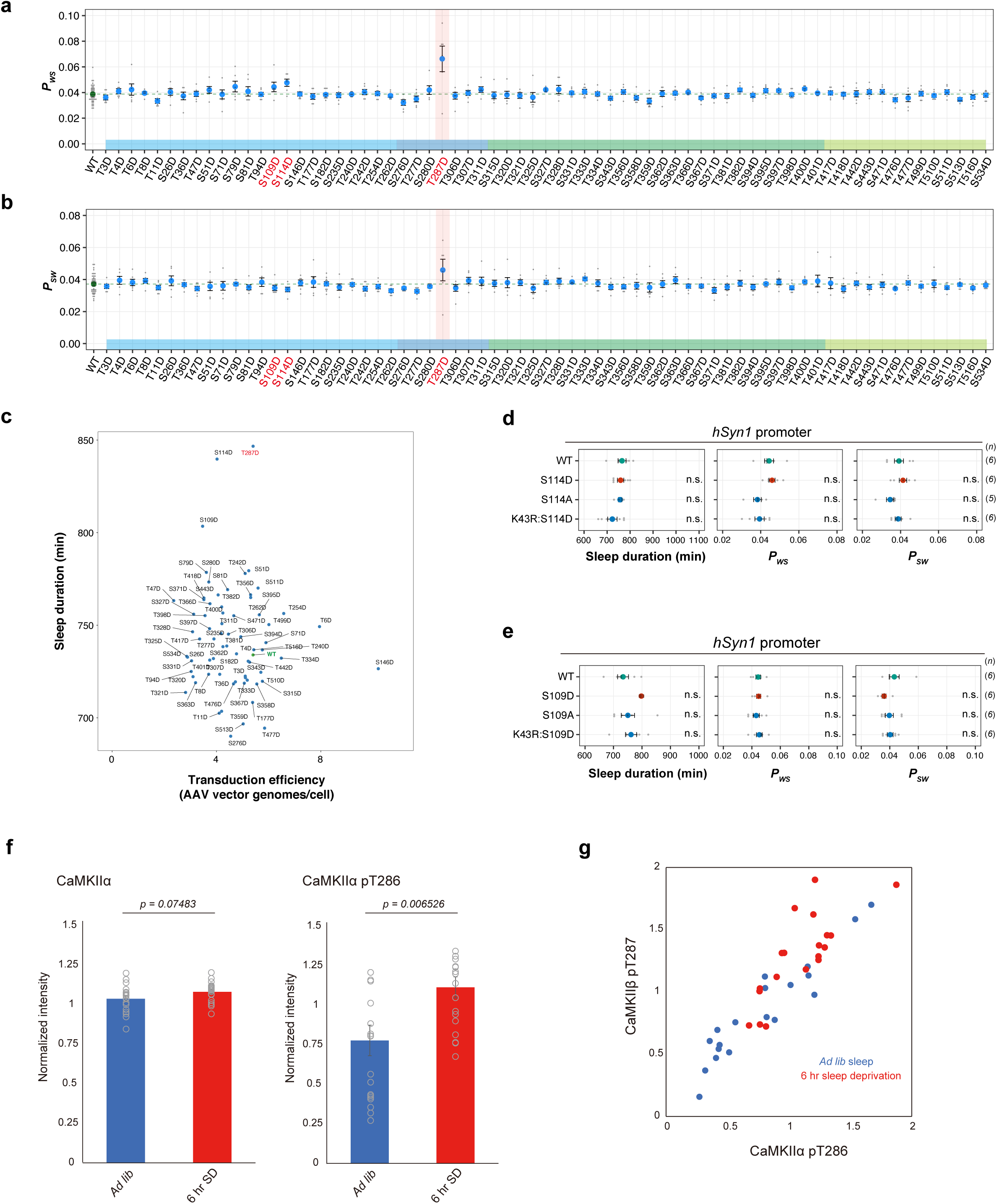
Phosphorylation of CaMKIIβ regulates sleep induction. **(a)** Daily *P*_WS_ **(a)** and *P*_SW_ **(b)** of mice expressing the CaMKIIβ phopshomimetic mutants (n = 6–10) shown in Figure 1c, averaged over six days. Dashed green lines represent averaged *P*_WS_ **(a)** and *P*_SW_ **(b)** of mice expressing wild-type CaMKIIβ (WT, n = 48). The multiple comparison test revealed no significant differences between mutants and WT. **(c)** Calculated transduction efficiency plotted against sleep duration. Transduction efficiency is an estimation of the number of AAV vector genomes present per cell in a mouse brain. After the SSS measurements, we purified the AAV vector genomes from the mice brains and then quantified them with a WPRE-specific primer set and normalized to mouse genomes. **(d-e)** Sleep/wake parameters of mice expressing S114-related CaMKIIβ mutants **(d)** and S109-related CaMKIIβ mutants **(e)**, averaged over six days. The shaded areas represent SEM. Multiple comparison test was performed against wild-type CaMKIIβ-expressing mice (WT). **(f)** Total CaMKIIα and T286-phosphorylated peptides from brains of sleep-deprived and control mice, analyzed by SRM quantitative mass spectrometry. Error bars: SEM **(g)** Correlation of phosphorylation of CaMKIIα T286 and CaMKIIβ T287 in each brain. Each point corresponds to the quantification value obtained from individual mouse brain. Error bars: SEM, *p < 0.05, **p < 0.01, ***p < 0.001, n.s.: no significance. **Figure 1-figure supplement 2-source data 1** Source data for Figure 1-figure supplement 2d, e, f **Figure 1-source data 2 and 3** These source data include source data for Figure 1-figure supplement 2f and 2g.

**Figure 1-figure supplement 3.**
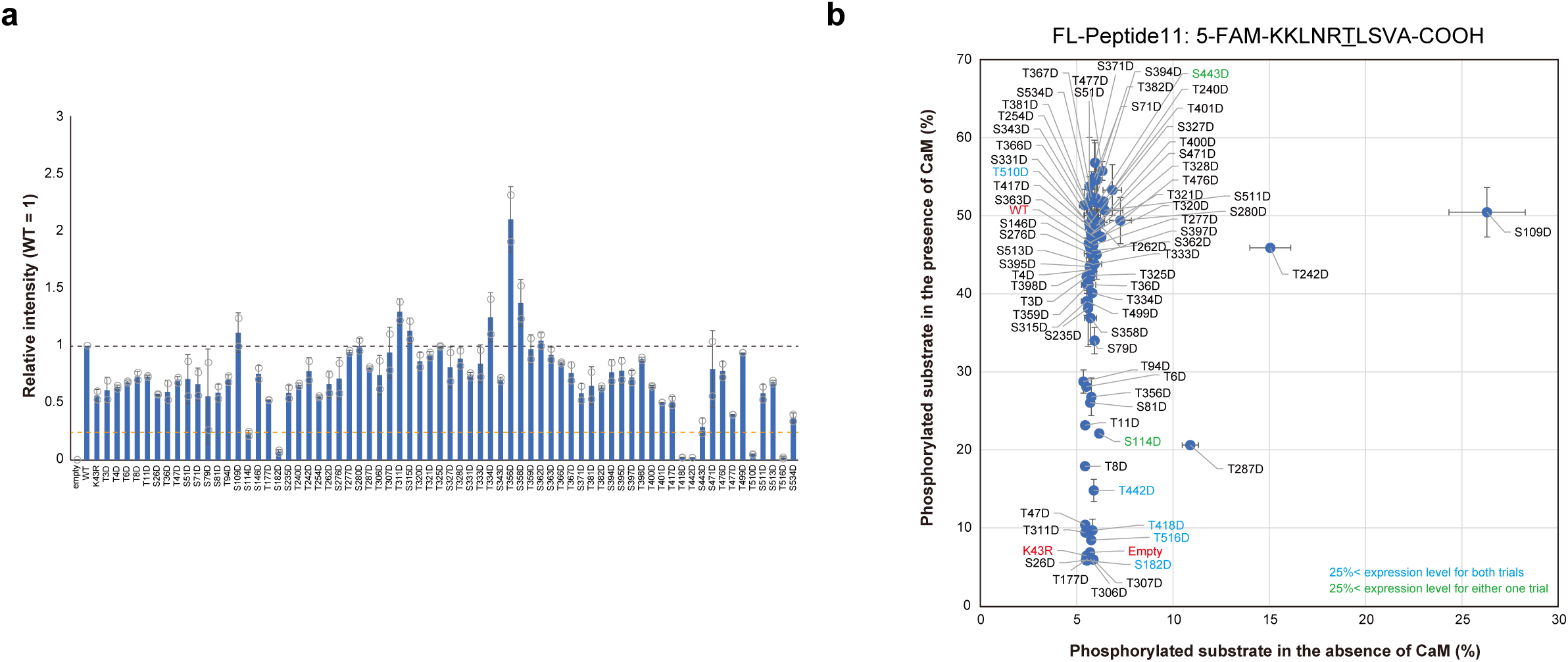
Biochemical evaluation of sleep-inducing CaMKIIβ mutants. **(a)** Expression levels of each mutant in the cell extracts used for the measurements shown in **(b)**. The expression level of each mutant was normalized relative to WT (black dashed line). The orange dashed line indicates 25% of the WT expression level. The represented values are the mean ± SD (n = 2, independent experiments). **(b)** *In vitro* kinase activity of CaMKIIβ phosphomimetic mutants. Phosphorylation (%) indicates the percentage of the phosphorylated substrate relative to the total peptide in the presence or absence of CaM. The represented values are the mean ± SD (n = 2, independent experiments). The mutants with blue labels exhibited < 25% lower expression than the WT in both trials. The mutants with green labels exhibited < 25% lower expression than the WT in one of the two trials. **Figure 1-figure supplement 3-source data 1** Source data for Figure 1-figure supplement 3a, b.

**Figure 2-figure supplement 1.**
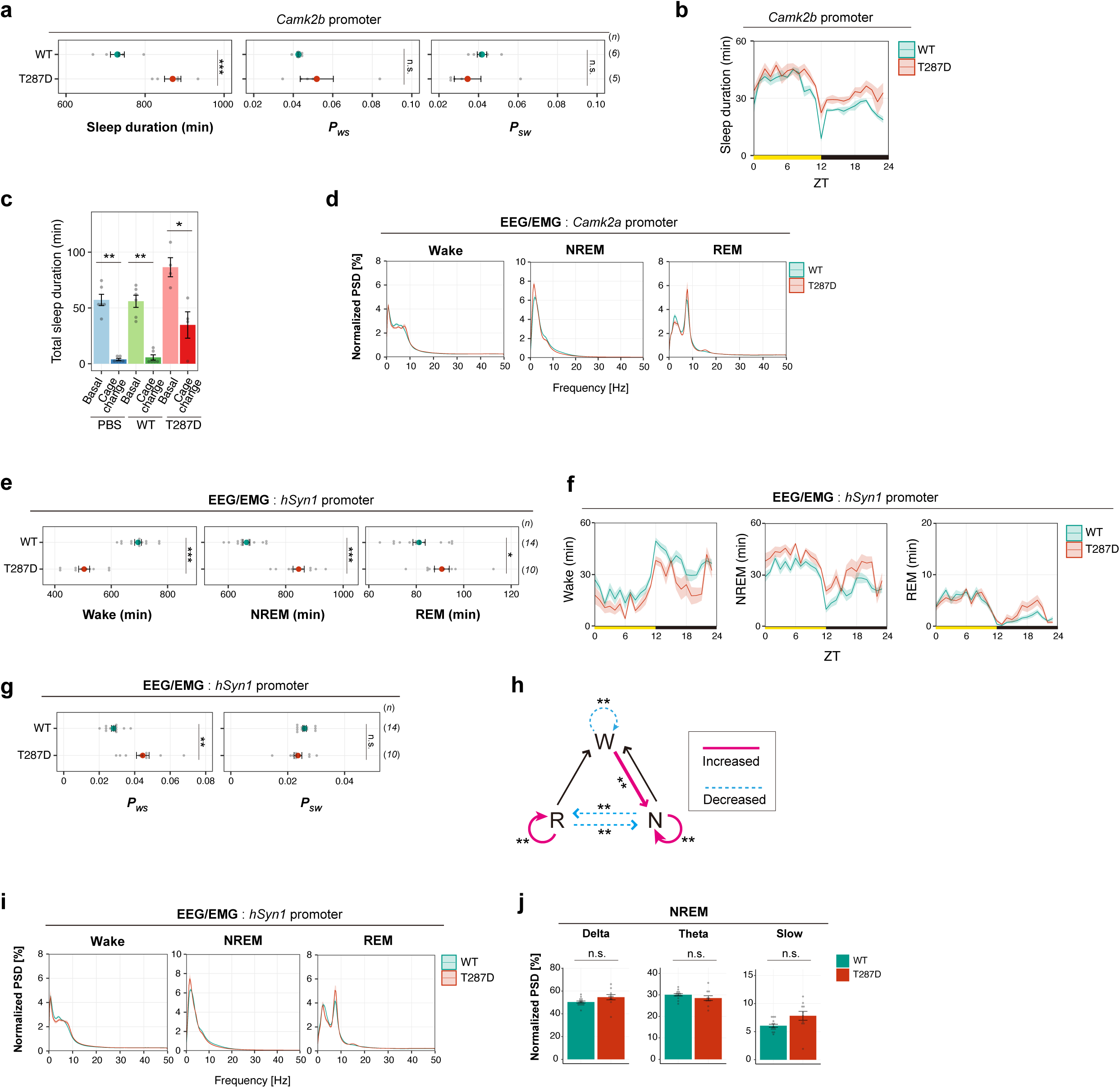
Phosphorylation of CaMKIIβ regulates NREM sleep induction and sleep needs. **(a-b)** Sleep/wake parameters **(a)** and sleep profiles **(b)**, averaged over six days, of mice expressing WT CaMKIIβ or the T287D mutant (T287D) under the *Camk2b* promoter. **(c)** Total sleep duration from ZT0 to ZT2 of mice expressing wild-type CaMKIIβ (WT, n = 6) and the CaMKIIβ T287D mutants (T287D, n = 4) after cage change at ZT0. PBS: PBS-injected control mice (n = 6). “Basal” represents the sleep duration from ZT0 to ZT2 averaged over three days before the day of the cage change. **(d)** EEG power spectra of mice expressing WT CaMKIIβ or the T287D mutant under the *Camk2a* promoter. **(e-g)** Sleep parameters **(e and g)** and sleep profiles **(f)** measured by EEG/EMG recordings for mice expressing CaMKIIβ WT or the T287D mutant under the *hSyn1* promoter. **(h)** Differences in transition probabilities (between wakefulness (W), NREM sleep (N), and REM sleep (R)) between WT CaMKIIβ or T287D-expressing mice under the *hSyn1* promoter. Magenta lines and dashed blue lines indicate when the values for the T287D-expressing mice are significantly (p < 0.05) higher and lower, respectively. **(i-j)** EEG power spectra **(i)** and NREM power density in typical frequency domains **(j)** of mice expressing WT CaMKIIβ or the T287D mutant under the *hSyn1* promoter. Error bars: SEM, *p < 0.05, **p < 0.01, ***p < 0.001, n.s.: no significance. **Figure 2-figure supplement 1-source data 1** Source data for Figure 2-figure supplement 1a-j

**Figure 4-figure supplement 1.**
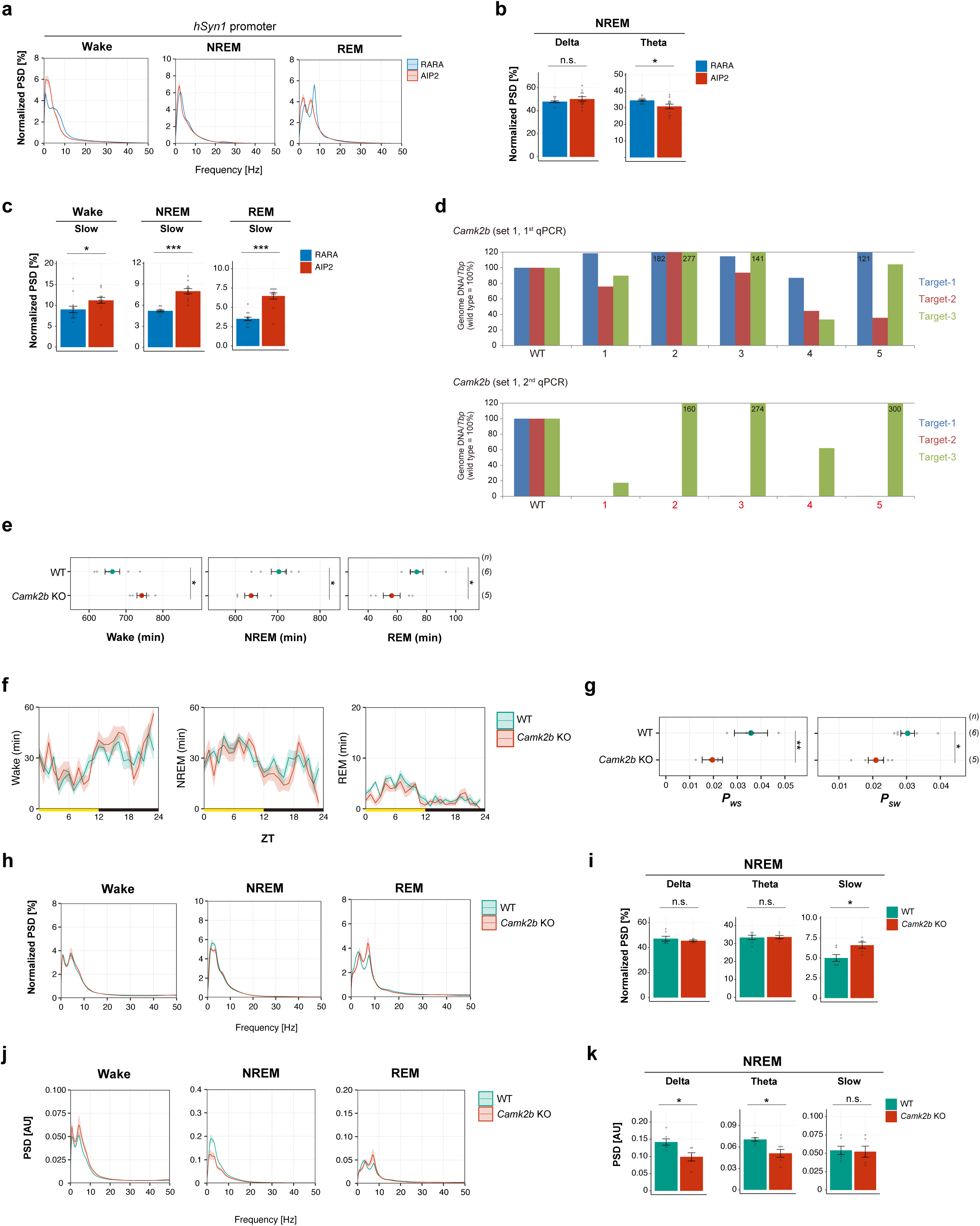
Sleep phenotypes of AIP2-exressing mice and *Camk2b* KO mice. **(a-b)** EEG power spectra **(a)** and NREM power density in delta and theta domains **(b)** of mice expressing AIP2 or the RARA mutant. **(c)** Power densities in slow domains of mice expressing AIP2 or the RARA mutant. **(d)** The genotype of *Camk2b* KO mice. The relative amount of intact DNA for each target sequence was normalized to be 100% for wild-type mouse (WT). The qPCR was performed with two independent primer sets (1^st^ and 2^nd^) for the three target sites. When the 0.5% criteria were met in either set, the mouse was considered a KO mouse. All the mice (n=5) were confirmed as KO mice by the 2^nd^ qPCR. **(e-g)** Sleep parameters **(e and g)** and sleep profiles **(f)** measured by EEG/EMG recordings for *Camk2b* KO mice and wild-type C57BL/6N mice (WT). **(h-i)** Normalized EEG power spectra **(h)** and NREM power density in typical frequency domains **(i)** of *Camk2b* KO mice and wild-type C57BL/6N mice (WT). EEG Power was normalized relative to the total power in each frequency band. **(j-k)** EEG power spectra **(j)** and NREM power density in typical frequency domains **(k)** of *Camk2b* KO mice and wild-type C57BL/6N mice (WT) without normalization. Error bars: SEM, *p < 0.05, **p < 0.01, ***p < 0.001, n.s.: no significance. **Figure 4-figure supplement 1-source data 1** Source data for Figure 4-figure supplement 1a, b, c, d, e, f, g, h, i, j, k

**Figure 5-figure supplement 1.**
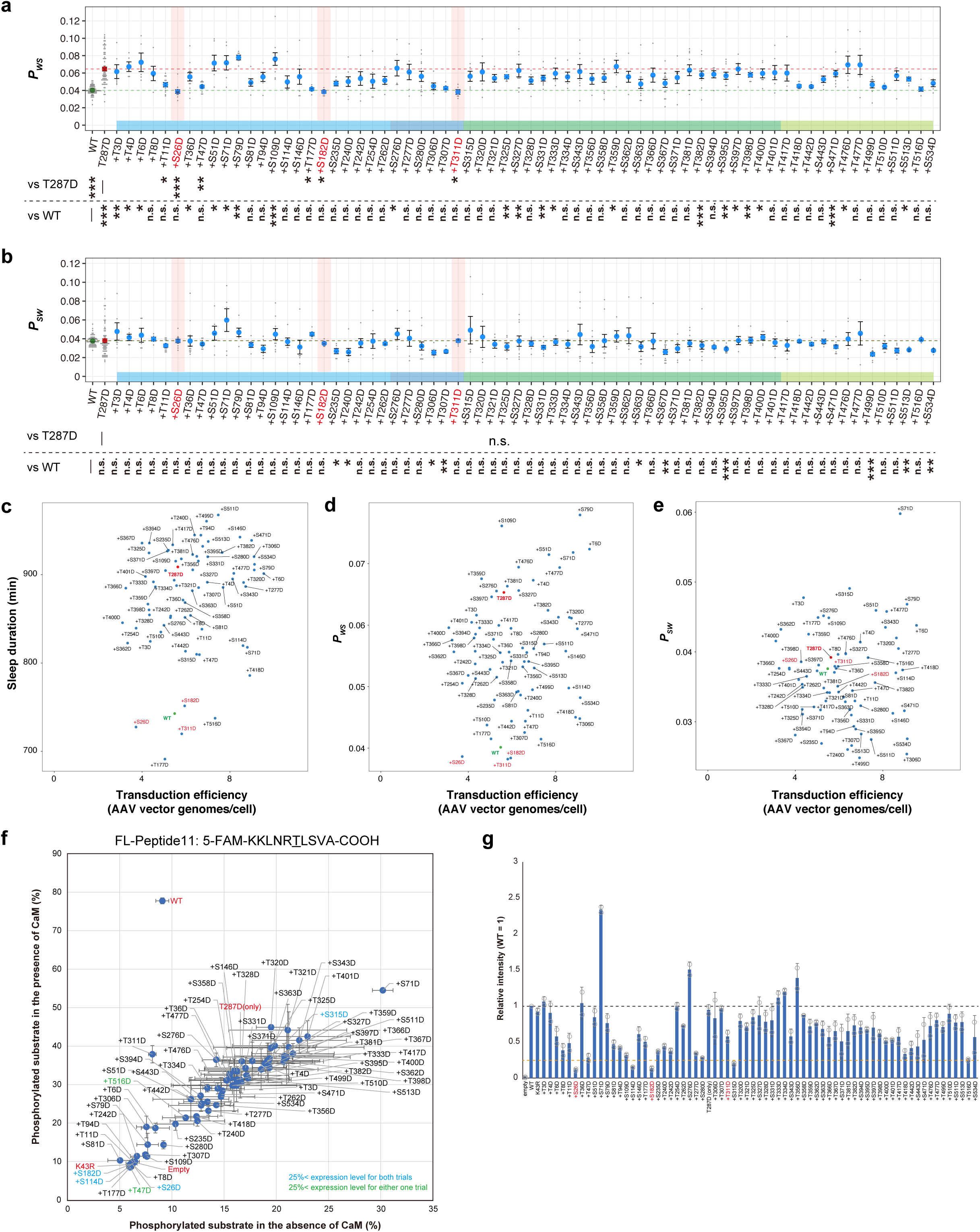
Multi-site phosphorylation of CaMKIIβ can cancel sleep induction. **(a-b)** Daily *P_WS_* **(a)** and *P_SW_* **(b)**, averaged over six days, of the mice expressing CaMKIIβ double-phopshomimetic mutants (n = 5–12) shown in Figure 5a. Dashed green and red lines represent the averaged sleep duration of wild-type CaMKIIβ-expressing mice (WT, n = 71) and CaMKIIβ T287D mutants-expressing mice (T287D, n = 68), respectively. The plus sign in a mutant name indicates a combination with T287D. Multiple comparison test was performed against WT (vs WT) or T287D (vs T287). In the comparison with the T287D mutant, “n.s.” labels are omitted for visibility in **(a)**. **(c-e)** Calculated transduction efficiency plotted against sleep duration **(c)**, *P_WS_* **(d)** and *P_SW_* **(e)**. Transduction efficiency is an estimation of the number of AAV vector genomes present per cell in a mouse brain. After the SSS measurements, we purified the AAV vector genomes from the mice brains and then quantified them with a WPRE-specific primer. The transduction efficiency is the quantification value normalized to a TATA-binding protein (*Tbp*)-specific primer using qPCR. **(f-g)** *In vitro* kinase activity and expression level of CaMKIIβ double-phosphomimetic mutants. The data represented as in **Figure 1-figure supplement 2f** and **g.** Error bars except in **(f-g)**: SEM, *p < 0.05, **p < 0.01, ***p < 0.001, n.s.: no significance. **Figure 5-figure supplement 1-source data 1** Source data for Figure 5-figure supplement 1a-g.

**Figure 6-figure supplement 1.**
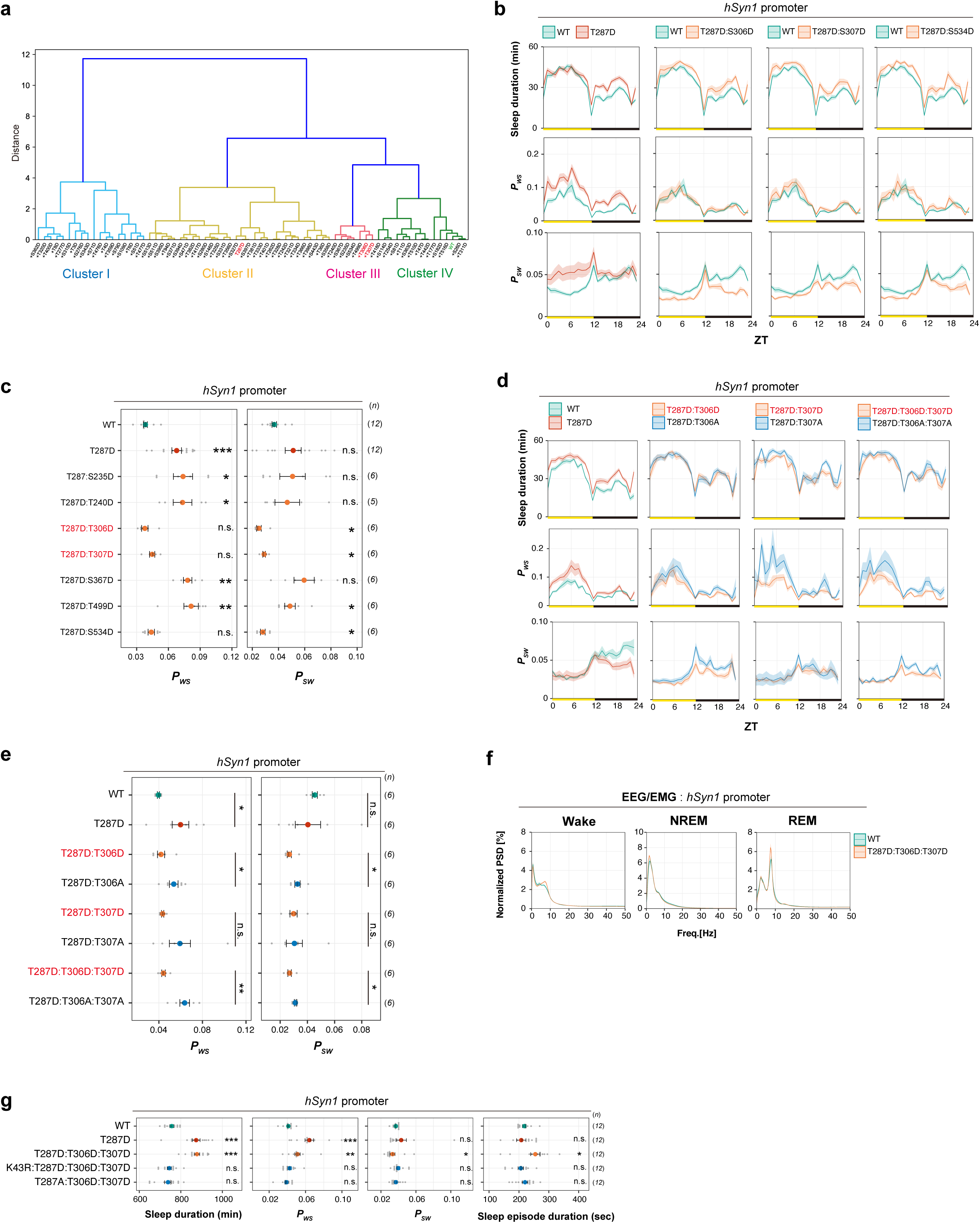
Multi-site phosphorylation of CaMKIIβ regulates sleep stabilization. **(a)** Hierarchical clustering dendrogram from sleep profiles of mice expressing the CaMKIIβ double-phopshomimetic mutants shown in Figure 6a. Cluster I contains mutants that increase *P_SW_*, (sleep destabilizing). Cluster II, the largest cluster, contains mutants that increase *P_WS_*, similar to the T287D mutant. Cluster III contains mutants that decrease *P_SW_* (sleep stabilizing). Cluster IV contains mutants with properties similar to WT. The “T287D-canceling” mutants such as +S26D, +S182D, and +T311D belong to cluster IV. The vertical branch length represents the degree of dissimilarity in sleep profiles among the mutants. Branch colors indicate clusters. **(b-c)** Profiles of sleep and transition probability **(b)** and *P_WS_* and *P_SW_* **(c)**, averaged over six days, of mice expressing the double-phopshomimetic mutants shown in **Figure 6b and 6c.** The shaded areas represent SEM. Multiple comparison test was performed against wild-type CaMKIIβ-expressing mice (WT). **(d-e)** Profiles of sleep and transition probability **(d)** and *P_WS_* and *P_SW_* **(e)**, averaged over six days, of mice expressing the CaMKIIβ mutants with sleep-stabilizing residues substituted with D or A shown in **Figure 6d and 6e**. The shaded areas represent SEM. Multiple comparison test was performed against wild-type CaMKIIβ-expressing mice (WT). **(f)** EEG power spectra of mice expressing WT CaMKIIβ and the T287D:T306D:T307D mutant shown in Figure 6f-j. **(g)** Sleep/wake parameters, averaged over six days, of mice expressing the CaMKIIβ T287D:T306D:T307D mutant with the K43R or T287A mutation. Multiple comparison test was performed against wild-type CaMKIIβ-expressing mice (WT). Error bars: SEM, *p < 0.05, **p < 0.01, ***p < 0.001, n.s.: no significance. **Figure 6-figure supplement 1-source data 1** Source data for Figure 6-figure supplement 1a-g

**Figure 6-figure supplement 2.**
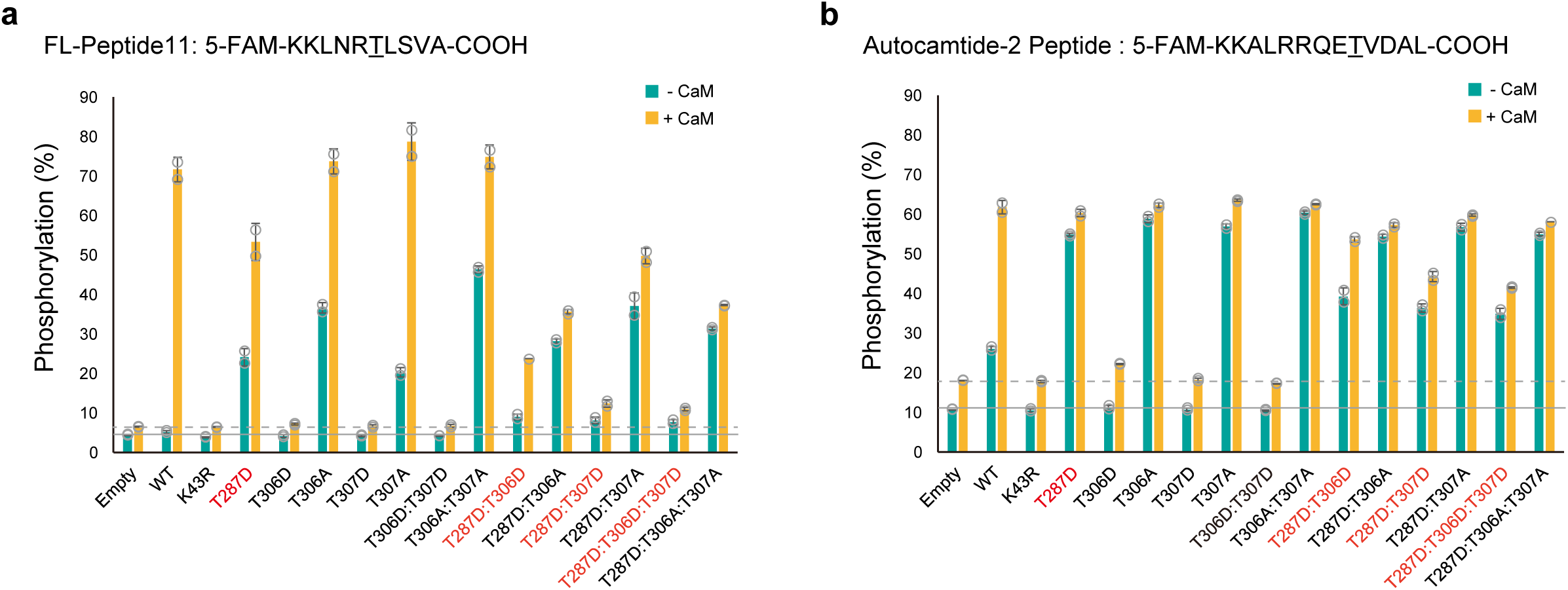
Ordered auto-phosphorylation of sleep-controlling residues in CaMKIIβ. **(a-b)** *In vitro* kinase activity of CaMKIIβ mutants against FL-Peptide 11 **(a)** or autocamtide-2 peptide **(b)** in the absence (-CaM, green bar) or presence (+CaM, yellow bar) of CaM. The amino acids phosphorylated by CaMKIIβ are underlined in the peptide sequences. Phosphorylation (%) indicates the percentage of the phosphorylated substrate relative to the total peptide. The reported values are the mean ± SD (n = 2 independent experiments). The dashed and solid lines indicate background signals measured in cell lysate transfected with the control empty vector (Empty). **Figure 6-figure supplement 2-source data 1** Source data for Figure 6-figure supplement 2a-b.

**Figure 7-figure supplement 1.**
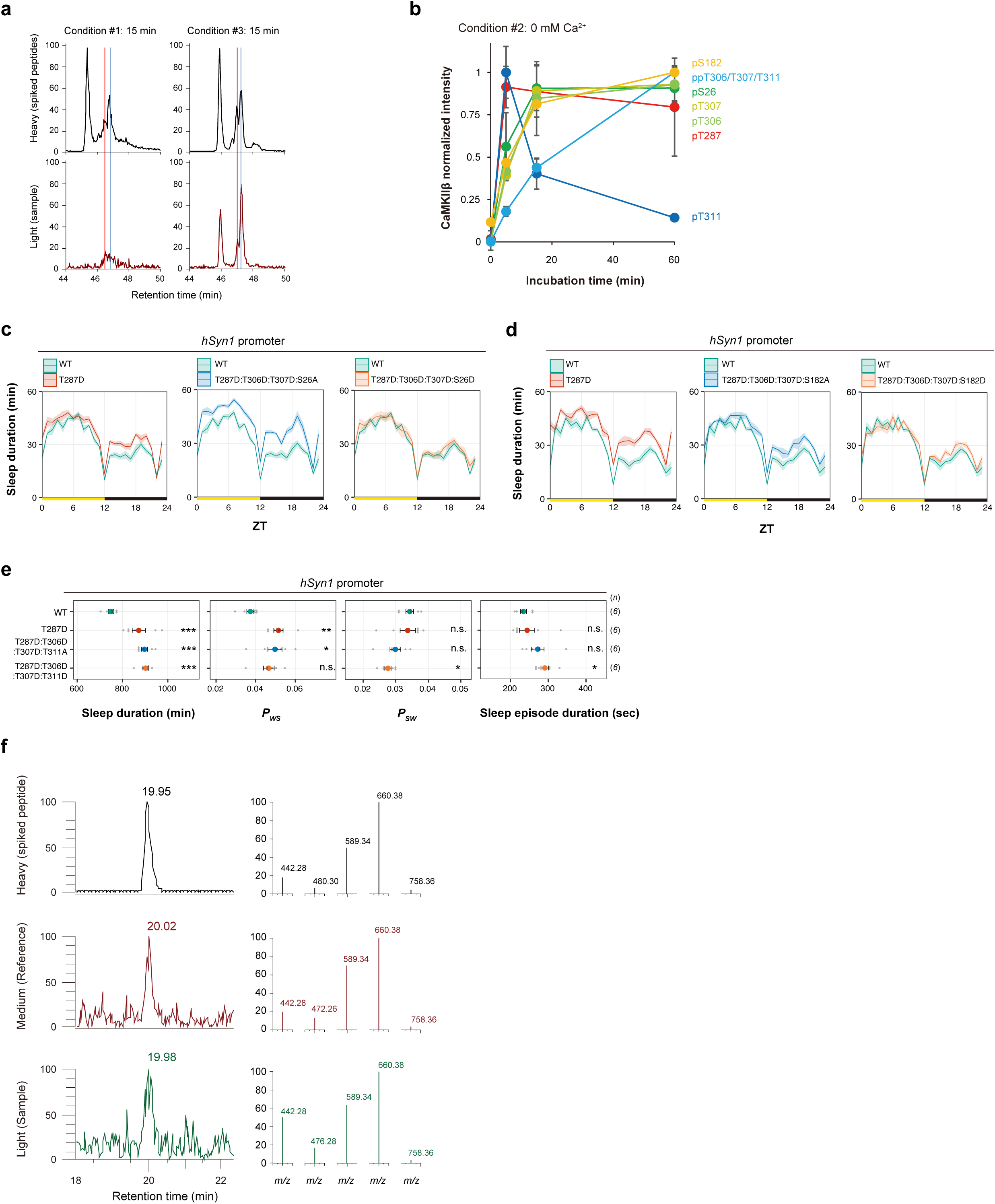
Ordered multi-site phosphorylation of CaMKIIβ underlies multi-step sleep regulation. **(a)** Example chromatogram of SRM measurement for pS182 peptide. Red line indicates the retention time of a peptide phosphorylated at S182. Red line indicates the retention time of a peptide phosphorylated at T177. **(b)** Phosphorylation time course for sleep-controlling residues *in vitro*. Time series changes of the residues in the condition #2 are extracted and overlaid from Figure 7a. The values are shown as relatives, with the maximum value of each residue in the time course as 1. **(c)** Profiles of sleep, averaged over six days, of mice expressing the quadruple-phosphomimetic CaMKIIβ mutants related to S26 shown in Figure 7b. The shaded areas represent SEM. **(d)** Profiles of sleep, averaged over six days, of mice expressing the quadruple-phosphomimetic CaMKIIβ mutants related to S182 shown in Figure 7c. The shaded areas represent SEM. **(e)** Sleep/wake parameters of mice expressing the quadruple-phosphomimetic CaMKIIβ mutants related to T311. Multiple comparison test was performed against wild-type CaMKIIβ-expressing mice (WT). **(f)** Phosphorylation of S26 (CaMKIIβ) or S25 (CaMKIIα) residue in mice brain. Mice brain samples shown in **Figure 1g and 1h** were also subjected to SRM analysis with mass-spectrometry method for analyzing the phosphorylation of S26 (CaMKIIβ) or S25 (CaMKIIα) residues. Representative chromatograms shown in left indicated that a synthesized and heavy-labeled phosphorylated peptide, of which sequence is identical to a trypsin-digested peptide sequence corresponding to S26 (CaMKIIβ) or S25 (CaMKIIα) was detected at retention time ∼20 min. Medium-labeled peptide sample (derived from internal control mixture) and light-labeled peptide sample (derived from individual samples) also showed a peak at retention time ∼20 min. The product ion spectrum on the right shows that each product ion from the five different transitions in the three samples has a similar intensity distribution. These results suggest that a peptide corresponding to phosphorylated S26 (CaMKIIβ) or S25 (CaMKIIα) was included in trypsin-digested mice brain samples. Error bars: SEM, *p < 0.05, **p < 0.01, ***p < 0.001, n.s.: no significance. **Figure 7-figure supplement 1-source data 1** Source data for Figure 7-figure supplement 1c-e **Figure 7-source data 1 and 2** These source data include source data for Figure 7-figure supplement 1a, b, f.

## MATERIALS and METHODS

### Plasmids

Mouse *Camk2b* cDNA (NM_007595) was subcloned into the pMU2 vector ^71^ that expresses genes under the CMV promoter. Note that the FLAG-tag involved in the original pMU2 vector was removed in the construct used in this study. Mutagenesis of pMU2-*Camk2b* was conducted by inverse PCR with Mighty Cloning Reagent Set (Blunt End) (Takara Bio, Japan) following to the manufacturer’s protocol.

For pAAV construction, the *Camk2b* sequence was transferred into the pAAV vector (kindly provided by Dr. Hirokazu Hirai) along with the *hSyn1* promoter ^72^, FLAG tag, *Camk2b* 3’UTR, WPRE, and SV40 polyA sequences as illustrated in Figure 1b. For the *Camk2b* 3’UTR used in this study, the evolutionarily conserved ∼350 bp (chr11:5,971,489-5,971,827, GRCm38/mm10) and ∼650 bp (chr11:5,969,672-5,970,313, GRCm38/mm10) regions in the mouse *Camk2b* 3’UTR was cloned and assembled tandemly. For double-floxed inverted open reading frame (DIO) constructs, the inverted FLAG-*Camk2b* sequence flanked by lox2272 and loxP was inserted between the *hSyn1* promoter and the *Camk2b* 3’UTR of the pAAV vector as illustrated in **Figure 3a**. For more targeted gene expression, the *hSyn1* promoter was replaced with other promoters (Figure 2 and Figure 1-figure supplement 2a). A vector containing *Camk2a* promoter sequence was a kind gift from Drs. Masamitsu Iino and Yohei Okubo (The University of Tokyo). For the *Camk2b* promoter, ∼1300 bp region (chr11:6,065,706-6,066,972, GRCm38/mm10) upstream of the TSS of the *Camk2b* gene was cloned using a pair of primers (5’-AGCACTCTGTCAAATGTACCTTTAG-3’; 5’-AGATCTGCTCGCTCTGTCCC-3’).

The mCherry-AIP2 was constructed by fusing the AIP2 sequence (KKKLRRQEAFDAL) to the C-terminus of mCherry via a (GGGGS)x3 linker. To construct pAAV, the mCherry-AIP2 sequences were inserted into the pAAV vector with the *hSyn1* promoter, dendritic targeting element (DTE) of mouse *Map2* gene, WPRE, and SV40 polyA sequences. The DTE of *Map2* were amplified and cloned from C57BL/6N mouse genomic DNA ^73^.

pUCmini-iCAP-PHP.eB for PHP.eB production was a gift from Dr. Viviana Gradinaru (Addgene plasmid # 103005).

### Animals and sleep phenotyping

All experimental procedures and housing conditions were approved by the Institutional Animal Care and Use Committee of RIKEN Center for Biosystems Dynamics Research and the University of Tokyo. All the animals were cared for and treated humanely in accordance with the Institutional Guidelines for Experiments using Animals. All mice had *ad libitum* access to food and water, and were maintained at ambient temperature and humidity conditions under a 12 h light/dark cycle. All C57BL/6N mice were purchased from CLEA Japan (Tokyo, Japan). The mice used in each experiment were randomly chosen from colonies. EEG/EMG recording for the *Camk2b* KO mice (**Figure 4-figure supplement 1**) were conducted at the University of Tokyo. Other animal experiments were performed in the RIKEN Center for Biosystems Dynamics Research.

### Mass spectrometry and western blotting of mice brain samples

C57BL/6N mice (CLEA Japan, Japan) were housed in a light-dark controlling rack (Nippon Medical & Chemical instruments, Japan) and habituated to a 12 h light/dark cycle for at least one week. At eight weeks old, half of the mice were subjected to the sleep deprivation protocol from ZT0 to ZT6. The sleep deprivation was conducted by gentle handling and cage changing ^42^ at every 2 h. The other mice were housed under *ad lib* sleep conditions. At ZT6, the mice were sacrificed by cervical dislocation and their forebrain was immediately frozen in liquid nitrogen. The brain samples were stored at −80°C. The frozen brains were cryo-crushed with a Coolmil (Tokken, Japan) pre-cooled in liquid nitrogen, and the brain powders were stored at −80°C.

The brain powders were then lysed and digested according to the phase-transfer surfactant (PTS) method ^74^. Approximately 10 mg of brain powder was added to the 500 μl of Solution B (12 mM sodium deoxycholate, 12 mM N-lauroylsarcosine sodium salt, 50 mM ammonium hydrogen carbonate) containing phosphatase inhibitors (1 mM sodium orthovanadate, 1 mM β-glycerophosphoric acid disodium salt pentahydrate, 4 mM sodium (+)-tartrate dihydrate, 2.5 mM sodium fluoride, 1.15 mM disodium molybdate (VI) dihydrate) pre-heated at 98 °C and sonicated extensively. After further incubation at 98 °C for 30 min, the samples were reduced with 10 mM dithiothreitol (FUJIFILM Wako Pure Chemical, Japan) at room temperature for 30 min, and then alkylated with 100 mM iodoacetamide (Sigma-Aldrich, U.S.A.) at room temperature for 30 min. The samples were then diluted to five-fold by adding Solution A (50 mM ammonium hydrogen carbonate) and digested them by adding 5 μg of lysyl endopeptidase (Lys-C) (FUJIFILM Wako Pure Chemical, Japan). After 37°C overnight incubation, 5 μg of trypsin (Roche, Switzerland) was added and the mixture was further incubated at 37°C overnight. After the digestion, an equal volume of ethyl acetate was added to the sample, which was acidified with 0.5% TFA and well mixed to transfer the detergents to the organic phase. The sample was then centrifuged at 2,380 x g for 15 min at room temperature, and an aqueous phase containing peptides was collected and dried with a SpeedVac (Thermo Fisher Scientific, U.S.A.).

The dried peptides were solubilized in 1 mL of 2% acetonitrile and 0.1% TFA. We prepared an internal control by mixing 500 μL of each peptide solution. The individual samples were the remaining 500 μL of each peptide solution. The internal control and individual samples were trapped and desalted on a Sep-Pak C18 cartridge (Waters, U.S.A.). Dimethyl-labeling was then applied to the peptides on the cartridge as previously described ^75^. Formaldehyde (CH_2_O, Nacalai Tesque, Japan) and NaBH_3_CN (Sigma-Aldrich, U.S.A.) were added to the individual samples (light label), and isotope-labeled formaldehyde (CD_2_O, Cambridge Isotope Laboratories, U.S.A.) and NaBH_3_CN (Sigma-Aldrich, U.S.A.) were added to the internal control mixture (medium label). The dimethyl-labeled peptides on the Sep-Pak cartridge were eluted with an 80% acetonitrile and 0.1% TFA solution. Then, equal amount of medium-labeled internal control mixture was added to each light-labeled individual sample. This allowed us to compare the relative amount of peptides in the individual samples with each other using the equally-added medium-labeled internal control mixture as a standard.

A one-hundredth of the mixture underwent LC-MS analysis to quantify the amount of CaMKIIα/β and total proteins. The remaining mixture was applied to High-Select™ Fe-NTA Phosphopeptide Enrichment Kit (Thermo Fisher Scientific, U.S.A.) to enrich the phosphorylated peptides following the manufacture’s protocol.

All analytical samples were dried with a SpeedVac (Thermo Fisher Scientific, U.S.A.) and dissolved in 2% acetonitrile and 0.1% TFA. Mass-spectrometry-based quantification of CaMKIIα/β-derived peptides was carried out by selected reaction monitoring (SRM) analysis using a TSQ Quantiva triple-stage quadrupole mass spectrometer (Thermo Fisher Scientific, U.S.A.). The following parameters were selected: positive mode, Q1 and Q3 resolutions of 0.7 full width of half maximum (FWHM), cycle time of 2 s, and gas pressure of 1.5 Torr. The mass spectrometer was equipped with an UltiMate 3000 RSLCnano nano-high performance liquid chromatography (HPLC) system (Thermo Fisher Scientific, U.S.A.), and a PepMap HPLC trap column (C18, 5 µm, 100 A; Thermo Fisher Scientific, U.S.A.) for loading samples. Samples were separated by reverse-phase chromatography using a PepMap rapid separation liquid chromatography (RSLC) EASY-Spray column (C18, 3 µm, 100 A, 75 µm x 15 cm; Thermo Fisher Scientific, U.S.A.) using mobile phases A (0.1% formic acid/H2O) and B (0.1% formic acid and 100% acetonitrile) at a flow rate of 300 nl/min (4% B for 5 min, 4%–35% B in 55 min, 35%–95% B in 1 min, 95% B for 10 min, 95%–4% B in 0.1 min and 4% B for 9.9 min). The eluted material was directly electro-sprayed into the MS. The SRM transitions of the target peptides were determined based on the pre-analysis of several samples including mice brains, 293T cells expressing CaMKIIβ, and synthesized peptides, and optimized using Pinpoint software, version 1.3 (Thermo Fisher Scientific, U.S.A.). The Quan Browser of the Quan Browser data system, version 3.0.63 (Thermo Fisher Scientific, U.S.A.) was used for data processing and quantification.

To estimate the relative amount of total peptides involved in each brain sample, approximately half of the light/medium mixture sample without the enrichment of phosphopeptides was analyzed by data-dependent MS/MS with a mass spectrometer (Q-Exactive Mass Spectrometer, Thermo Fisher Scientific, U.S.A.) equipped with an HPLC system containing nano HPLC equipment (Advance UHPLC, Bruker Daltonics, U.S.A.) and an HTC-PAL autosampler (CTC Analytics, Switzerland) with a trap column (0.3 x 5 mm, L-column, ODS, Chemicals Evaluation and Research Institute, Japan). An analytical sample were loaded into the LC-MS system to be separated by a gradient using mobile phases A (0.1% formic acid) and B (0.1% formic acid and 100% acetonitrile) at a flow late 300 nL/min (4% to 32% B in 190 min, 32% to 95% B in 1 min, 95% B for 2 min, 95% to 4% B in 1 min and 2% B for 6 min) with a homemade capillary column (200 mm length, 100 μm inner diameter) packed with 2 μm C18 resin (L-column2, Chemicals Evaluation and Research Institute, Japan). The eluted peptides were then electrosprayed (1.8-2.3 kV) and introduced into the MS equipment (positive ion mode, data-dependent MS/MS). MS data were analyzed by Proteome Discoverer version 2.2 (Thermo Fisher Scientific, U.S.A.) with the Swiss-Prot section of UniProtKB mouse database (as of August 9^th^, 2018). The relative amount of CaMKIIα/β protein was normalized to the median of all quantified proteins for each sample, with the effect derived from different amounts of start materials being excluded.

For the western-blotting analysis, brain powder was lysed in the 3x Laemmli sample buffer (20% glycerol, 2.25% sodium dodecyl sulphate (SDS), 187.5 mM Tris-HCl at pH 6.8, 0.015% bromophenol blue) pre-heated at 98 °C and sonicated extensively. Approximately 0.1 mg of brain powder (∼ 10 µg protein) was subjected to each lane of hand-made polyacrylamide gel. The samples were separated by SDS–polyacrylamide gel electrophoresis and then transferred to a polyvinylidene difluoride (PVDF) membrane (Hybond-P PVDF membranes, Merck, Germany) by a wet-transfer apparatus (Vep-3, Thermo Fisher Scientific, U.S.A.). The membrane was washed by TTBS (0.9% NaCl, 0.1% Tween-20, and 100 mM Tris-HCl at pH 7.5) and non-specific protein binding was blocked by incubating with Blocking One solution (Nacalai Tesque, Japan) for 1 hr at room temperature. FLAG-tagged protein was detected by an anti-FLAG M2 antibody conjugated with horseradish peroxidase (A8592, Sigma-Aldrich, U.S.A.). CaMKIIβ and α-Tublin were detected by primary antibodies anti-CaMK2β (#139800, Thermo Fisher Scientific, U.S.A.) or anti-alpha Tubulin [DM1A] (ab7291, Abcam), respectively, followed by the incubation with a secondary antibody anti-mouse IgG HRP conjugate (W4021, Promega, U.S.A.). All the primary antibodies were diluted 1/3000 in 10% Blocking One/TTBS (50 mM Tris, 0.5 M NaCl, 0.05% Tween-20, pH 7.4) and incubated with the membrane for overnight at 4 °C. The secondary antibody was diluted 1/2000 in 10% Blocking One/TTBS (50 mM Tris, 0.5 M NaCl, 0.05% Tween-20, pH 7.4) and incubated with the membrane for 1 hr at room temperature. Immunoreactivities were detected with Clarity Western ECL Substrate for Chemiluminescent Western Blot Detection (Bio-rad, U.S.A.) and ChemiDoc XRS+ system (Bio-rad, U.S.A.).

### Tissue clearing and LSFM imaging

AAV-administrated mice were perfusion-fixed under anesthesia, and brains were isolated. Isolated brains were fixed overnight in 4% PFA and then washed with PBS. For clearing the mouse brain, second-generation CUBIC protocols were used. The detailed protocol can be found in a previous report ^76^. For delipidation, the brain was treated with CUBIC-L (10% (w/w) N-butyldiethanolamine and 10% (w/w) Triton X-100) solution at 37°C for 5 days. For nuclear staining, the brain was rinsed with PBS and incubated in 1:250 diluted RedDot2 (Biotium, 40061) in staining buffer (10% (w/w) Triton X-100, 10% (w/w) Urea, 5% (w/w) *N,N,N’,N’*-Tetrakis(2-hydroxypropyl)ethylenediamine, 500mM NaCl) for 3 days at 37°C. The stained brain sample was washed with PBS and then treated in CUBIC-R+ solution (45% (w/w) antipyrine, 30% (w/w) nicotinamide, 0.5% (v/v) *N*-butyldiethanolamine) for 3 days at 25°C for RI matching. For whole-brain imaging, the cleared brain sample was embedded in a CUBIC-R+ gel, which contains 2% (w/w) agarose in the CUBIC-R+ solution and set in a customized light-sheet microscopy (LSFM) ^76^. Dual-colored images were simultaneously acquired with illumination objective lens (MVPLAPO 1×, Olympus, Japan), 10× detection objective lens (XLPLN10XSVMP, Olympus, Japan), Dichroic mirror (DMSP650L, Thorlabs, U.S.A.) and following laser and fluorescence filters: RedDot2 [Ex: 594 nm, Em: 700 nm bandpass (FB700-40, Thorlabs, U.S.A.)], mCherry [Ex: 594 nm, Em: 625 nm bandpass (ET625/30m, Chroma Technology, U.S.A.)]. Stacked brain images were reconstructed and visualized by the Imaris software (Bitplane).

### CaMKIIβ kinase assay

293T cells were grown in culture medium consisting of Dulbecco’s Modified Eagle Medium (DMEM) (high glucose, Thermo Fisher Scientific, U.S.A.), 10% FBS (Sigma-Aldrich) and 100 U/ml penicillin-streptomycin (Thermo Fisher Scientific, U.S.A.) at 37°C with 5% CO_2_. The cells were plated at 2 × 10^4^ cells per well in 24-well plates 24 h before transfection. The cells in each well were transfected with 1.6 μg PEI (Polyethylenimine, Linear, MW 25000, Polysciences, U.S.A.) and 400 ng of pMU2-*Camk2b* plasmids. 24 hr after the transfection, the medium in each well was replaced with flesh culture medium. The cells were stayed for another 48 h, and collected by removing all the culture medium. The remaining cells on the 24-well plate were stored at −80 °C.

The cells were lysed with 200 µl of cell lysis buffer (50 mM HEPES-NaOH pH 7.6, 150 mM NaCl, 0.5 mM CaCl_2_, 1 mM MgCl_2_, and 0.25% (v/v) NP-40) containing protease inhibitors (100 mM phenylmethanesulfonyl fluoride, 0.1 mM Aprotinin, 2 mM Leupeptin hemisulfate, 1 mM Pepstatin A, and 5 mM Bestatin). Followed by extensive sonication, the cell lysates were collected and stored at −80 °C.

The relative expression levels of CaMKIIβ in each cell lysate was estimated by dot blot. A PVDF membrane (Hybond-P PVDF membranes, Merck, Germany) was immersed in 100% methanol (Nacalai Tesque, Japan) and then soaked in water for at least 10 min. Excess water was removed from the membrane, 2 µl of four-fold diluted cell lysate was spotted on the membrane. The membrane was then dried completely, immersed in 100% Methanol and equilibrated in water. The membrane was incubated in Blocking One solution (Nacalai Tesque, Japan) for 1 hr at room temperature. After the blocking reaction, the membrane was incubated for 2 hr with the primary antibody anti-CaMK2β (#139800, Thermo Fisher Scientific, U.S.A.) diluted at 1/3000 in 10% Blocking One/TTBS (50 mM Tris, 0.5 M NaCl, 0.05% Tween-20, pH 7.4). The membrane was washed with TTBS, and incubated for 1 h with the secondary antibody anti-mouse IgG HRP conjugate (W4021, Promega, U.S.A.) diluted at 1/3000 in 10% Blocking One/TTBS. Immunoreactivities of the blotted proteins were detected with Clarity Western ECL Substrate for Chemiluminescent Western Blot Detection (Bio-rad, U.S.A.) and ChemiDoc XRS+ system (Bio-rad, U.S.A.). The images were analyzed with Image Lab software (version 6.01, Bio-rad, U.S.A.). For each dot-blot experiment, serial dilution of cell lysate expressing the WT CaMKIIβ was spotted to confirm that the quantification of the dot blot signal was within the linear range of detection.

The kinase activity of CaMKIIβ-expressed cell lysate was calibrated as follows. First, a serial dilution of cell lysate expressing WT CaMKIIβ was prepared. Then 5 µl of each diluted cell lysate was mixed with 15 µl of cell lysis buffer containing 0.33 mM ATP and 5 µM ProfilerPro Kinase Peptide Substrate 11 5-FAM-KKLNRTLSVA-COOH (PerkinElmer, U.S.A.) in the presence or absence of 0.66 µM CaM (Sigma-Aldrich, U.S.A.). After incubating at 37°C for 10 min, and the reaction was stopped by incubating at 98°C for 10 min. 100 µL of 2% ACN/0.1% TFA was added to the reaction mixture and the mixture was analyzed by mobility shift assay (LabChip EZ Reader II; PerkinElmer, U.S.A.). The kinase activity is the percentage of phosphorylated peptide signal over the total substrate peptide signal. Based on the kinase activity obtained from the serial dilution of cell lysate, we determined two critical dilution ratios. One is a dilution rate that gives the ∼50% kinase activity in the calibration curve in the presence of CaM (called “half-max dilution rate”). The other dilution rate (called “background dilution rate”) is based on the calibration curve in the absence of CaM, where most of the kinase activity should come from the endogenous proteins in 293T cells. We determined the “background dilution rate” to give the phosphorylation rate around 10% or less in the absence of CaM.

With these two critical dilution rates, we normalized the relative expression levels of each WT or mutant CaMKIIβ. First, all the cell lysates were diluted to the “background dilution rate” in a cell lysis buffer. We also prepared a lysate of the cells treated with the PEI transfection procedure without vector plasmid (called PEI-treated cell lysate) and diluted it to the same “background dilution rate.” Next, the diluted cell lysate expressing the WT CaMKIIβ was further diluted to reach the final dilution rate (equivalent to the “half-max dilution rate”) by mixing with diluted PEI-treated cell lysate. For the CaMKIIβ mutants (except those with 25% or lower expression levels compared to WT CaMKIIβ), the mixing ratio between CaMKIIβ-expressed lysate and PEI-treated lysate were adjusted based on the relative expression level of CaMKIIβ mutants quantified by dot blot. Through these processes, we obtained a series of diluted cell lysates with the same background kinase activity level and the same relative expression levels of WT or mutant CaMKIIβ.The kinase activity of WT CaMKIIβ is expected to be around 50%.

The quantification of kinase activity was carried out by mixing 5 μl of cell lysates (diluted as described above) and 15 µl of cell lysis buffer containing 0.33 mM (**Figure 1-figure supplement 3**) or 3.3 mM (**Figure 5-figure supplement 1** and **8**) ATP and 5 µM ProfilerPro Kinase Peptide Substrate 11 5-FAM-KKLNRTLSVA-COOH (PerkinElmer, U.S.A.), in the presence or absence of 0.66 µM CaM (Sigma-Aldrich, U.S.A.). FAM-labeled autocamtide-2 5-FAM-KKALRRQETVDAL-COOH, synthesized with a peptide synthesizer Syro Wave (Biotage, Sweden) using Fmoc solid-phase chemistry, was used in the experiment shown in Figure 6j. After incubating at 37°C for 10 min, and the reaction was stopped by incubating at 98°C for 10 min. 100 µL of 2% ACN/0.1% TFA was added to the reaction mixture and the mixture was analyzed by mobility shift assay (LabChip EZ Reader II and operation software version 2.2.126.0; PerkinElmer, U.S.A.).

### Mass spectrometry of purified CaMKIIβ

The spike peptides were synthesized with a peptide synthesizer Syro Wave (Biotage, Sweden) using Fmoc solid-phase chemistry. The synthesized peptides were treated with dithiothreitol and iodoacetamide as described above. The peptides were desalted by using hand-made C18 StageTips ^77^. The desalted peptides on the StageTips were subjected to dimethyl-labeling with isotope-labeled formaldehyde (^13^CD_2_O, ISOTEC, U.S.A.) and NaBD_3_CN (Cambridge Isotope Laboratories, U.S.A.) (heavy label) as described previously ^75^. The dimethyl-labeled spike peptides were eluted with an 80% acetonitrile and 0.1% TFA solution, and dried with a SpeedVac (Thermo Fisher Scientific, U.S.A.).

For the time course sampling for autophosphorylation detection, one timepoint sample contains 0.3 μM purified GST-CaMKIIβ protein (Carna Biosciences, Japan), 50 mM HEPES-NaOH pH 7.6, 150 mM NaCl, 1 mM MgCl_2_, 0.25% (v/v) NP-40 and 2.5 mM ATP. The sample without CaM was sampled and used as “0 min” time point. Then, 0.5 mM CaCl_2_ and 10 mM EGTA were added to the indicated conditions shown in Figure 7a. The kinase reaction was initiated by adding 0.5 μM CaM to each sample. During the time course sampling, 10 mM EGTA was added for the condition named “0.5 mM Ca^2+^, 10 mM EGTA at 5 min (Condition #4)”. Note that for the quantification of S182 phosphorylation, 10-fold higher concentration of purified GST-CaMKIIβ and CaM were used because of the low signal sensitivity of the corresponding phosphorylated peptide.

The kinase reaction was terminated by adding an equal volume of Solution B and incubating at 98 °C for 30 min. The samples were reduced, alkylated, and digested by proteases according to the PTS method ^74^ as described above except that 1 μg of Lys-C and 1 μg of trypsin were used for most of the samples, and 1 μg of Lys-C and 1 μg of Glu-C (Promega, U.S.A.) were used for the sample for quantifying S182 phosphorylation.

The dried peptides were solubilized in 1 mL of a 2% acetonitrile and 0.1% TFA solution, and trapped on C18 StageTips ^77^. The trapped peptides were subjected to dimethyl-labeling with formaldehyde (light label) as described above. An additional GST-CaMKIIβ sample independent from the time course sampling were prepared as an internal control reference, and subjected to dimethyl-labeling with CD_2_O (medium label). The dimethyl-labeled peptides on the tip were eluted with an 80% acetonitrile and 0.1% TFA solution. Then, 1/30 volume of the light-labeled samples were isolated and mixed with equal amounts of medium label peptides. This allowed us to compare the relative amount of GST-CaMKIIβ in the individual time course samples with each other using the medium-labeled internal control.

The remainder of the light-labeled samples were mixed with the mixture of heavy labeled spike peptides and applied to High-Select™ Fe-NTA Phosphopeptide Enrichment Kit (Thermo fisher Scientific, U.S.A.) to enrich the phosphorylated peptides. This allowed us to compare the relative amount of phosphorylated peptides in the individual time course samples with each other using the heavy labeled spike peptides.

All analytical samples were dried with a SpeedVac (Thermo Fisher Scientific, U.S.A.) and dissolved in a 2% acetonitrile and 0.1% TFA solution. Mass-spectrometry-based quantification was carried out by SRM analysis using a TSQ Quantiva triple-stage quadrupole mass spectrometer (Thermo Fisher Scientific, U.S.A.) as described above. The amount of each phosphorylated peptide was normalized to the amount of total GST-CaMKIIβ quantified using the average amounts of several non-phosphorylated peptides.

### Production of *Camk2b* KO mice

*Camk2b* KO mice were generated using the Triple-target CRISPR method described previously ^24^. C57BL/6N females (4–6 weeks old, CLEA Japan, Japan) were superovulated and mated with C57BL/6N males (CLEA Japan, Japan). The fertilized eggs were collected from the ampulla of the oviduct of plugged C57BL/6N females by micro-dissection and kept in KSOM medium (Merck, Germany or ARK Resource, Japan) in a 5% CO_2_ incubator at 37°C. The design of gRNAs for *Camk2b* was previously shown as set 1 in a previous study ^10^. In the previous study, an independent set of gRNA called set 2 was also tested. A significant decrease in the sleep duration was observed both in set 1 and set 2 gRNA-injected mice, suggesting that at least a major part of sleep phenotype is not due to the off-target effect of injected gRNAs ^10^. The synthesized gRNAs for *Camk2b* (150 ng/µl in total) and *Cas9* mRNA (100 ng/µl) were co-injected into the cytoplasm of fertilized eggs in M2 medium (Merck, Germany or ARK Resource, Japan) at room temperature. After microinjection, the embryos were cultured for 1 h in KSOM medium (Merck, Germany, or ARK Resource, Japan) in a 5% CO_2_ incubator at 37°C. 15–30 embryos were then transferred to the oviducts of pseudopregnant female ICR mice.

Genotyping of KO mice was conducted with the same protocol described previously ^24^. qPCR was performed using genomic DNA purified from tails of WT and KO mice and primers which were annealed to the target sequences. The target site abundance was calculated using a standard curve obtained from wild-type genomic DNA. The amount of *Tbp* ^78^ was quantified with a pair of primers (5’-CCCCCTCTGCACTGAAATCA-3’; 5’-GTAGCAGCACAGAGCAAGCAA-3’) and used as an internal control. When the amplified intact DNA by qPCR is less than 0.5% of wild-type genome, we judged that the target DNA is not detectable. When any of three targets was not detected, we classified the animal as a KO. When we could not confirm KO genotype by the qPCR, we performed 2nd qPCR using the alternative primer which was independent of 1st qPCR. In the case of Camk2b set 1 KO, first and second targets of triple CRISPR gRNA were judged as not detectable by 2 nd qPCR. The result of qPCR is shown in **Figure 4-figure supplement 1d** and the primer list used for the qPCR is shown below.

1st qPCR primer pairs:

*Camk2b* set 1, target #1

Forward: 5’-CCACAGGGGTGATCCTGTATATCCTGC-3’
Reverse: 5’-CTGCTGGTACAGCTTGTGTTGGTCCTC-3’
*Camk2b* set 1, target #2

Forward: 5’-GGAAAATCTGTGACCCAGGCCTGAC-3’
Reverse: 5’-TCTGTGGAAATCCATCCCTTCGACC-3’
*Camk2b* set 1, target #3

Forward: 5’-GAACCCGCACGTGCACGTCATTGGC-3’
Reverse: 5’-CCCTGGCCATCGATGTACTGTGTG-3’

2nd qPCR primer pairs:

*Camk2b* set 1, target #1

Forward: 5’-CAGAAAGGTGGGTAGCCCACCAGCAGG-3’
Reverse: 5’-CTATGCTGCTCACCTCCCCATCCACAG-3’
*Camk2b* set 1, target #2

Forward: 5’-GCCTGAAGCTCTGGGCAACCTGGTCG-3’
Reverse: 5’-CCACCCCAGCCTTTTCACTCACGGTTCTC-3’
*Camk2b* set 1, target #3

Forward: 5’-GCATCGCCTACATCCGCCTCACAC-3’
Reverse: 5’-CGGTGCCACACACGGGTCTCTTCGGAC-3’

### Production of *Camk2b^FLAG/FLAG^* mice

FLAG-tag sequence was inserted into the endogenous *Camk2b* locus (prior to the stop codon) by single-stranded oligodeoxynucleotide (ssODN) and CRISPR/Cas9-mediated knock-in. The gRNA target sequence and a donor sequence were selected according to previous study ^79^. Preparation of gRNA and Cas9, and general procedures for obtaining the genetically modified mouse were conducted according to previous study ^10^. Following primer sequences were used to produce gRNA targeting the *Camk2b* locus.

*Camk2b-*FLAG gRNA primer forward #1

5’-CACTATAGGCAGTGGCCCCGCTGCAGTGGTTTTAGAGCTAGAAATAGC -3’

*Camk2b-*FLAG gRNA primer forward #2

5’-GGGCCTAATACGACTCACTATAGGCAGTGGCCCCGCTGCAGTGG -3’

*Camk2b-*FLAG gRNA primer reverse #1

5’-AAAAGCACCGACTCGGTGCC -3’

A donor ssODN (*sequence*) was synthesized by Integrated DNA Technologies. *Camk2b-*FLAG ssODN (capital letter: FLAG tag sequence)

5’-aagagacccgtgtgtggcaccgccgcgacggcaagtggcagaatgtacatttccactgctcgggcgctccagtggccccgctgcagGACTACAAGGACGACGATGACAAGtgaggtgagtccctgcggtgtgcgtagggcagtgcggcatgcgtgggacagtgcagcgtgcatggggtgtggcccagtgcagcgtgc -3’

1∼2 pL of RNase free water (Nacalai Tesque Inc.) containing 100 ng/µl gRNA, 100 ng/µl Cas9 mRNA and 100 ng/µl ssODN was injected into the cytoplasm of fertilized eggs in M2 medium (Merck, Germany or ARK Resource, Japan) at room temperature. After microinjection, the embryos were cultured for 1 h in KSOM medium (Merck, Germany, or ARK Resource, Japan) in a 5% CO_2_ incubator at 37°C. 15–30 embryos were then transferred to the oviducts of pseudopregnant female ICR mice.

Genomic DNA of F_0_ mice tails was extracted with NucleoSpin Tissue kit (Takara Bio, Japan) according to the manufacturer’s protocol. The genotyping PCR was conducted by using following primer pairs to select heterozygous or homozygous FLAG knock-in offspring. Genotyping was based on the size and direct sequencing of the PCR amplicon. The obtained heterozygous or homozygous FLAG knock-in F_0_ mice were crossed with wildtype C57BL/6N mice to obtain heterozygous FLAG knock-in F_1_ mice.

*Camk2b*-FLAG genotyping primer pairs:

Pair #1

Forward: 5’-ACGACCAACTCCATTGCTGAC -3’
Reverse: 5’-CTACATCCGCCTCACACAGTACATC -3’
Pair #2

Forward: 5’-ACGACCAACTCCATTGCTGAC -3’
Reverse: 5’-GACTACAAGGACGACGATGACAAG -3’
Pair #3

Forward: 5’-CTTGTCATCGTCGTCCTTGTAGTC -3’
Reverse: 5’-CTACATCCGCCTCACACAGTACATC -3’

### Sleep measurement with the SSS

The SSS system enables fully automated and noninvasive sleep/wake phenotyping^24^. The SSS recording and analysis were carried out according to the protocol described previously ^24^. The light condition of the SSS rack was set to light/dark (12 h periods) or constant dark. Mice had *ad libidum* access to food and water. In the normal measurement, eight-week-old mice were placed in the SSS chambers for one to two weeks for sleep recordings. For data analysis, we excluded the first day and used six days of measurement data. For the *Cry1/2* DKO and *Per1/2* DKO mutant mice, recordings were performed under light/dark conditions for two weeks followed by constant dark conditions for two weeks. For data analysis, we excluded the first day and used four days of measurement data under each light condition. Sleep staging was performed in every 8-second epoch.

Sleep parameters, such as sleep duration, *P_WS_*, and *P_SW_* were defined previously ^24^. In the SSS, sleep staging was performed every 8 seconds, which is the smallest unit called “epoch”. When we focus on two consecutive epochs, there are four combinations: keeping awake state (wake to wake), keeping sleep state (sleep to sleep), transition from wakefulness to sleep (wake to sleep), and transition from sleep to wakefulness (sleep to wake). Transition probabilities were calculated from all two consecutive epochs in the measurement period. The definition of transition probabilities are as follows: *P_WS_* (transition probability from wake to sleep) is defined as *P_WS_* = *N_WS_* / (*N_WS_* + *N_WW_*), and *P_SW_* (transition probability from sleep to wake) is defined as *P_SW_* = *N_SW_* / (*N_SW_* + *N_SS_*), where *N_mn_* is the number of transitions from state m to n (m, n ∈ {sleep, awake}) in the observed period. The balance between *P_WS_* and *P_SW_* determines the total sleep time, i.e., mice with longer sleep time tend to have increased *P_WS_* and/or decreased *P_SW_*. *P_WS_* and *P_SW_* are independent of each other, and it can be deduced from the definition that *P_WS_* + *P_WW_* = 1 and *P_SW_* + *P_SS_* = 1. The sleep episode duration is the average of the time spent in each consecutive sleep phase during the observed period.

### Sleep measurement with EEG/EMG recording

For EEG/EMG recording, AAV-administrated six-week-old C57BL/6N mice were used for surgery. For the recording of *Camk2b* KO mice, 16-17-week-old *Camk2b* KO mice and WT control mice at the same age were used for surgery. Wired and wireless recording method are used in parallel for EEG/EMG measurements, and we have confirmed that these two methods give qualitatively comparable results.

For wireless recordings, anesthetized mice were implanted a telemetry transmitter (DSI, U.S.A). As EEG electrodes, two stainless steel screws were connected with lines from the transmitter and embedded in the skull of the cortex (anteroposterior, +1.0 mm; right, +1.5 mm from bregma or lambda). As EMG electrodes, two lines from the transmitter were placed in the trapezius muscles. After the surgery, the mice were allowed to recover for at least ten days. EEGs and EMGs were recoded wirelessly. The mice had access to food and water. The sampling rate was 100 Hz for both EEG and EMG. The detailed methods were described previously ^80^.

For wired recordings, mice were implanted with EEG and EMG electrodes for polysomnographic recordings. To monitor EEG signals, two stainless steel EEG recording screws with 1.0 mm in diameter and 2.0 mm in length were implanted on the skull of the cortex (anterior, +1.0 mm; right, +1.5 mm from bregma or lambda). EMG activity was monitored through stainless steel, Teflon-coated wires with 0.33 mm in diameter (AS633, Cooner Wire, California, U.S.A) placed into the trapezius muscle. The EEG and EMG wires were soldered to miniature connector with four pins in 2 mm pitch (Hirose Electric, Japan). Finally, the electrode assembly was fixed to the skull with dental cement (Unifast III, GC Corporation, Japan). After 10 days of recovery, the mice were placed in experimental cages with a connection of spring supported recording leads. The EEG/EMG signals were amplified (Biotex, Japan), filtered (EEG, 0.5–60 Hz; EMG, 5–128 Hz), digitized at a sampling rate of 128 Hz, and recorded using VitalRecorder software (KISSEI Comtec, Japan).

For the sleep staging, we used the FASTER method ^80^ with some modifications to automatically annotate EEG and EMG data. 24 h of recording data were used for the analysis. Sleep staging was performed every 8-second epoch. Finally, the annotations were manually checked.

The power spectrum density was calculated for each epoch by fast Fourier transformation (FFT) with Welch’s averaging method. Briefly, each 8 s segment was further divided into eight overlapping sequences. The overlapping length was 50% of each sequence. The Hamming window was applied onto the sequences before the FFT and the obtained spectrum was averaged over the eight sequences. The dirty segments were excluded from the subsequent processes ^80^. The power spectrum of each behavioral state (Wake, NREM, REM) was calculated by averaging the power spectra (1-50 Hz) of segments within each state over the observation period. The calculated power spectra were normalized by the total power. The power density in typical frequency domains were calculated as the summation of the powers in each frequency domain (slow, 0.5-1 Hz; delta, 0.5-4 Hz; theta, 6-10 Hz).

Transition probabilities between wakefulness, NREM sleep, and REM sleep were calculated same as previously reported ^81^. For example, *P_NW_* = *N_NW_* / (*N_NW_* +*N_NR_ + N_NN_*), where *N_mn_* is the number of transitions from state m to n (m,n ∈ {wake, NREM sleep, REM sleep}) in the observed period.

### Cage change experiment

For cage change experiment (**Figure 2-figure supplement 1**), AAV-administrated mice (9-week-old) were placed in the SSS chambers and habituated to the environment for three days. On the fourth day, the SSS chamber was replaced with a new one at ZT0. The sleep data of the fourth day was analyzed. The data of the first three days were used for baseline calculation.

### ES-mice production

Genetically modified mice were produced using the previously reported ES-mouse method, which allows us to analyze the behavior of F0 generation mice without crossing ^82, 83^. Mouse ES cells (ESCs) were established from blastocysts in 3i medium culture conditions as described previously ^84^. Mouse strains used for the ESC establishment were as follows: *Cry1*^-/-^:*Cry2*^-/-^, *Cry1*^-/-^:*Cry2*^-/-^mouse ^38^; *Per1*^- /-^:*Per2*^-/-^, *Per1*^-/-^:*Per2*^-/-^ mouse ^40^; *Vglut2*-*Cre*, heterozygous *Slc17a6^tm2(cre)Lowl^*/J mouse (The Jakson Laboratory, JAX stock #016963) ^85^; *Gad2*-*Cre*, heterozygous *Gad2^tm2(cre)Zjh^*/J mouse (The Jakson Laboratory, JAX stock #010802) ^86^.

Male ESCs were cultured as described previously ^82, 83^. Before cultivation, PURECoat^TM^ amine dishes (Beckton-Dickinson, NJ, U.S.A.) was treated with a medium containing LIF plus 6-bromoindirubin-30-oxime (BIO) ^87^ for more than 5 h at 37°C with 5% CO_2_. ESCs were seeded at 1 × 10^5^ cells per well and maintained at 37°C in 5% CO_2_ under humidified conditions with a 3i culture medium (Y40010, Takara Bio, Japan) without feeder cells. The expanded ESCs were collected by adding 0.25% trypsin-EDTA solution and prepared as a cell suspension. 10–30 ESCs were injected into each ICR (CLEA Japan, Japan) 8-cell-stage embryo and the embryos were transferred into the uterus of pseudopregnant ICR female mice (SLC, Japan). We determined the contribution of the ESCs in an obtained ES-mouse by its coat color following a previously reported protocol ^82, 83^. The ES mice uncontaminated with ICR-derived cells were used for the experiment.

### AAV production

The protocol for AAV production was based on the previously reported protocol ^88^ with some modifications. AAV pro 293T (Takara Bio, Japan) was cultured in 150 mm dishes (Corning, USA) in a culture medium containing DMEM (high glucose) (Thermo Fisher Scientific, U.S.A.), 10% (v/v) FBS, and penicillin-streptomycin (Thermo Fisher Scientific, U.S.A.) at 37°C in 5% CO_2_ under humidified conditions. pAAV, pUCmini-iCAP-PHPeB and pHelper plasmid (Agilent, U.S.A.) were transfected into cells at 80%–90% confluency using polyethyleneimine (Polysciences, U.S.A.). We employed a pAAV: pUCmini-iCAP-PHPeB: pHelper plasmid ratio of 1:4:2 based on micrograms of DNA (e.g. 5.7 μg of pAAV, 22.8 μg of pUCmini-iCAP-PHP, and 11.4 μg of pHelper). On the day following the transfection, the culture medium was replaced with 20 ml of a culture medium containing DMEM (high glucose, Glutamax) (Thermo Fisher Scientific, U.S.A.), 2% (v/v) FBS, MEM Non-Essential Amino Acids solution (NEAA) (Thermo Fisher Scientific, U.S.A.), and penicillin-streptomycin. On the third day following the transfection, the culture medium was collected and replaced with 20 ml of new culture medium containing DMEM (high glucose, Glutamax), 2% (v/v) FBS, MEM NEAA, and penicillin-streptomycin. The collected culture medium was stored at 4°C. On the fifth day following the transfection, the cells and the culture medium were collected and combined with the stored medium. The suspension was separated into supernatant and cell pellet by centrifugation (2000 × g, 20min). From the supernatant, AAVs were concentrated by adding polyethylene glycol at a final concentration of 8% followed by centrifugation. From the cells, AAVs were extracted in a Tris-MgCl_2_ buffer (10 mM Tris pH 8.0, 2 mM MgCl_2_) by repetitive freeze-thaw cycles. The obtained extract containing AAV was treated with Benzonase (100 U/ml) in a Tris-MgCl_2_ buffer, and then AAVs were purified by ultracentrifugation at 350,000 × g for 2 h 25 min (himac CP80WX and P70AT rotor, HITACHI, Japan) with Iodixanol density gradient solutions (15%, 25%, 40%, and 60% (wt/vol)). Viral particles were contained in a 40% solution, and this solution was ultrafiltered with an Amicon Ultra-15 device (100 kDa, Merck, Germany) to obtain the AAV stock solution for administration to mice.

To determine the AAV titer, virus solution was treated with Benzonase (50 U/ml, 37°C, 1 h) followed by Proteinase K (0.25 mg/ul, 37°C, 1 h). Subsequently, the viral genome was obtained by phenol-chloroform-isoamyl alcohol extraction followed by isopropanol precipitation. The AAV titer (vg/ml) was calculated by quantifying the number of WPRE sequences in the sample by qPCR using plasmid as a standard. The qPCR protocol was 60 s at 95°C for preheating (initial denaturation) and 45 cycles from 10 s at 95°C to 30 s at 60°C using TB Green *Premix Ex Taq*^TM^ GC (Takara Bio, Japan).

### Retro orbital injection of AAV to mice

Six-week-old male mice were anesthetized with 2%–4% isoflurane and injected with 100 μl of AAV in their retro orbital sinus. **Table 1** summarizes the AAVs used in this study and their administration conditions. The AAV-administrated mice were subjected to sleep phenotyping at eight-week-old.

**Table 1.** Summary of AAV applications and conditions

| Applications in the paper | Use cases | pAAV vector | Dosage (vg/mouse) |
| --- | --- | --- | --- |
| Neuronal expression of *Camk2b* | Fig.1d-f, Fig.5, Fig.6, Fig.7b-c, Fig.1-Sup1b, Fig1-Sup2a-e, Fig.2-Sup1e-j, Fig.5-Sup1a-e, Fig.6-Sup1, Fig.7-Sup1a-c | P(hSyn1)-Camk2b-3'UTR-WPRE-SV40pA | 5.0 x 10<sup>10</sup> |
| Neuronal expression of H2B-mCherry | Fig.1c, Fig.1-Sup1a | P(hSyn1)-H2B-mCherry-3'UTR-WPRE-SV40pA | 5.0 x 10<sup>10</sup> |
| Camk2a promoter-driven expression of *Camk2b* | Fig.2, Fig.2-Sup1d | P(Camk2a)-Camk2b-3'UTR-WPRE-SV40pA | 1.0 x 10<sup>11</sup> |
| Camk2b promoter-driven expression of *Camk2b* | Fig.2-Sup1a-d | P(Camk2b)-Camk2b-3'UTR-WPRE-SV40pA | 2.0 x 10<sup>11</sup> |
| Cre-dependent expression of *Camk2b* | Fig.3b-e | P(hSyn1)-DIO-Camk2b-3'UTR-WPRE-SV40pA | 2.0 x 10<sup>11</sup> |
| Neuronal expression of *Camk2b* deletion mutant | Fig.4b-c | P(hSyn1)-Camk2b(del)-3'UTR-WPRE-SV40pA | 2.5 x 10<sup>10</sup> |
| Neuronal expression of AIP2 | Fig.4e-j, Fig.4-Sup1a-c | P(hSyn1)-mCherry-AIP2-Map2DTE-WPRE-SV40pA | 2.0 x 10<sup>11</sup> |

### Estimation of transduction efficiency

Transduction efficiency was estimated based on previous reports ^89, 90^. After the sleep phenotyping, the brain hemisphere except for the olfactory bulb and cerebellum was collected from the AAV administrated mouse. Brain DNA was purified using an Agencourt DNAdvance (BECKMAN COULTER, U.S.A.). The copy numbers of both the AAV vector genomes and mouse genomic DNA were quantified with a standard curve generated from known amounts of DNA. Vector genomes per cell were calculated by dividing the copy number of AAV vector genomes by diploid copies of the *Tbp* gene in the sample. The copy number of the AAV vector genomes and the *Tbp* gene were determined with WPRE-binding primers (5’- CTGTTGGGCACTGACAATTC-3’, 5’-GAAGGGACGTAGCAGAAGGA-3’) and Tbp-binding primers (5’-CCCCCTCTGCACTGAAATCA-3’; 5’- GTAGCAGCACAGAGCAAGCAA-3’) ^78^, respectively. The qPCR protocol was 60 s at 95°C for preheating (initial denaturation) and 45 cycles from 10 s at 95°C to 30 s at 60°C using a TB Green *Premix Ex Taq*^TM^ GC (Takara Bio, Japan).

### Clustering analysis

The character of each mutant was extracted by principal component analysis using the values of *P_WS_* and *P_SW_*. The first and second principal components were used for hierarchical clustering using Ward’s algorithm. The threshold was set to 40% of the distance between the farthest clusters (**Figure 6-figure supplement 1a**). The principal component analysis and clustering were performed using Python 3.8.0 with the numpy 1.18.5, scikit-learn 0.23.1 and scipy 1.5.0 libraries.

### Statistics

No statistical method was used to predetermine the sample size. The sample sizes were determined based on previous experiences and reports. Experiments were repeated at least two times with the independent sets of the animals or independently prepared cell lysates. The series of single/double phosphomimetic screening was not repeated, but the mutants we focused on from the screening results were further analyzed in detail through additional independent experiments. In the sleep analysis, individuals with abnormal measurement signals or weakened individuals were excluded from the sleep data analyses because of their difficulties in accurate sleep phenotyping.

Statistical analyses were performed by Microsoft Excel and R version 3.5.2. Statistical tests were performed by two-sided. To compare two unpaired samples, the normality was tested using the Shapiro test at a significance level of 0.05. When the normality was not rejected in both groups, the homogeneity of variance was tested using the *F*-test at a significance level of 0.05. When the null hypothesis of a normal distribution with equal variance for the two groups was not rejected, a Student’s *t*-test was used. When the normality was not rejected but the null hypothesis of equal variance was rejected, a Welch’s *t*-test was used. Otherwise, a two-sample Wilcoxon test was applied.

To compare more than two samples against an identical sample, the normality was tested with the Kolmogorov-Smirnov test at a significance level of 0.05. When the normality was not rejected in all groups, the homogeneity of variance was tested with Bartlett’s test at a significance level of 0.05. When the null hypothesis of a normal distribution with equal variance was not rejected for all groups, Dunnett’s test was used. Otherwise, Steel’s test was applied.

For multiple comparisons between each group, the Tukey-Kramer test was used when the null hypothesis of a normal distribution with equal variance was not rejected for all groups. Otherwise, Steel-Dwass test was applied.

In this study, p < 0.05 was considered significant (*p < 0.05, **p < 0.01, ***p < 0.001, and n.s. for not significant). **Figure 8** summarizes the workflow for selecting statistical method and the statistical analyses used in each experiment of this study and *P* values.

**Figure 8.**
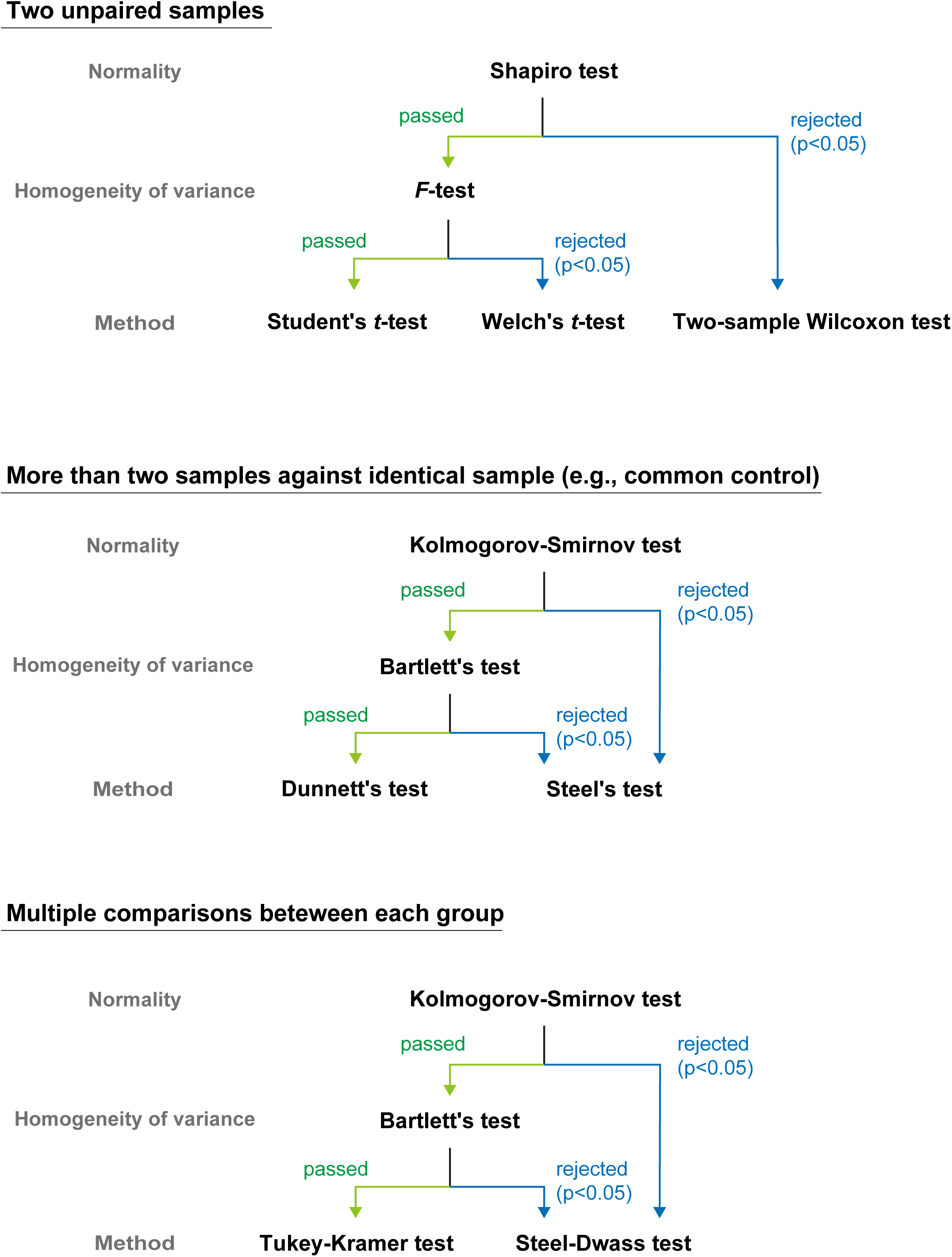
Workflow for selecting the statistical method. Workflow for selecting the statistical test methods used in this study. Based on the purpose of the comparison, normality and equality of variance were checked, and appropriate statistical method was selected. Details are provided in the Methods section.

## ACKNOWLEDGMENTS

We thank all the lab members at RIKEN Center for Biosystems Dynamics Research (BDR) and the University of Tokyo, in particular, Masazumi Tanaka, Jun-ichi Kuroda, and Ayaka Saito for technical assistance of the biochemical analysis; Etsuo A. Susaki, Rurika Itofusa, Takeyuki Miyawaki, Chika Shimizu and Kimiko Itayama for AAV preparation; Masako Kunimi and Ruriko Inoue, Sachiko Tomita, for help with sleep phenotyping; Yumika Sugihara, Natsumi Hori, Eriko Matsushita, and Yuichi Uranyu for animal experiment. We also thank members at LARGE, RIKEN BDR for help with ES-mouse production; Chiaki Masuda for kind instructions on AAV production; Hirokazu Hirai for providing pAAV plasmid and helpful suggestions on the experiment.

This work was supported by grants from the Brain/MINDS JP20dm0207049, Science and Technology Platform Program for Advanced Biological Medicine JP21am0401011, AMED-CREST 18gm0610006h0006 (AMED/MEXT) (H.R.U.), Grant-in-Aid for Scientific Research (S) JP25221004 (JSPS KAKENHI) (H.R.U.) and Scientific Research (C) JP20K06576 (JSPS KAKENHI) (K.L.O.), Grant-in-Aid for Early-Career Scientists JP19K16115 (JSPS KAKENHI) (D.T.), HFSP Research Grant Program RGP0019/2018 (HFSP) (H.R.U.), ERATO JPMJER2001 (JST) (H.R.U.) and an intramural Grant-in-Aid from the RIKEN BDR (H.R.U.). The authors would like to thank Enago (www.enago.jp) for the English language review.

## COMPETING INTERESTS

H.R.U conducted a collaborative research project with Thermo Fisher Scientific Inc. Y.N. is an employee of Thermo Fisher Scientific, Inc. The company provided support in the form of salary for Y.N., and technical advice on the setup of mass spectrometers. However, the company did not have any additional role in the study design, data collection and analysis, decision to publish, or preparation of the manuscript.

## LIST OF SOURCE FILES

**Figure1-sourcedata 1**

Sleep phenotypes of mice expressing CaMKIIβ single D mutants.

**Figure1-sourcedata 2**

Quantified values of CaMKIIα/β-derived peptides.

**Figure1-sourcedata 3**

SRM transition list for the MS-based quantification of CaMKIIα/β-derived peptides.

**Figure1-figure-supplement 1-sourcedata 1**

Uncropped image of western blotting data.

**Figure1-figure-supplement 1-sourcedata 2**

Raw image files of western blotting data.

**Figure1-figure-supplement 2-sourcedata 1**

Sleep phenotypes of mice expressing CaMKIIβ single D mutants.

**Figure1-figure-supplement 3-sourcedata 1**

*In vitro* expression levels and kinase activities of CaMKIIβ single D mutants.

**Figure2-sourcedata 1**

Sleep phenotypes of mice expressing CaMKIIβ T287D-related mutants.

**Figure2-figure-supplement 1-sourcedata 1**

Sleep phenotypes of mice expressing CaMKIIβ T287D-related mutants.

**Figure3-sourcedata 1**

Sleep phenotypes of Cre-mice expressing CaMKIIβ T287D mutant.

**Figure4-sourcedata 1**

Sleep phenotypes of mice expressing kinase domain of CaMKIIβ or CaMKII inhibitor peptide.

**Figure4-figure-supplement 1-sourcedata 1**

EEG/EMG data of *Camk2b* knockout mice or mice expressing CaMKII inhibitor peptide.

**Figure5-sourcedata 1**

Sleep phenotypes of mice expressing CaMKIIβ double D mutants.

**Figure5-figure-supplement 1-sourcedata 1**

Sleep phenotypes and *in vitro* kinase activities of CaMKIIβ double D mutants.

**Figure6-sourcedata 1**

Sleep phenotypes of mice expressing CaMKIIβ T306D:T307D-related mutants.

**Figure6-figure-supplement 1-sourcedata 1**

Sleep phenotypes of mice expressing CaMKIIβ T306D:T307D-related mutants.

**Figure6-figure-supplement 2-sourcedata 1**

*In vitro* kinase activities of CaMKIIβT306D:T307D-related mutants.

**Figure7-sourcedata 1**

Quantified values of peptides derived from purified CaMKIIβ.

**Figure7-sourcedata 2**

SRM transition list for the MS-based quantification of CaMKIIβ-derived peptides.

**Figure7-sourcedata 3**

Sleep phenotypes of mice expressing CaMKIIβ with the combined mutation at sleep maintenance and cancelation of sleep induction residues.

**Figure7-figure-supplement 1-sourcedata 1**

Sleep phenotypes of mice expressing CaMKIIβ with the combined mutation at sleep maintenance and cancelation of sleep induction residues.

## REFERENCES

1 Partch, C. L. Orchestration of Circadian Timing by Macromolecular Protein Assemblies. J Mol Biol, (2020).

2 Ode, K. L. & Ueda, H. R. Design Principles of Phosphorylation-Dependent Timekeeping in Eukaryotic Circadian Clocks. Cold Spring Harb Perspect Biol 10, (2018).

3 Millius, A., Ode, K. L. & Ueda, H. R. A period without PER: understanding 24-hour rhythms without classic transcription and translation feedback loops. F1000Res 8, (2019).

4 Konopka, R. J. & Benzer, S. Clock mutants of Drosophila melanogaster. Proc Natl Acad Sci U S A 68, 2112–2116, (1971).

5 Toh, K. L. et al. An hPer2 phosphorylation site mutation in familial advanced sleep phase syndrome. Science 291, 1040–1043, (2001).

6 Xu, Y. et al. Modeling of a human circadian mutation yields insights into clock regulation by PER2. Cell 128, 59–70, (2007).

7 Masuda, S. et al. Mutation of a PER2 phosphodegron perturbs the circadian phosphoswitch. Proc Natl Acad Sci U S A 117, 10888–10896, (2020).

8 Isojima, Y. et al. CKIepsilon/delta-dependent phosphorylation is a temperature-insensitive, period-determining process in the mammalian circadian clock. Proc Natl Acad Sci U S A 106, 15744–15749, (2009).

9 Frank, M. G. & Heller, H. C. The Function(s) of Sleep. Handb Exp Pharmacol 253, 3–34, (2019).

10 Tatsuki, F. et al. Involvement of Ca(2+)-Dependent Hyperpolarization in Sleep Duration in Mammals. Neuron 90, 70–85, (2016).

11 Diering, G. H. et al. Homer1a drives homeostatic scaling-down of excitatory synapses during sleep. Science 355, 511–515, (2017).

12 Wang, Z. et al. Quantitative phosphoproteomic analysis of the molecular substrates of sleep need. Nature 558, 435–439, (2018).

13 Bruning, F. et al. Sleep-wake cycles drive daily dynamics of synaptic phosphorylation. Science 366, (2019).

14 Ode, K. L. & Ueda, H. R. Phosphorylation Hypothesis of Sleep. Front Psychol 11, 575328, (2020).

15 Funato, H. et al. Forward-genetics analysis of sleep in randomly mutagenized mice. Nature 539, 378–383, (2016).

16 Park, M. et al. Loss of the conserved PKA sites of SIK1 and SIK2 increases sleep need. Sci Rep 10, 8676, (2020).

17 Mikhail, C., Vaucher, A., Jimenez, S. & Tafti, M. ERK signaling pathway regulates sleep duration through activity-induced gene expression during wakefulness. Sci Signal 10, (2017).

18 Bayer, K. U. & Schulman, H. CaM Kinase: Still Inspiring at 40. Neuron 103, 380–394, (2019).

19 Coultrap, S. J. & Bayer, K. U. CaMKII regulation in information processing and storage. Trends Neurosci 35, 607–618, (2012).

20 Miller, S. G., Patton, B. L. & Kennedy, M. B. Sequences of autophosphorylation sites in neuronal type II CaM kinase that control Ca2(+)- independent activity. Neuron 1, 593–604, (1988).

21 Meyer, T., Hanson, P. I., Stryer, L. & Schulman, H. Calmodulin trapping by calcium-calmodulin-dependent protein kinase. Science 256, 1199–1202, (1992).

22 Mullasseril, P., Dosemeci, A., Lisman, J. E. & Griffith, L. C. A structural mechanism for maintaining the ’on-state’ of the CaMKII memory switch in the post-synaptic density. J Neurochem 103, 357–364, (2007).

23 Thornquist, S. C., Langer, K., Zhang, S. X., Rogulja, D. & Crickmore, M. A. CaMKII Measures the Passage of Time to Coordinate Behavior and Motivational State. Neuron 105, 334–345 e339, (2020).

24 Sunagawa, G. A. et al. Mammalian Reverse Genetics without Crossing Reveals Nr3a as a Short-Sleeper Gene. Cell Rep 14, 662–677, (2016).

25 Chemelli, R. M. et al. Narcolepsy in orexin knockout mice: molecular genetics of sleep regulation. Cell 98, 437–451, (1999).

26 Griffith, L. C. Regulation of calcium/calmodulin-dependent protein kinase II activation by intramolecular and intermolecular interactions. J Neurosci 24, 8394–8398, (2004).

27 Patton, B. L., Miller, S. G. & Kennedy, M. B. Activation of type II calcium/calmodulin-dependent protein kinase by Ca2+/calmodulin is inhibited by autophosphorylation of threonine within the calmodulin-binding domain. J Biol Chem 265, 11204–11212, (1990).

28 Colbran, R. J. & Soderling, T. R. Calcium/calmodulin-independent autophosphorylation sites of calcium/calmodulin-dependent protein kinase II. Studies on the effect of phosphorylation of threonine 305/306 and serine 314 on calmodulin binding using synthetic peptides. J Biol Chem 265, 11213–11219, (1990).

29 Lou, L. L. & Schulman, H. Distinct autophosphorylation sites sequentially produce autonomy and inhibition of the multifunctional Ca2+/calmodulin- dependent protein kinase. J Neurosci 9, 2020–2032, (1989).

30 Baucum, A. J., 2nd, Shonesy, B. C., Rose, K. L. & Colbran, R. J. Quantitative proteomics analysis of CaMKII phosphorylation and the CaMKII interactome in the mouse forebrain. ACS Chem Neurosci 6, 615–631, (2015).

31 Myers, J. B. et al. The CaMKII holoenzyme structure in activation-competent conformations. Nat Commun 8, 15742, (2017).

32 Vyazovskiy, V. V., Cirelli, C., Pfister-Genskow, M., Faraguna, U. & Tononi, G. Molecular and electrophysiological evidence for net synaptic potentiation in wake and depression in sleep. Nat Neurosci 11, 200–208, (2008).

33 Blanco, W. et al. Synaptic Homeostasis and Restructuring across the Sleep-Wake Cycle. PLoS Comput Biol 11, e1004241, (2015).

34 Chan, K. Y. et al. Engineered AAVs for efficient noninvasive gene delivery to the central and peripheral nervous systems. Nat Neurosci 20, 1172–1179, (2017).

35 Torres-Ocampo, A. P. et al. Characterization of CaMKIIalpha holoenzyme stability. Protein Sci 29, 1524–1534, (2020).

36 Nathanson, J. L., Yanagawa, Y., Obata, K. & Callaway, E. M. Preferential labeling of inhibitory and excitatory cortical neurons by endogenous tropism of adeno-associated virus and lentivirus vectors. Neuroscience 161, 441–450, (2009).

37 Kon, N. et al. CaMKII is essential for the cellular clock and coupling between morning and evening behavioral rhythms. Genes Dev 28, 1101–1110, (2014).

38 van der Horst, G. T. et al. Mammalian Cry1 and Cry2 are essential for maintenance of circadian rhythms. Nature 398, 627–630, (1999).

39 Vitaterna, M. H. et al. Differential regulation of mammalian period genes and circadian rhythmicity by cryptochromes 1 and 2. Proc Natl Acad Sci U S A 96, 12114–12119, (1999).

40 Bae, K. et al. Differential functions of mPer1, mPer2, and mPer3 in the SCN circadian clock. Neuron 30, 525–536, (2001).

41 Zheng, B. et al. Nonredundant roles of the mPer1 and mPer2 genes in the mammalian circadian clock. Cell 105, 683–694, (2001).

42 Suzuki, A., Sinton, C. M., Greene, R. W. & Yanagisawa, M. Behavioral and biochemical dissociation of arousal and homeostatic sleep need influenced by prior wakeful experience in mice. Proc Natl Acad Sci U S A 110, 10288–10293, (2013).

43 Zou, D. J. & Cline, H. T. Expression of constitutively active CaMKII in target tissue modifies presynaptic axon arbor growth. Neuron 16, 529–539, (1996).

44 Ishida, A. et al. Critical amino acid residues of AIP, a highly specific inhibitory peptide of calmodulin-dependent protein kinase II. FEBS Lett 427, 115–118, (1998).

45 Murakoshi, H. et al. Kinetics of Endogenous CaMKII Required for Synaptic Plasticity Revealed by Optogenetic Kinase Inhibitor. Neuron 94, 37–47 e35, (2017).

46 Kool, M. J. et al. CAMK2-Dependent Signaling in Neurons Is Essential for Survival. J Neurosci 39, 5424–5439, (2019).

47 Colbran, R. J. Inactivation of Ca2+/calmodulin-dependent protein kinase II by basal autophosphorylation. J Biol Chem 268, 7163–7170, (1993).

48 Takao, K. et al. Visualization of synaptic Ca2+ /calmodulin-dependent protein kinase II activity in living neurons. J Neurosci 25, 3107–3112, (2005).

49 Bhattacharyya, M. et al. Flexible linkers in CaMKII control the balance between activating and inhibitory autophosphorylation. Elife 9, (2020).

50 Wang, Z., Palmer, G. & Griffith, L. C. Regulation of Drosophila Ca2+/calmodulin-dependent protein kinase II by autophosphorylation analyzed by site-directed mutagenesis. J Neurochem 71, 378–387, (1998).

51 Lu, C. S., Hodge, J. J., Mehren, J., Sun, X. X. & Griffith, L. C. Regulation of the Ca2+/CaM-responsive pool of CaMKII by scaffold-dependent autophosphorylation. Neuron 40, 1185–1197, (2003).

52 Hanson, P. I. & Schulman, H. Inhibitory autophosphorylation of multifunctional Ca2+/calmodulin-dependent protein kinase analyzed by site-directed mutagenesis. J Biol Chem 267, 17216–17224, (1992).

53 Wayman, G. A., Lee, Y. S., Tokumitsu, H., Silva, A. J. & Soderling, T. R. Calmodulin-kinases: modulators of neuronal development and plasticity. Neuron 59, 914–931, (2008).

54 Okamoto, K., Narayanan, R., Lee, S. H., Murata, K. & Hayashi, Y. The role of CaMKII as an F-actin-bundling protein crucial for maintenance of dendritic spine structure. Proc Natl Acad Sci U S A 104, 6418–6423, (2007).

55 Borgesius, N. Z. et al. betaCaMKII plays a nonenzymatic role in hippocampal synaptic plasticity and learning by targeting alphaCaMKII to synapses. J Neurosci 31, 10141–10148, (2011).

56 Bayer, K. U., De Koninck, P., Leonard, A. S., Hell, J. W. & Schulman, H. Interaction with the NMDA receptor locks CaMKII in an active conformation. Nature 411, 801–805, (2001).

57 Strack, S., McNeill, R. B. & Colbran, R. J. Mechanism and regulation of calcium/calmodulin-dependent protein kinase II targeting to the NR2B subunit of the N-methyl-D-aspartate receptor. J Biol Chem 275, 23798–23806, (2000).

58 Honda, T. et al. A single phosphorylation site of SIK3 regulates daily sleep amounts and sleep need in mice. Proc Natl Acad Sci U S A 115, 10458–10463, (2018).

59 Cheng, D. et al. Relative and absolute quantification of postsynaptic density proteome isolated from rat forebrain and cerebellum. Mol Cell Proteomics 5, 1158–1170, (2006).

60 Takeuchi, Y., Yamamoto, H., Fukunaga, K., Miyakawa, T. & Miyamoto, E. Identification of the isoforms of Ca(2+)/Calmodulin-dependent protein kinase II in rat astrocytes and their subcellular localization. J Neurochem 74, 2557–2567, (2000).

61 Jewett, K. A. et al. Tumor necrosis factor enhances the sleep-like state and electrical stimulation induces a wake-like state in co-cultures of neurons and glia. Eur J Neurosci 42, 2078–2090, (2015).

62 Saberi-Moghadam, S., Simi, A., Setareh, H., Mikhail, C. & Tafti, M. In vitro Cortical Network Firing is Homeostatically Regulated: A Model for Sleep Regulation. Sci Rep 8, 6297, (2018).

63 Valk, E. et al. Multistep phosphorylation systems: tunable components of biological signaling circuits. Mol Biol Cell 25, 3456–3460, (2014).

64 Wright, P. E. & Dyson, H. J. Intrinsically disordered proteins in cellular signalling and regulation. Nat Rev Mol Cell Biol 16, 18–29, (2015).

65 Karandur, D. et al. Breakage of the oligomeric CaMKII hub by the regulatory segment of the kinase. Elife 9, (2020).

66 Cook, S. G., Buonarati, O. R., Coultrap, S. J. & Bayer, K. U. CaMKII holoenzyme mechanisms that govern the LTP versus LTD decision. Sci Adv 7, (2021).

67 Gangopadhyay, S. S., Gallant, C., Sundberg, E. J., Lane, W. S. & Morgan, K. G. Regulation of Ca2+/calmodulin kinase II by a small C-terminal domain phosphatase. Biochem J 412, 507–516, (2008).

68 Yilmaz, M., Gangopadhyay, S. S., Leavis, P., Grabarek, Z. & Morgan, K. G. Phosphorylation at Ser(2)(6) in the ATP-binding site of Ca(2)(+)/calmodulin-dependent kinase II as a mechanism for switching off the kinase activity. Biosci Rep 33, (2013).

69 Fustin, J. M. et al. Two Ck1delta transcripts regulated by m6A methylation code for two antagonistic kinases in the control of the circadian clock. Proc Natl Acad Sci U S A 115, 5980–5985, (2018).

70 Narasimamurthy, R. et al. CK1delta/epsilon protein kinase primes the PER2 circadian phosphoswitch. Proc Natl Acad Sci U S A 115, 5986–5991, (2018).

71 Ukai, H. et al. Melanopsin-dependent photo-perturbation reveals desynchronization underlying the singularity of mammalian circadian clocks. Nat Cell Biol 9, 1327–1334, (2007).

72 Thiel, G., Greengard, P. & Sudhof, T. C. Characterization of tissue-specific transcription by the human synapsin I gene promoter. Proc Natl Acad Sci U S A 88, 3431–3435, (1991).

73 Blichenberg, A. et al. Identification of a cis-acting dendritic targeting element in MAP2 mRNAs. J Neurosci 19, 8818–8829, (1999).

74 Masuda, T., Tomita, M. & Ishihama, Y. Phase transfer surfactant-aided trypsin digestion for membrane proteome analysis. J Proteome Res 7, 731–740, (2008).

75 Boersema, P. J., Raijmakers, R., Lemeer, S., Mohammed, S. & Heck, A. J. Multiplex peptide stable isotope dimethyl labeling for quantitative proteomics. Nat Protoc 4, 484–494, (2009).

76 Matsumoto, K. et al. Advanced CUBIC tissue clearing for whole-organ cell profiling. Nat Protoc 14, 3506–3537, (2019).

77 Rappsilber, J., Mann, M. & Ishihama, Y. Protocol for micro-purification, enrichment, pre-fractionation and storage of peptides for proteomics using StageTips. Nat Protoc 2, 1896–1906, (2007).

78 Tsujino, K. et al. Establishment of TSH beta real-time monitoring system in mammalian photoperiodism. Genes Cells 18, 575–588, (2013).

79 Mikuni, T., Nishiyama, J., Sun, Y., Kamasawa, N. & Yasuda, R. High-Throughput, High-Resolution Mapping of Protein Localization in Mammalian Brain by In Vivo Genome Editing. Cell 165, 1803–1817, (2016).

80 Sunagawa, G. A., Sei, H., Shimba, S., Urade, Y. & Ueda, H. R. FASTER: an unsupervised fully automated sleep staging method for mice. Genes Cells 18, 502–518, (2013).

81 Niwa, Y. et al. Muscarinic Acetylcholine Receptors Chrm1 and Chrm3 Are Essential for REM Sleep. Cell Rep 24, 2231–2247 e2237, (2018).

82 Ukai, H., Kiyonari, H. & Ueda, H. R. Production of knock-in mice in a single generation from embryonic stem cells. Nat Protoc 12, 2513–2530, (2017).

83 Ode, K. L. et al. Knockout-Rescue Embryonic Stem Cell-Derived Mouse Reveals Circadian-Period Control by Quality and Quantity of CRY1. Mol Cell 65, 176–190, (2017).

84 Kiyonari, H., Kaneko, M., Abe, S. & Aizawa, S. Three inhibitors of FGF receptor, ERK, and GSK3 establishes germline-competent embryonic stem cells of C57BL/6N mouse strain with high efficiency and stability. Genesis 48, 317–327, (2010).

85 Vong, L. et al. Leptin action on GABAergic neurons prevents obesity and reduces inhibitory tone to POMC neurons. Neuron 71, 142–154, (2011).

86 Taniguchi, H. et al. A resource of Cre driver lines for genetic targeting of GABAergic neurons in cerebral cortex. Neuron 71, 995–1013, (2011).

87 Sato, H., Amagai, K., Shimizukawa, R. & Tamai, Y. Stable generation of serum- and feeder-free embryonic stem cell-derived mice with full germline-competency by using a GSK3 specific inhibitor. Genesis 47, 414–422, (2009).

88 Challis, R. C. et al. Systemic AAV vectors for widespread and targeted gene delivery in rodents. Nat Protoc 14, 379–414, (2019).

89 Deverman, B. E. et al. Cre-dependent selection yields AAV variants for widespread gene transfer to the adult brain. Nat Biotechnol 34, 204–209, (2016).

90 Liguore, W. A. et al. AAV-PHP.B Administration Results in a Differential Pattern of CNS Biodistribution in Non-human Primates Compared with Mice. Mol Ther 27, 2018–2037, (2019).

